# The Discovery of Small Molecule Inhibitors of cFLIP that Sensitise Tumour Cells to TRAIL

**DOI:** 10.1101/2024.04.22.590574

**Authors:** Gilda Giancotti, Rhiannon French, Olivia Hayward, Kok Yung Lee, Timothy Robinson, Andreia M. Ribeiro da Silva, Athina Varnava, Marion MacFarlane, Richard W.E. Clarkson, Andrew D. Westwell, Andrea Brancale

**Author notes:** These authors contributed equally to this study (Gilda Giancotti, Rhiannon French, Olivia Hayward). Corresponding authors; Professor Andrea Brancale, PhD, University of Chemistry and Technology, Prague (UCT Prague), Department of Organic Chemistry, Technicka 5, 16628 Prague Czech Republic; Professor Richard Clarkson PhD, Director, European Cancer Stem Cell Research Institute, Cardiff University, School of Biosciences, Hadyn Ellis Building, Maindy Road, Cardiff, CF24 4HQ, Wales, U.K.; Professor Andrea Brancale, PhD, University of Chemistry and Technology in Prague (UCT Prague).

## Abstract

The TNF-related apoptosis-inducing ligand (TRAIL) has potential as a therapeutic agent as it has previously been shown to induce apoptosis in triple-negative breast cancer. Recombinant human TRAIL has shown promise in pre-clinical studies of breast cancer. TRAIL exhibits specificity for triple-negative and treatment-resistant disease subsets. However, several studies have demonstrated that patient tumours exhibit resistance to TRAIL and TRAIL-receptor agonists. We have previously demonstrated that suppression of the TRAIL-receptor inhibitor cFLIP can sensitise breast cancer stem cells to apoptosis inducers, but development of pharmacological inhibitors of cFLIP have been impeded by concerns over structural similarities between cFLIP and the pro-apoptotic procaspase-8.

We used molecular dynamics to model the interactions between cFLIP, procaspase-8 and the TRAIL-receptor Death Inducing Signalling Complex (TRAIL-DISC), followed by virtual pharmacophore screening and in-cell viability assays to identify a small-molecule (OH14, **3**) that selectively inhibited cFLIP binding to the DISC and promoted TRAIL-mediated apoptosis in breast cancer cell lines. When used in combination with TRAIL, OH14 significantly impaired breast cancer cell viability in primary derived and established cell culture.

Given the relatively low (micromolar) potency of the initial hit compound inhibitor OH14 (**3**), limiting its utility as a preclinical development candidate, we carried out structure-activity relationship studies to find a cFLIP inhibitor with more potent cellular activity. Our findings confirm the proof-of-principle that selective pharmacological inhibition of cFLIP can be used to target a vulnerability in breast cancer cells.

## Introduction

Although the development of targeted therapeutics has improved the survival rates of individuals with breast cancer, there remains a subset of patients with triple-negative and metastatic disease for whom the only option is chemotherapy.^1^ Consequently, limited treatment efficacy remains a significant clinical problem in the management of advanced breast cancer, demonstrating the need to continue to develop targeted strategies.

The extrinsic apoptosis pathway mediated by the TNF-related apoptosis-inducing ligand (TRAIL) receptors has potential as a therapeutic target as it has previously been shown to induce apoptosis in triple-negative breast cancer cell lines with a mesenchymal-like phenotype^7^. TRAIL induces apoptosis by binding to one of two death receptors DR4 (TRAIL-R1) and DR5 (TRAIL-R2), leading to the recruitment of FADD, the formation of the Death-Inducing Signalling Complex (DISC) and activation/cleavage of procaspase-8^5^. Recombinant human TRAIL has shown promise in pre-clinical studies of breast cancer. TRAIL exhibits specificity for triple-negative and treatment-resistant disease subsets, suggesting that it may have efficacy for those patients with more aggressive disease^6,7^. However, a Phase II clinical trial of a TRAIL-receptor agonist Tigatuzumab, used in combination with paclitaxel, showed only modest effects in stage 4 breast cancer patients^8^. This and previous studies in lymphoma, lung and colorectal cancer, demonstrate that patient tumours exhibit resistance to TRAIL and TRAIL-receptor agonists in the clinic.^8–11^

We, and others have shown that TRAIL-resistance in breast cancer cell lines is mediated, at least in part, by the apoptosis inhibitor cFLIP; inhibition of cFLIP by siRNA can sensitise previously resistant breast cancer cell lines and specifically bCSCs to TRAIL^3,12^. Pre-clinical studies have also shown that indirect inhibition of cFLIP via the use of HDAC inhibitors such as vorinostat or droxinostat which reduce cFLIP expression can sensitise breast cancer cell lines to TRAIL both *in vitro* and in xenograft models^13^. However, such non-specific inhibitors have shown toxicity in phase I trials^14, 15^.

The development of a targeted and selective cFLIP inhibitor has previously proven difficult due to homology with the pro-apoptotic procaspase 8. This homology is important for how cFLIP functions. cFLIP and procaspase-8 are both recruited to the DISC via their Death Effector Domain 1 (DED1); procaspase 8 to FADD and cFLIP primarily to procaspase 8, to exert activating and inhibitory functions respectively^16^. However, the specific residues in the DED1 required for these respective interactions have not been fully characterised. In this study we have compared the procaspase 8-FADD and cFLIP-procaspase 8 interactions and have identified targetable differences between the two complexes. Mutagenesis of key residues in cFLIP DED1 impaired the inhibitory function of cFLIP thus confirming the importance of the DED1 domain for incorporation of cFLIP into the DISC complex. These findings led us to develop small molecule inhibitors of cFLIP function, which at micromolar concentrations perturbed the binding of cFLIP to the DISC without affecting the recruitment of procaspase-8 to FADD, thus allowing procaspase 8 cleavage and activation. Our initial hit compound from virtual screening (OH14) sensitised breast cancer cell lines, specifically breast cancer stem cells, to TRAIL *in vitro* leading to a reduction in tumour initiation *in vivo*. We then carried out structure-activity relationship studies based on OH14 to optimise cellular potency and selectivity of our novel cFLIP inhibitors. Our discovery and early optimisation work reported here describes for the first time the proof-of-principle that pharmacological inhibition of c-FLIP impacts on breast cancer cell viability mediated by the extrinsic apoptosis pathway.

## Results

### Molecular modelling of procaspase8:cFLIP interaction identifies a binding pocket which differs from that of FADD:procaspase 8

The schematic shown in Figure 1A summarises a current model of DISC protein interactions. TRAIL binding induces receptor trimerization and the recruitment of FADD, via death domain (DD) interactions between FADD and DR4 or DR5 receptors. FADD in turn induces procaspase-8/10 recruitment via death effector domain (DED) interactions between FADD and the DED1 of procaspase-8^5,16^. It was previously thought that cFLIP primarily competes with procaspase-8 by binding directly to FADD to block downstream signalling^17^. However, analysis of DISC formation suggests that the complex consists of a ‘chain’ of procaspase-8 molecules binding to each other via their DED1 pocket and exposed DED2 FL motif ^5,16,18^. cFLIP, which occurs as both long (L) and short (S) isoforms, exerts an inhibitory function by incorporation into these procaspase 8 protein chains and therefore is predominantly but not exclusively recruited to FADD indirectly via procaspase 8^16^ (Figure 1A). Recombinant cFLIP has also been shown to bind directly to FADD via DED2 however this was observed in the absence of caspase 8, and so its physiological relevance remains to be confirmed^19^. cFLIP and procaspase 8 therefore exert opposing functions but both are mediated by binding via DED1. We set out to model these interactions in order to identify targetable differences unique to cFLIP.

**Figure 1:**
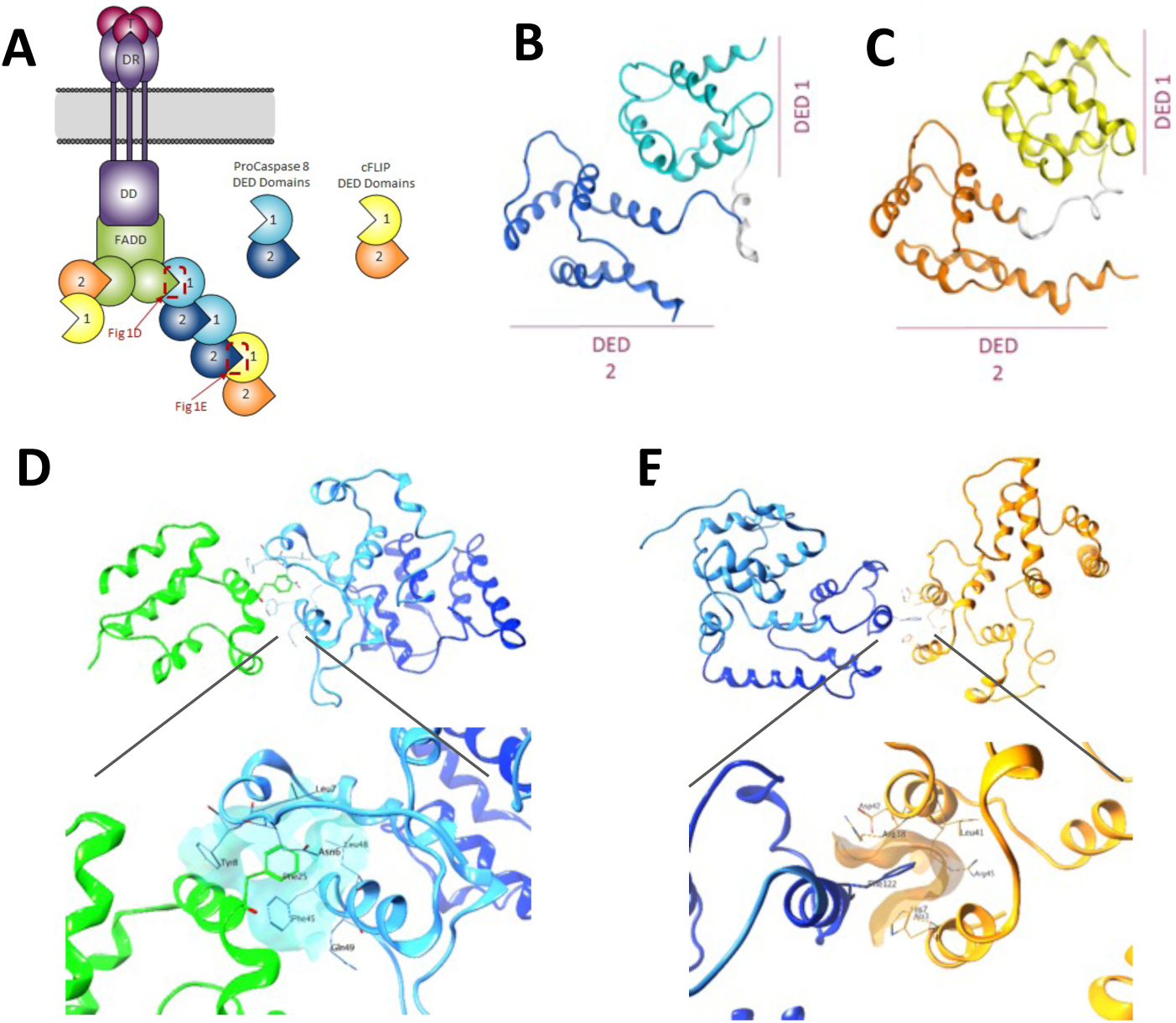
Molecular Modelling identifies a binding pocket of cFLIP DED1:procaspase 8 DED2 interaction which differs from procaspase 8 DED1:FADD DED2 interaction. **A:** Schematic of protein binding at DISC: Upon activation of receptor, Procaspase 8 binds to FADD via DED1. Recruited cFLIPL or S binds to Procaspase 8 also via DED 1. See text for further details. **B/C:** 3D structure of the two DEDs of (**B**) procaspase 8 and (**C**) cFLIP modelled *in silico* using a sequence homology model with viral FLIP (MC159). **D:** Interaction between procaspase-8 DED1 pocket and FL motif on DED2 of FADD including a close-up detailing the FADD:procaspase-8 binding pocket identifying key procaspase 8 residues. **E:** Interaction between c-FLIP DED1 and procaspase 8 DED2 including a close-up detailing the cFLIP:procaspase 8 binding pocket identifying key cFLIP residues. Key: FADD = green, Procaspase 8 = blue, cFLIP = yellow/orange.

To compare the structures of cFLIP and procaspase 8, we modelled both proteins and their interactions *in silico.* At the time we performed our *in silico* calculation, no crystal structures of cFLIP and procaspase-8 DEDs were yet available. We therefore constructed them based on a comparative homology model using MOE. The protein MC159 (PDB ID: 2BBR), which has been used previously for modelling DED interactions ^5,16,20,21^ was identified by sequence comparison (**Supplementary Figure 1A**) as the best candidate for the construction of cFLIP and procaspase-8 DED homology models (**Figure 1B and C**). Recently, the structure of FADD:caspase-8 has been resolved, and our model proved to be comparable to the experimental structure (**Supplementary Figure 1B-C**).^22,23^ Model comparison revealed only 22% structural homology between cFLIP and procaspase-8. Comparison of procaspase8:FADD and the cFLIP:procaspase8 models revealed that these interactions also involved different residues (Figure 1D and E). Unique residues were identified in the cFLIP DED1 pocket as potentially important for the interaction with procaspase 8, these being H7, K18, R38 and R45 (**Figure 2A**).

**Figure 2:**
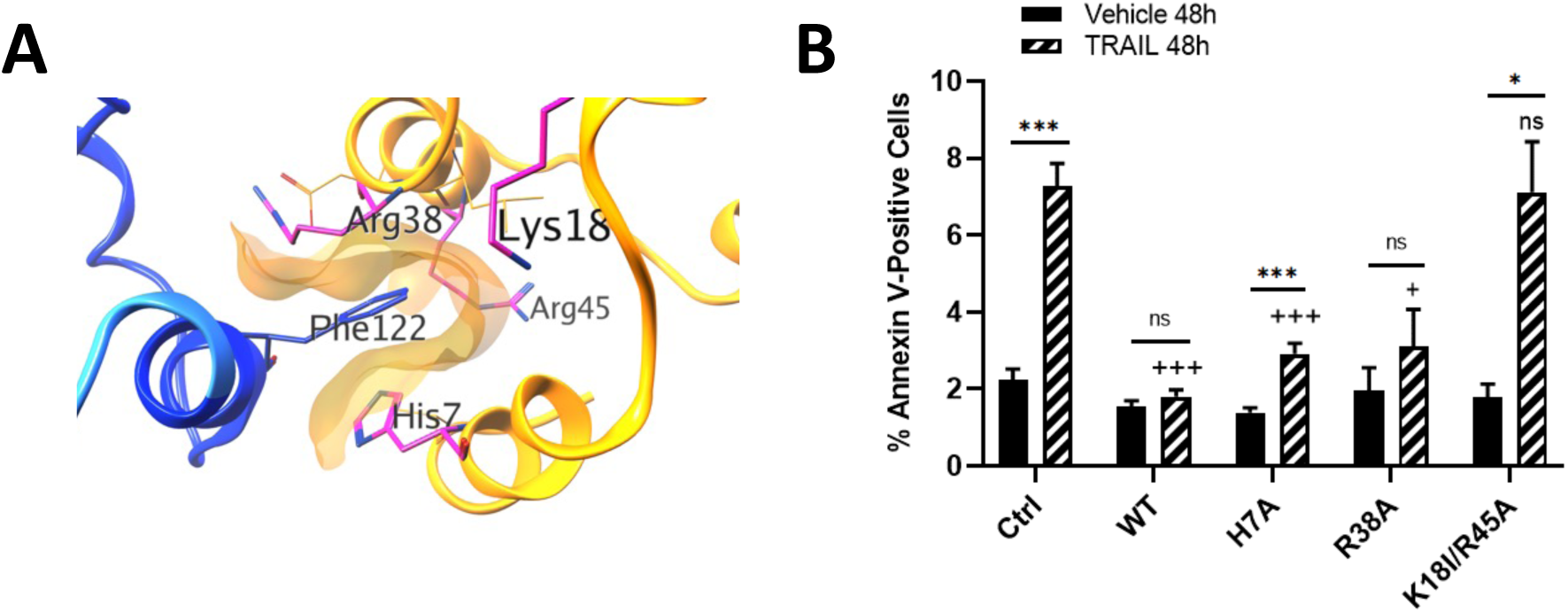
Mutagenesis of key residues in DED1 pocket of cFLIP influences TRAIL mediated cell death. **A:** Model of the positioning of the residues targeted for mutation within the DED1 pocket of cFLIP in relation to a key interactive residue Phe122 within procaspase 8. Key: Procaspase 8 = yellow/orange, cFLIP = blue. **B:** HeLa cells stably transfected with WT or mutant cFLIPL (**Supplementary Figure 2A**) were treated with TRAIL for 48 hours and cell death determined by annexin V staining using Incucyte real-time analysis. Cell death is expressed as the percentage of annexin positive cells following treatment and error bars represent standard error of the mean from six independent experiments, determined by empirical observation for comparison between TRAIL treated control and mutant samples. * = p<0.05, ** = p<0.01 between vehicle and TRAIL treated for each construct; + = p<0.05, ++ = p<0.01 between Ctrl TRAIL and mutant construct TRAIL. Equivalent analysis of cell death at 72 hours of TRAIL treatment was also measured (**Supplementary Figure 2B**).

### Mutagenesis of key residues confirms relevant interaction

To test the predictions of our models and determine whether these four unique residues in the cFLIP DED1 pocket were critical for cFLIP:procaspase 8 binding, we generated cFLIP expression constructs harbouring the single mutations (H7A or R38A) or the double mutation (K18I and R45A). HeLa cells overexpressing wild-type or mutant cFLIP constructs were first analysed for total cFLIP expression levels to allow for subsequent comparison. Each sub-line exhibited approximately 50 (K18I/R45A) to 75-fold (R38A and H7A) overexpression with no statistical significance between cFLIP levels of wild-type and mutant lines (**Supplementary Figure 2A**). We then assessed the ability of each construct to protect cells from TRAIL-induced apoptosis. Overexpression of wild-type cFLIP elicited near total protection of HeLa cells from TRAIL after 48 hours (**Figure 2B**) or 72 hours (**Supplementary Figure 2B**) treatment, consistent with previous reports of the effect of cFLIP upregulation on DISC-mediated apoptosis^3,4,12^. In contrast, overexpression of the H7A or R38A cFLIP mutants only partially (but still significantly) protected from TRAIL-induced apoptosis, while the K18I/R45A double mutant completely impaired cFLIP-mediated protection to the extent that TRAIL-induced cell death was no longer statistically different from the empty vector control (Figure 2B). This indicates that while both H7 and R38 have a minor contribution to the inhibitory function of cFLIP, one or both K18 and R45 residues in the cFLIP binding pocket are particularly important for cFLIP activity.

### An in-silico screen identifies a small molecule that suppresses TRAIL resistance

Having supported our cFLIP model by mutagenesis we used it to virtually screen a library of over 350,000 commercially available compounds (SPECS library) with the aim of identifying potential inhibitors specific to the cFLIP DED1 binding pocket. Initially, the library was filtered through a pharmacophore query designed from the interaction contacts between cFLIP DED1 and FADD. Approximately 14,000 virtual ‘hits’ were identified, and these were docked and scored based on their ability to bind to the c-FLIP pocket, using GLIDE. Subsequently, the top 2,000 compounds were analysed for their ability to disrupt the procaspase8_DED1:FADD interaction *in silico,* and potential inhibitors of this interaction were discounted. Finally, following a visual inspection of the resulting hits within pockets of c-FLIP and procaspase-8, 19 compounds were then selected for biological analysis based on specificity for the cFLIP DED1 pocket. These 19 compounds were analysed for their ability to sensitise the TRAIL-resistant breast cancer cell lines^3,4,7^ MCF-7 (ER+ve) and BT474 (ER/HER2+ve) to TRAIL *in vitro* (**Supplementary Figures 3-5**). Using this screen, we identified one compound, known as OH14 (**3**), which sensitized MCF-7 cells to TRAIL to a similar extent as cFLIP suppression by siRNA. OH14 (**3**), synthesised from 2,4-dichloro-5-methyl-benzenesulfonyl chloride (**1**) and anthranilic acid (**2**) (**Scheme 1**), was then selected for further testing of its capacity to specifically promote TRAIL-DISC mediated apoptosis and to target CSC viability.

**Scheme 1:.**
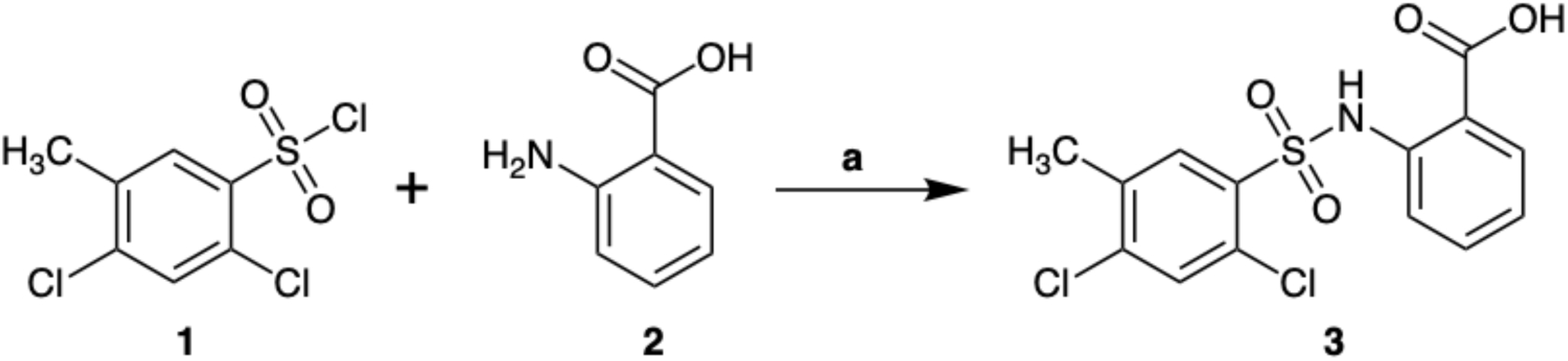
Reagents and conditions: NaoH, H_2_O, 70°C, 18h

### OH14 (**3**) promotes TRAIL sensitivity via cFLIP

Firstly, the within-cell potency of OH14 (**3**) was determined in both HeLa cells and the breast cancer cell line, MCF7, using different apoptosis assays. Cells were treated with a range of OH14 concentrations for 1 h prior to 20 ng/ml TRAIL for 24 hours. A dose-dependent sensitization effect was observed with a significant increase in cleaved caspase 3/7 (**Figure 3A**) and Annexin V (**Figure 3B**) staining seen at concentrations of 75 μM and 100 μM OH14.

**Figure 3:**
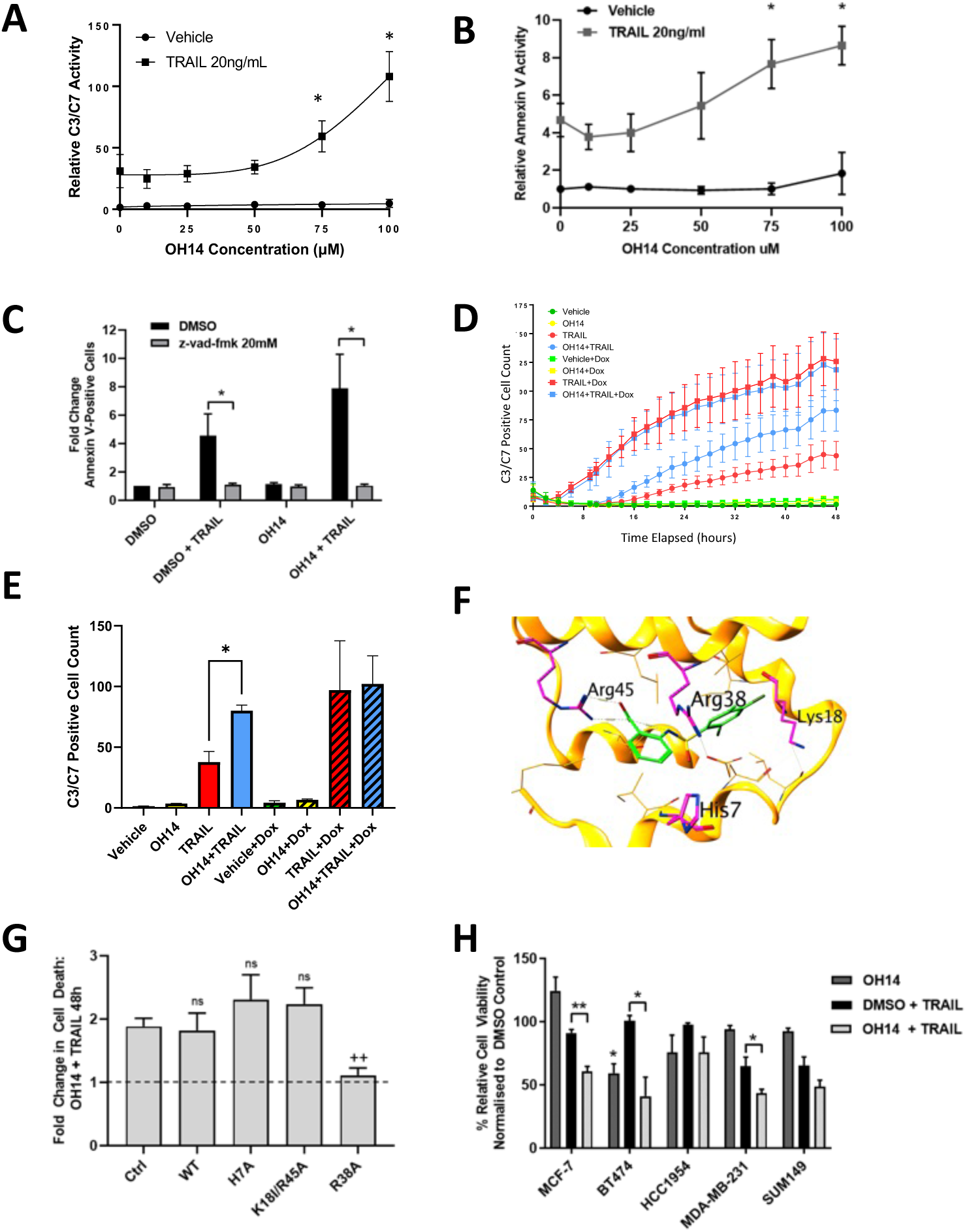
OH14 (**3**) Sensitises cancer cells to TRAIL via cFLIP. **A and B:** MCF7 cells (**A**) and HeLa cells (**B**) were treated with a range of concentrations of OH14 followed by 20ng/ml TRAIL for 48 hours and viability assessed by cleaved caspase 3/7 (**A**) and Annexin V (**B**) staining using Incucyte® real-time analysis. * = p<0.05 vs untreated control, n=3 determined empirically by significance at maximum dose. **C:** HeLa cells were treated with 100 μM OH14 in the presence of 20 mM Z-Vad-fmk pan-caspase inhibitor followed by 20ng/ml TRAIL for 18 hours and viability assessed by Annexin V staining using Incucyte real-time analysis. * = p<0.05 vs untreated control, n=3 determined empirically by significance of DMSO plus TRAIL. **D and E:** MCF7 cells stably overexpressing a doxycycline inducible shRNA targeting cFLIP were treated with doxycycline and/or 20ng/ml TRAIL and/or 75 μM OH14 for up to 48 hours and monitored for cleaved caspase 3/7 by Incucyte real-time analysis. Graph (**D**) represents one of three independent repeats and bar chart (**E**) represents mean of 3 independent experiments at 48 hours, determined empirically by significance of TRAIL plus OH14 in the absence of Dox. * = p<0.05 vs TRAIL alone. **F:** Model of OH14 binding to DED1 pocket of cFLIP highlights the proximity of the interaction between OH14 (green) with Arg38. **G:** HeLa cells stably transfected with WT or mutant cFLIPL constructs were treated with 100 μM OH14 followed by 20 ng/ml TRAIL for 18 hours and viability assessed by Annexin V staining using Incucyte® real-time analysis. Control is empty vector transfected cells. Data presented as fold change in cell death compared to TRAIL-treated cells for each construct. N=3 *= p<0.05 compared to control (for absolute values of % annexin V positive cells, see Supplementary Figure 8). **H**. A panel of breast cancer cell lines were treated with 100 μM OH14 followed by 20 ng/ml TRAIL for 18 hours and viability assessed by CellTitre-Blue assay. N=3 * = p<0.05, ** = p<0.01 compared to TRAIL alone.

Despite the requirement for relatively high concentrations of OH14 to sensitize to TRAIL, the effects of OH14 were both cFLIP dependent and mediated exclusively by caspase activation. Thus, inhibition of caspase activity by the pan-caspase inhibitor z-vad-fmk completely rescued the TRAIL-induced apoptosis observed in the presence of OH14 (**Figure 3C**). Moreover, conditional suppression of cFLIP expression through doxycycline inducible shRNA targeting of cFLIP (**Supplementary Figure 7**), abrogated the sensitizing effects of OH14 to TRAIL for up to 48 hours in real time analysis of apoptosis (**Figure 3D and 3E**). This confirmed that the sensitizing effects of OH14 were dependent upon the presence of cFLIP.

Furthermore, we confirmed the selectivity of OH14 for the cFLIP DED1 binding site pocket (**Figure 3F**) using the wild-type and mutant cFLIP constructs described above (**Figure 2B and 3G**). Overexpression of wild type cFLIP did not impair the ability of OH14 to sensitize to TRAIL, confirming that OH14 maintains its ability to sensitize to TRAIL at this level of cFLIP overexpression (**Figure 3G**). As predicted from the pharmacophore model, overexpression of the K18I/R45A double-mutant cFLIP had no significant effect on OH14 function compared to empty vector control (**Figure 3F** and **3G**) as this mutant cFLIP molecule was functionally inert (**Figure 2B**). OH14 also efficiently induced TRAIL sensitization in the H7A cFLIP mutant line, indicating that OH14 was still able to bind and de-repress this partially inhibitory mutant. However, OH14 was unable to re-sensitize cells overexpressing the cFLIP mutant R38A to TRAIL, demonstrating the importance of the R38 residue in cFLIP for OH14 function and thus providing additional evidence for its selective action on the cFLIP DED1 pocket (**Figure 3G** and **Supplementary Figure 8**).

Finally, we tested the cell-type specificity of OH14 in a panel of breast cancer cell lines. OH14 significantly sensitized MCF-7 (ER+ve), BT474 (HER2+ve) and MDA-MB-231 (triple-negative) cell lines to TRAIL (**Figure 3H**), with similar but statistically insignificant trends also observed in HCC1954 (HER2+ve) and SUM149 (triple-negative) breast cancer cells.

### OH14 (**3**) inhibits binding of cFLIP to the DISC components

Having established the specificity of OH14 (**3**) for cFLIP mediated cell sensitization we next set out to confirm the effect of OH14 on disrupting cFLIP interactions with DISC proteins. Initially, recruitment of cFLIP to the DISC *in vitro* was demonstrated by FRET analysis using FADD-CFP and cFLIP-YFP FRET constructs expressed in HeLa cells. A significant increase in FRET fluorescence intensity was observed following 2 hours pre-treatment with TRAIL, indicating a TRAIL-dependent recruitment of cFLIP-YFP and FADD-CFP in intact cells in culture (**Figure 4A**). This FRET interaction was abrogated when OH14 was co-cultured with TRAIL under the same conditions, suggesting that the TRAIL-mediated proximal recruitment of cFLIP with FADD was disrupted by OH14.

**Figure 4:**
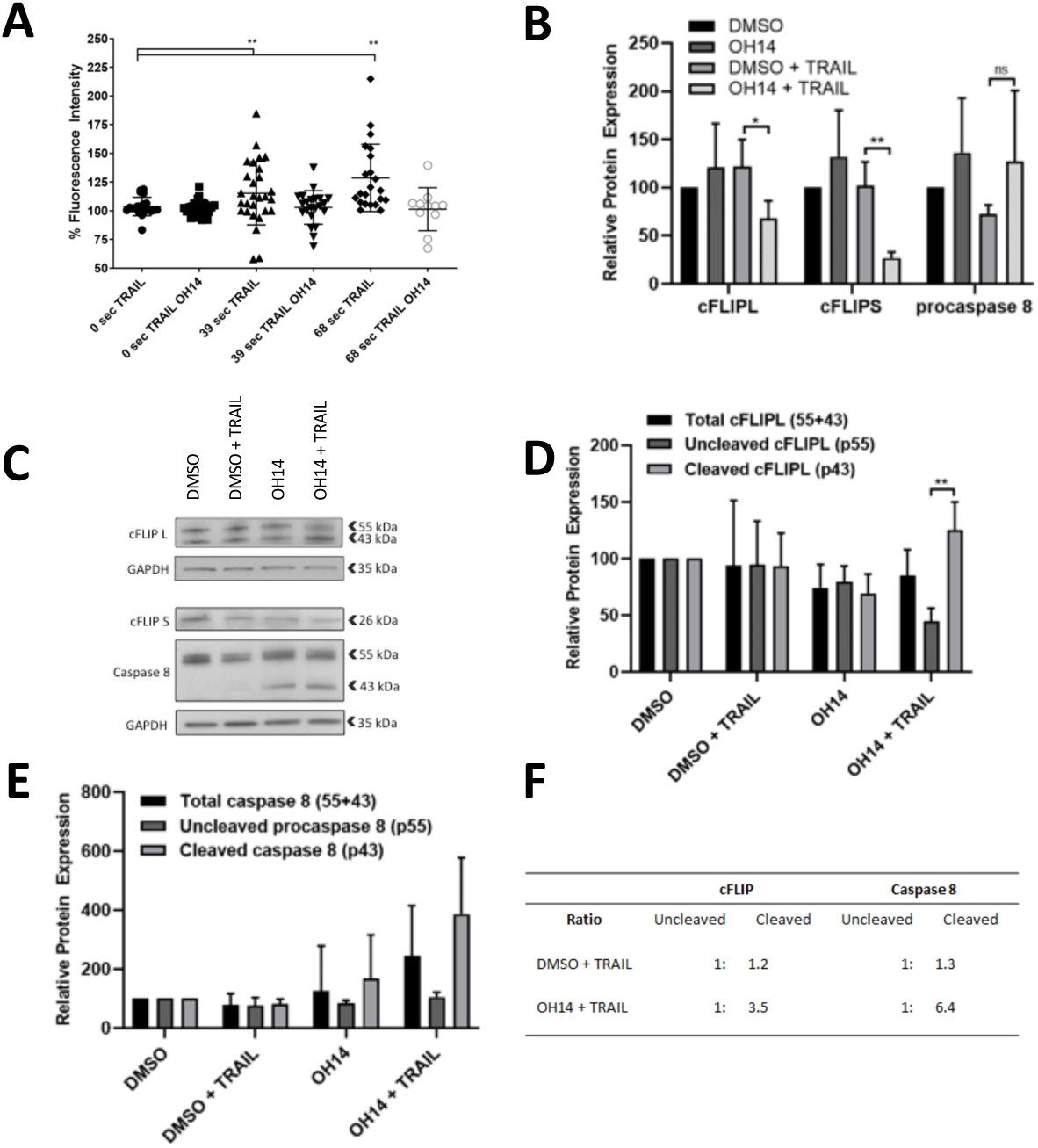
OH14 impairs cFLIP recruitment to FADD complex and promotes cFLIP and caspase 8 cleavage. **A:** HeLa cells were treated with the pan-caspase inhibitor Z-VAD-FMK 1 hour before transfection with cFLIP-YFP or FADD-CFP FRET constructs for 24 hours. Cells were then treated with 100 µM OH14 for 1 hour followed by 2 hours 20 ng/ml TRAIL and fluorescence was measured at 0-, 39-, and 68-seconds following photon-bleaching. Each data point represents a recording made on an individual cell. N=20-28 per timepoint**. B:** Densitometry of western blot analysis of FADD immunoprecipitates in MCF-7 cells pre-treated with 100 μM OH14 for 1 hour followed by 2 hours with 20 ng/ml TRAIL. Average of four independent experiments empirically determined by significance of TRAIL vs OH14+TRAIL (**Supplementary Figure 9**) normalised to DMSO control. ** = p<0.01 vs TRAIL alone, *** = p<0.001 vs TRAIL alone. **C:** Western blot analysis of total cell lysates of MCF-7 cells pre-treated with 100 μM OH14 for 1 hour followed by 2h 20 ng/ml TRAIL. **D/E:** Densitometry of three independent replicates of the western blot analysis shown in **C**, highlighting cFLIP cleavage products (**D**) and procaspase 8 cleavage products (**E**). **F:** Ratio of cleaved to uncleaved proteins measured in D and E. **p < 0.01.

We also assessed the ability of OH14 to disrupt this interaction in MCF7 cells by co-immunoprecipitation. MCF7 cells were pre-treated with TRAIL and/or OH14 followed by immunoprecipitation of FADD from cell lysates. While only semi-quantitative at best, significant differences between means from four independent experiments confirmed that OH14 disrupted the TRAIL-induced interaction between FADD and both short and long forms of cFLIP but did not inhibit the interaction between FADD and procaspase 8 (**Figure 4B** and **Supplementary Figure 9**). In the absence of TRAIL however, OH14 had no effect on FADD:cFLIP interactions. This is consistent with the chain elongation model of cFLIP recruitment to the DISC summarised in **Figure 1**, whereby in the uninduced state cFLIP likely interacts with FADD via its DED2 domain, while TRAIL-induced recruitment of procaspase 8 to the DISC allows cFLIP to be recruited via its DED1 domain. This supports the model of OH14 involvement in DED1 interactions.

OH14 had no significant effect on total cFLIP levels, either with or without TRAIL stimulation, which suggests that the disruption of TRAIL mediated recruitment of cFLIP to the DISC described above is not due to destabilisation of total cFLIP protein levels at this dose and duration of treatment (**Figure 4C** and **4D**). However, the ratio of noncleaved cFLIP (55kDa) to the cleaved form (43kDa) was significantly altered by OH14 when cells were stimulated with TRAIL, resulting in an increase in cleaved cFLIP (Figure 4C, 4D and 4F). As cFLIP is cleaved by caspase-8, this result is indicative of caspase activation and consistent with the increase in apoptosis seen in the presence of OH14 (**Figure 3**). Indeed, OH14 treatment induced procaspase 8 cleavage to the active 43kDa form (Figure 4C -4F).

Taken together these results suggest that OH14 disrupts the recruitment of cFLIP to the DISC, allowing for caspase activation and potentiation of the TRAIL signal.

### Synthesis of OH14 (**3**) analogues for SAR studies

Having confirmed the efficacy of the original compound OH14 (**3**) to sensitize cells to TRAIL via disruption of the DISC, we set out to investigate the structure-activity relationships around this chemical scaffold. Initially, given the importance of the sulfonamide linker for the arrangement in the c-FLIP pocket, this functional group was retained. On the other hand, changes on both aromatic rings were contemplated. The original 2,4-dichloro-5-methyl substituent was replaced with numerous hydrophobic groups at different positions of the ring with the aim of investigating whether other hydrophobic substituents may improve penetration and stability of the molecule in the pocket. The replacement of the phenyl ring with different heterocycles and bigger rings such as naphthalene and quinoline was explored. The carboxylic acid group is deemed to be crucial for the activity of **3** because of its proposed ability to interact with Arg45 of c-FLIP, thus preventing the binding between this amino acid and Phe25 of FADD. However, the presence of a carboxylic acid moiety is often associated with limitations such as reduced ability to cross the lipophilic layers of the cellular membrane or potential toxicity derived from metabolism. Replacement of the carboxylic acid group with different alkyl esters and bioisostere groups such as tetrazoles was therefore considered. Carboxamide and N-alkyl-carboxamides were also explored as substituents of the carboxylic acid to investigate how these functional groups may affect activity of the original hit. **Schemes 2-5** summarise all the explored sulfonamide modifications and how they were synthesised. All molecules conserve a central scaffold characterised by the sulfonamide linker which connects the differently substituted hydrophobic ring and the functionalised ring (**supplemental Table 1**).

**Scheme 2:**
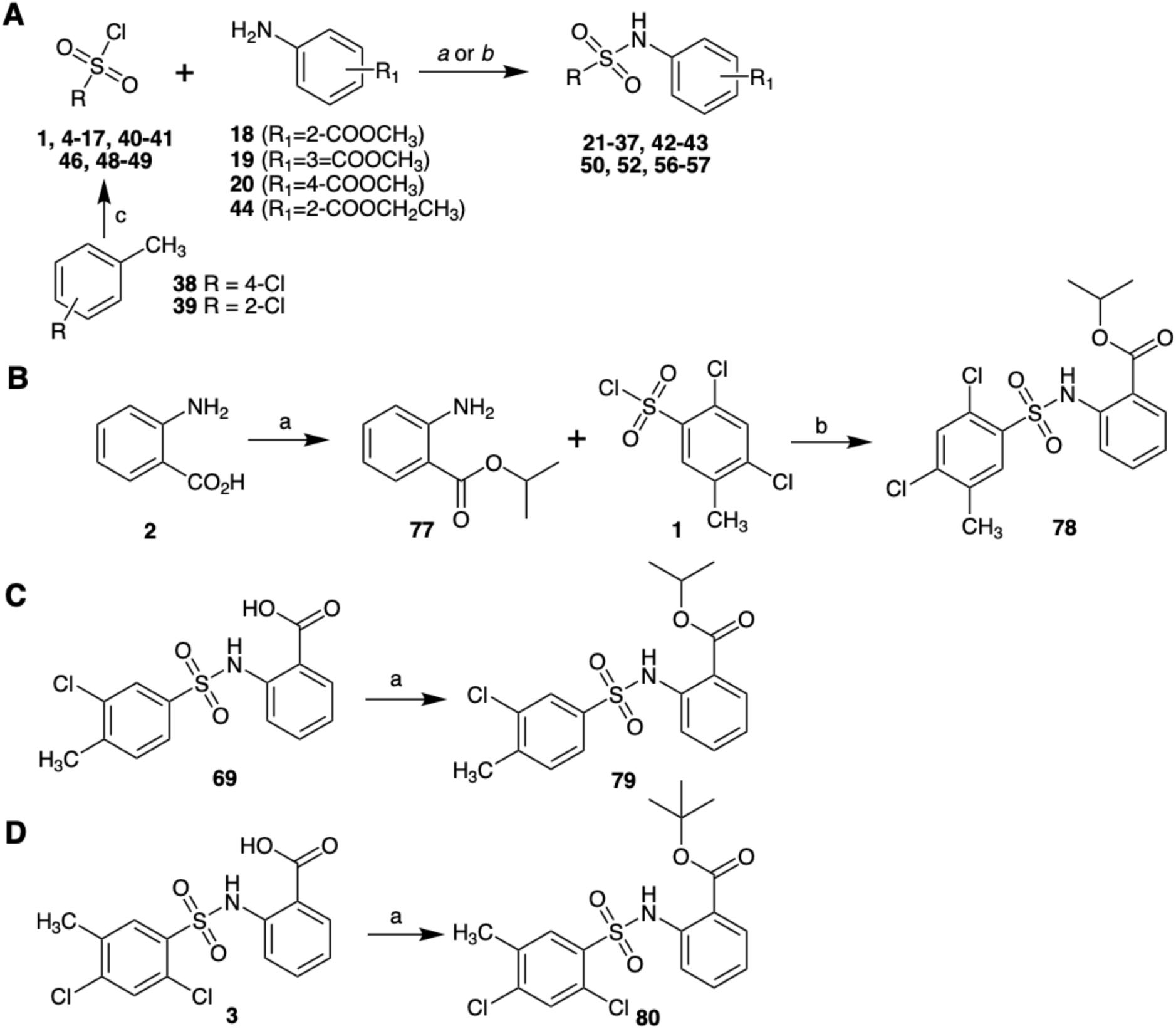
Synthesis of alkyl(arylsulfonamidobenzoates). *Reagents and conditions:* **A:** *a:* pyridine, room temp., 6h; *b:* pyridine, CH2Cl2, 0°C; *c:* chlorosulfonic acid, CHCl3, 0°C to room temp., 45 min. **B:** *a: i-*PrOH, 1.25M HCl, reflux, 48h; *b:* pyridine, room temp., 6h. **C:** *a:* TBTU, DIPEA, DMF, room temp., 30 min., then *i-*PrOH, room temp., 7h. **D:** *a: t-*BuOH, MgSO4, H2SO4, CH2Cl2, room temp., 18h.

To investigate how the linker might affect the activity of the compounds, other functional groups were considered as replacement of the sulfonamide group. Initially an amine group, which retains the original tetrahedral geometry required for the adjustment of the molecules to the curved-shaped cavity of the c-FLIP pocket, as well as the original distance between the two aromatic rings was considered. The designed amine derivatives retain the carboxylic acid function in the *ortho* position of the original anthranilic ring (**Supplementary Table 2**). However, using the same criteria discussed above, the substitution of the carboxylic acid moiety with a methyl ester group, or with the carboxylic acid bioisosteres, tetrazole and oxadiazole rings, was also considered. In addition, modifications designed for the hydrophobic ring were mostly selected according to the early biological data obtained for the sulfonamide derivatives. The replacement of the original phenyl ring with bigger aromatic rings such as naphthalene and quinoline, and with the heterocycle pyridine was also explored.

Finally, we prepared two additional small series of analogues, characterised by the replacement of the original sulfonamide moiety with a methylene and an amide group. The central scaffold of the new series of compounds is represented in **Supplementary Table 3**. The methylene linker shortens the distance between the two aromatic rings, whilst retaining the desired molecular geometry for the correct arrangement of the molecule in the pocket of c-FLIP. On the contrary, the introduction of an amide linker would induce a relatively planar conformation, allowing to investigate how changes in the molecular geometry might affect the activity of the compounds.

### Synthesis of alkyl(arylsulfonamidobenzoates)

The synthesis of ester derivatives of OH14, classified as alkyl(arylsulfonamidobenzoates) is shown in **Scheme 2**. Given the commercial availability of most of the sulfonyl chlorides required for the synthesis of the new analogues, the one step reaction with methyl aminobenzoates **18-20** was carried out to obtain the analogues **21-37** and **42-43**, via nucleophilic attack by the amino group of anthranilic ester on the electrophilic sulfur atom, followed by elimination of chloride. The two starting sulfonyl chlorides **40-41** were obtained by chlorosulfonation between chlorosulfonic acid and the corresponding toluenes **38-39.** Compounds **50,52, 56-57** were obtained via the same mechanism, however reaction conditions were slightly changed from *a* to *b*.

Alternative esters such as the isopropyl ester and the *tert*-butyl ester were considered through the synthesis of compounds **78**, **79** and **80**. These derivatives aimed to explore whether larger alkyl groups, which show higher stability to hydrolysis compared to the methyl ester, may influence the activity of the compounds. To make a direct comparison between the different alkyl esters, the isopropyl derivative and the *tert*-butyl derivative of **21** and **43** were synthesised. The isopropyl 2-aminobenzoate was obtained via esterification of the anthranilic acid **2**. Although the time of the reaction was increased to 48 hours, only a partial conversion of the starting material into the corresponding isopropyl ester was reached, probably due to the low reactivity of the isopropyl alcohol (overall 19% yield). The following step involves the reaction with sulfonyl chloride **1** to give the final product **78** (Scheme 2B). Compound **79** (Scheme 2C), the isopropyl derivative of **43,** was obtained via esterification of the carboxylic acid derivative **69**. Two different synthetic strategies were applied. Here, the coupling agent 2-(1H-benzotriazol-1-yl)-N,N,N’,N’tetramethylaminium tetrafluoroborate (TBTU) was used, leading to the formation of the corresponding isopropyl ester **79**. The *tert-*butyl derivative **80** (Scheme 1D) was synthesised by reacting the carboxylic acid and the tertiary alcohol in the presence of H_2_SO_4_ absorbed on MgSO_4_.

Isolated yields and identities for all the alkyl(arylsulfonamidobenzoates) compounds synthesised are shown in **Supplementary Table 4**.

### Synthesis of (arylsulfonamido)benzoic acids (**3, 64-67, 72-76**)

The reaction schemes to synthesise the (arylsulfonamido)benzoic acids is shown in **Scheme 3**. Analogues **3** (OH14) and **64-67** were synthesised by reacting anthranilic acid **2** with the corresponding arylsulfonyl chlorides (**1**, **59-62)** under basic conditions, whereas derivatives **72-76** were obtained by basic hydrolysis of the corresponding methyl esters **34-37, 57**.

**Scheme 3:**
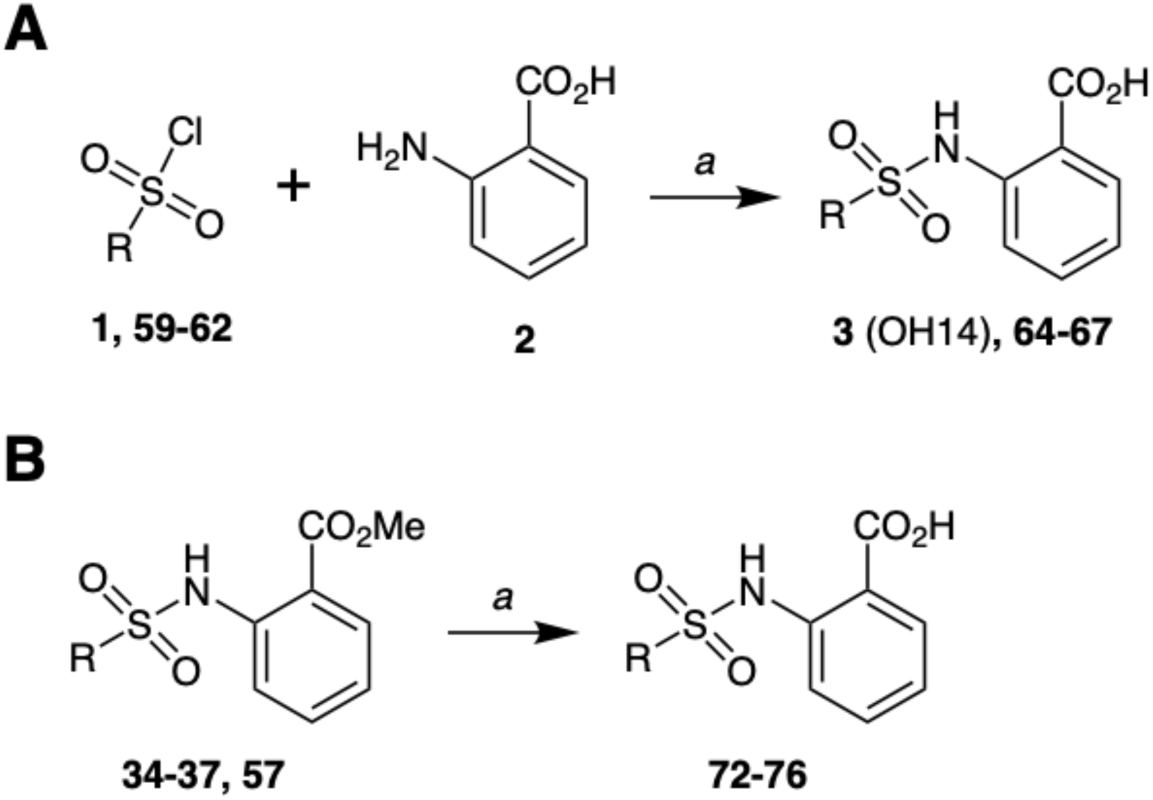
Synthesis of (arylsulfonamido)benzoic acids. *Reagents and conditions:* **A:** *a:* NaOH, H2O, 70°C, 16h. **B:** *a:* LiOH, THF, MeOH, H2O, 70°C, 16h.

### Synthesis of arylsulfonamido-N-(1H-tetrazol-5-yl)benzene derivatives (**88-93**)

Analogues **88-93** feature a tetrazole group as carboxylic bioisosteres, providing the possibility of better membrane permeability through increased lipophilicity, plus improved metabolic stability and reduced cytotoxicity. The tetrazole intermediate **84** was synthesised via a 1,3-dipolar cycloaddition between *ortho-*aminobenzonitrile **82** and sodium azide, followed by the reaction of **84** with arulsulfonyl chlorides in pyridine (**Scheme 4**).

**Scheme 4:**
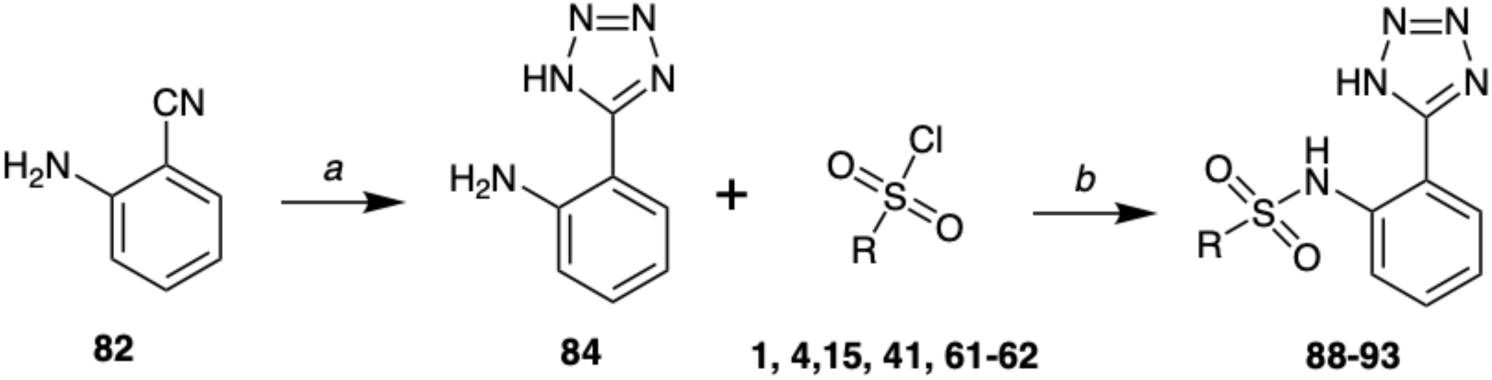
Synthesis of arylsulfonamido-N-(1H-tetrazol-5-yl)benzene derivatives. *Reagents and conditions: **a**:* NaN3, NH4Cl, DMF, 120°C, 72h. **b**: pyridine, room temp., 6h.

### Synthesis of alkyl(arylsulfonamido)benzamides (**101-106**)

Analogues **101**-**106** are characterised by the replacement of the methyl ester of compounds **21** and **43** with differently substituted carboxamides to increase the hydrolytic stability of these molecules. The synthetic strategy applied to obtain the carboxamide derivatives involves direct amidation of the carboxylic acids using the coupling reagent N,N’-carbonyldiimidazole (CDI) (**Scheme 5A**). To obtain the primary carboxamide derivative of **3** the conditions of the reaction were slightly changed. The organic solvent was replaced by the 1-butyl-3-methylimidazolium tetrafluoroborate ([bmim]BF4) ionic liquid to promote the solubility of the ammonium acetate which was used as source of ammonia (**Scheme 5B**). Product **106** was obtained with an isolated yield of 10%.

**Scheme 5:**
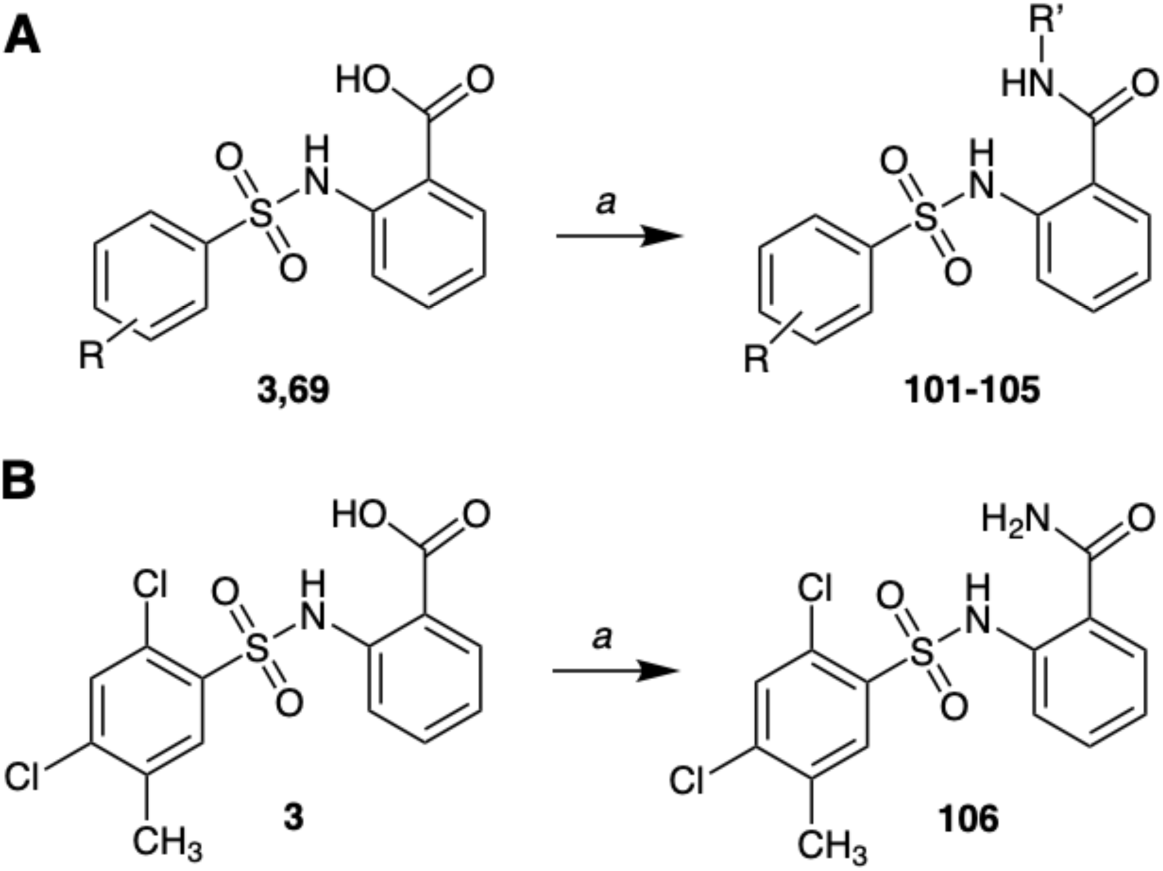
Synthesis of alkyl(arylsulfonamido)benzamides. *Reagents and conditions:* **A***: a*: R’-NH2, CDI, THF, room temp., 16h. **B***: a:* CDI, [bmim]BF4, 80°C, 2h; then NH4OAc, NEt3, 80°C, 16h.

### Synthesis of methyl(arylamino)benzoates (**121, 123, 129, 131, 133-134, 137-138**)

Structural variation of the sulfonamide linker that is present in the original hit compound **3** (OH14) was tested through synthesis of a range of amine-linked derivatives, initially preserving the methyl ester group with the aim to increase the lipophilicity of the compounds to enhance their ability to cross the cellular membrane. In derivatives **137-138** the amine linker was elongated by adding an additional carbon atom. The synthetic methods used to obtain the amine-linker methyl ester derivatives are shown in **Scheme 6**. Methyl ester derivatives **121** and **123** were synthesised via esterification of the corresponding carboxylic acids **113** and **115**, using the coupling agent TBTU in the presence of DIPEA and methanol. Derivatives **129, 131, 133-134** were obtained in good yields via reductive amination of the corresponding aldehydes, and derivatives **137-138** were obtained in low isolated yields (due to unreacted starting material) via nucleophilic substitution of substituted bromides using methyl-2-amino benzoate **18**. Isolated yields and identities for all the methyl(arylamino)benzoates synthesised are shown in **Supplementary Table 5**.

**Scheme 6:**
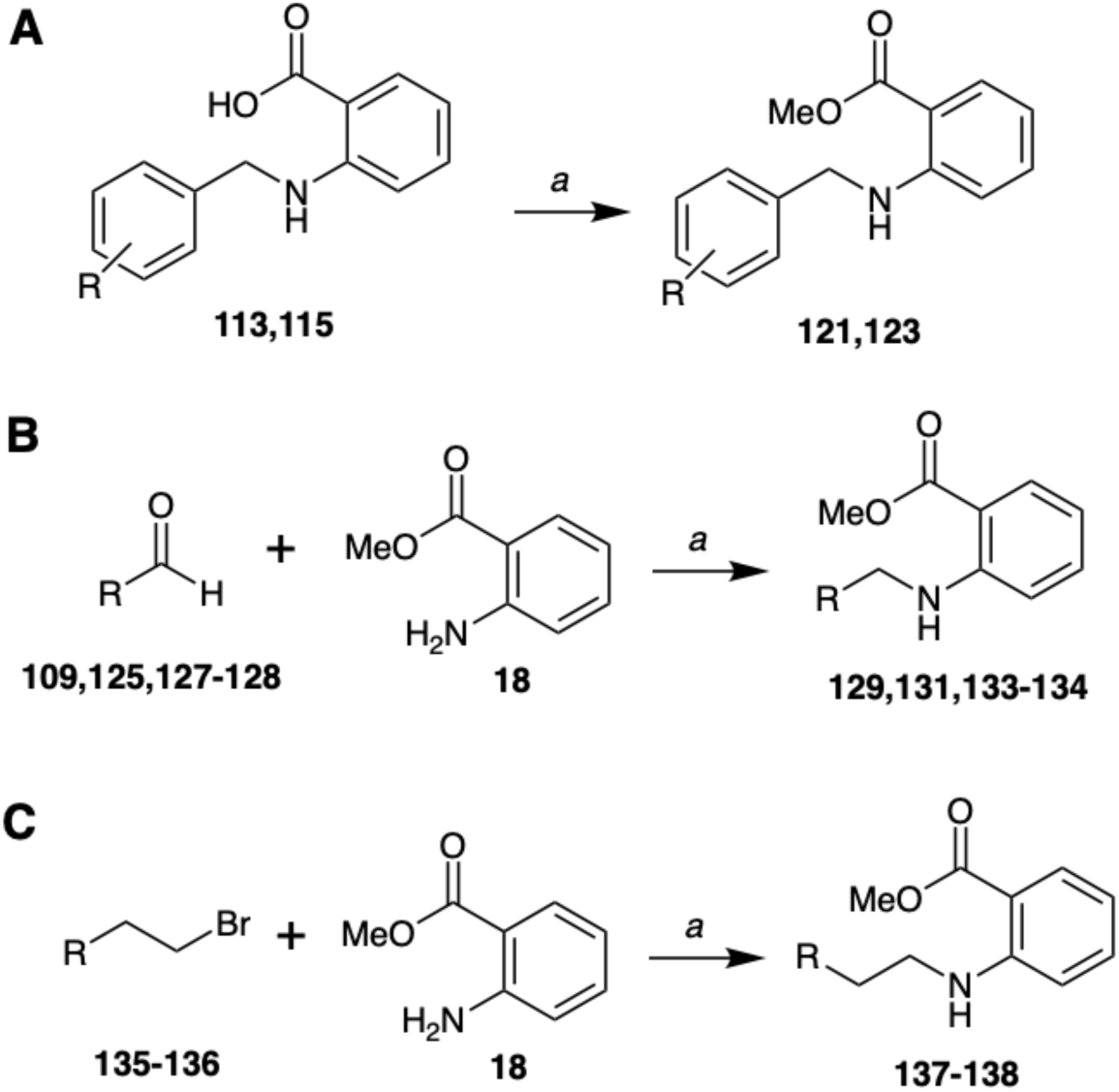
Synthesis of methyl(arylamino)benzamides. *Reagents and conditions:* **A***: a*: TBTU, DIPEA, DMF, room temp., 30 min.; then MeOH, room temp., 7h. **B***: a:* MeOH, reflux, 16h; then CH3COOH, NaBH4, room temp., 7h. **C:** *a:* CH3CN, reflux, 48h.

### Synthesis of (arylamino)benzoic acids (**113, 115, 117, 119, 120, 139-143, 156**)

A series of (arylamino)benzoic acids, structurally related to the methyl(arylamino)benzoates described above, were synthesised according to the methods outlined in **Scheme 7**. Derivatives **113, 115, 117, 119-120** were obtained via reductive amination of the appropriate aldehyde using *ortho* aminobenzoic acid, and sodium borohydride (NaBH_4_) as reducing agent. The carboxylic acid derivatives **139-143** were obtained via hydrolysis of the corresponding methyl esters. To investigate whether changing the disposition of the -CH_2_-NH-linker may affect the ability of compounds to occupy the cavity of c-FLIP, the inversion of the amine linker (compound **156**) was considered. Access to analogue **156** was secured via initial benzylic bromination of the starting methyl 2-methylbenzoate **165**, using N-bromosuccinimide (NBS) in the presence of azobisisobutyronitrile (AIBN, radical initiator), to give intermediate compound **161**. Reacting **161** with the aniline **153**, in presence of MeOH and NEt_3_., provided an intermediate ester, which was converted to **156** by basic hydrolysis.

**Scheme 7:**
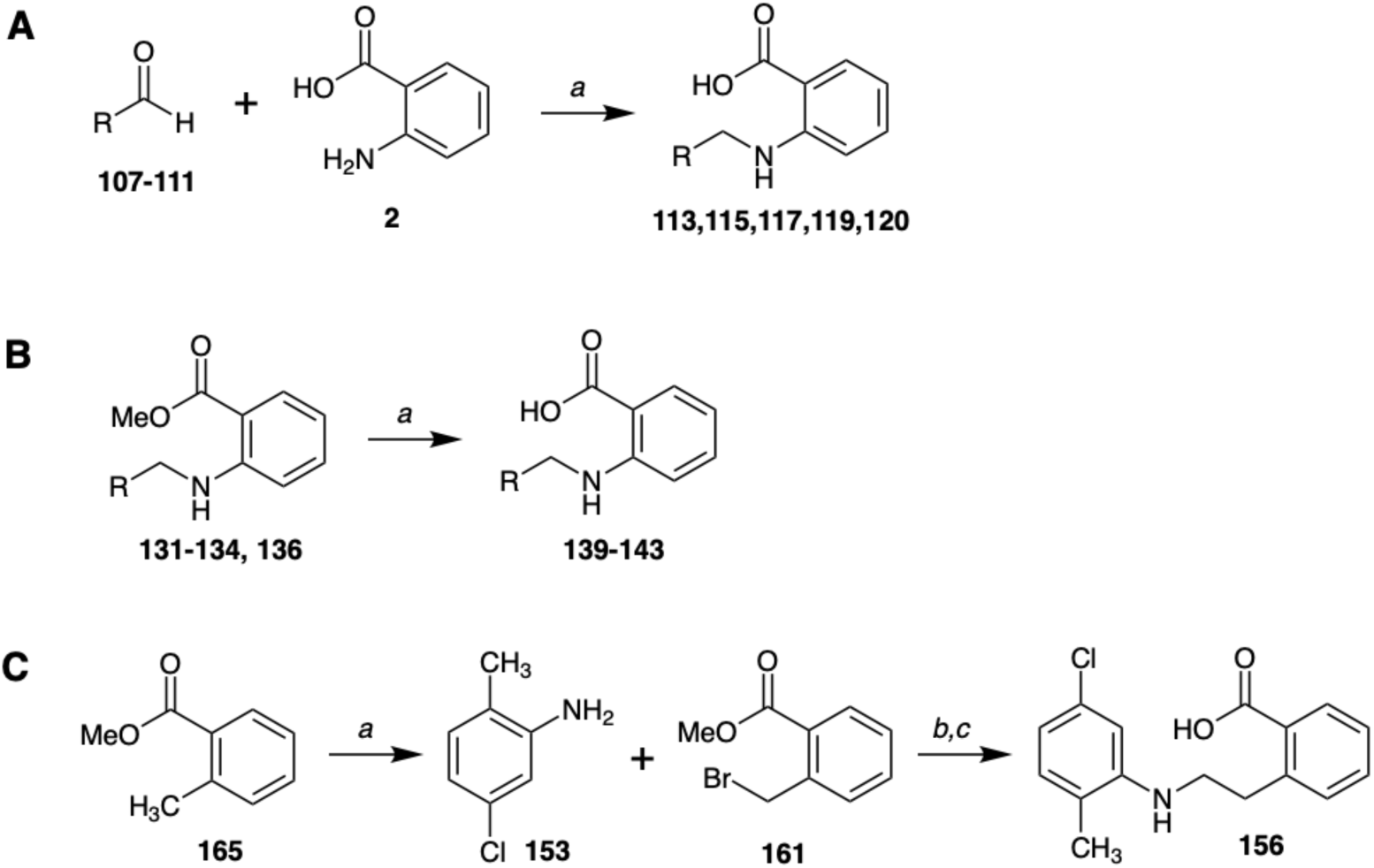
Synthesis of (arylamino)benzoic acids. *Reagents and conditions:* **A***: a*: MeOH, reflux, 16h; then CH3COOH, NaBH4, room temp., 7h. **B***: a:* LiOH, THF, MeOH, H2O, 70°C, 16h. **C:** *a:* NBS, AIBN, CHCl3, 70°C, 5h; *b:* NEt3, CH3OH, reflux, 16h. *c:* LiOH, THF, MeOH, H2O, 70°C, 16h.

### Synthesis of N-alkyl-2-(1H-tetrazol-5-yl)anilines (**144-149**) and 3-(2-((3-chloro-4-methylbenzyl)amino)phenyl-1,2,4-oxadiazol-5(4H)-one (**150**)

The reaction scheme to synthesise tetrazole (**144-149**) and oxadiazolone (**150**) analogues via reductive amination of intermediate anilines is shown in **Scheme 8**. Notably, an attempt was made to synthesise the corresponding oxadiazolone derivative with a sulfonamide linker, however this was not successful due to rapid degradation (less than 24h) of the product under ambient conditions.

**Scheme 8:**
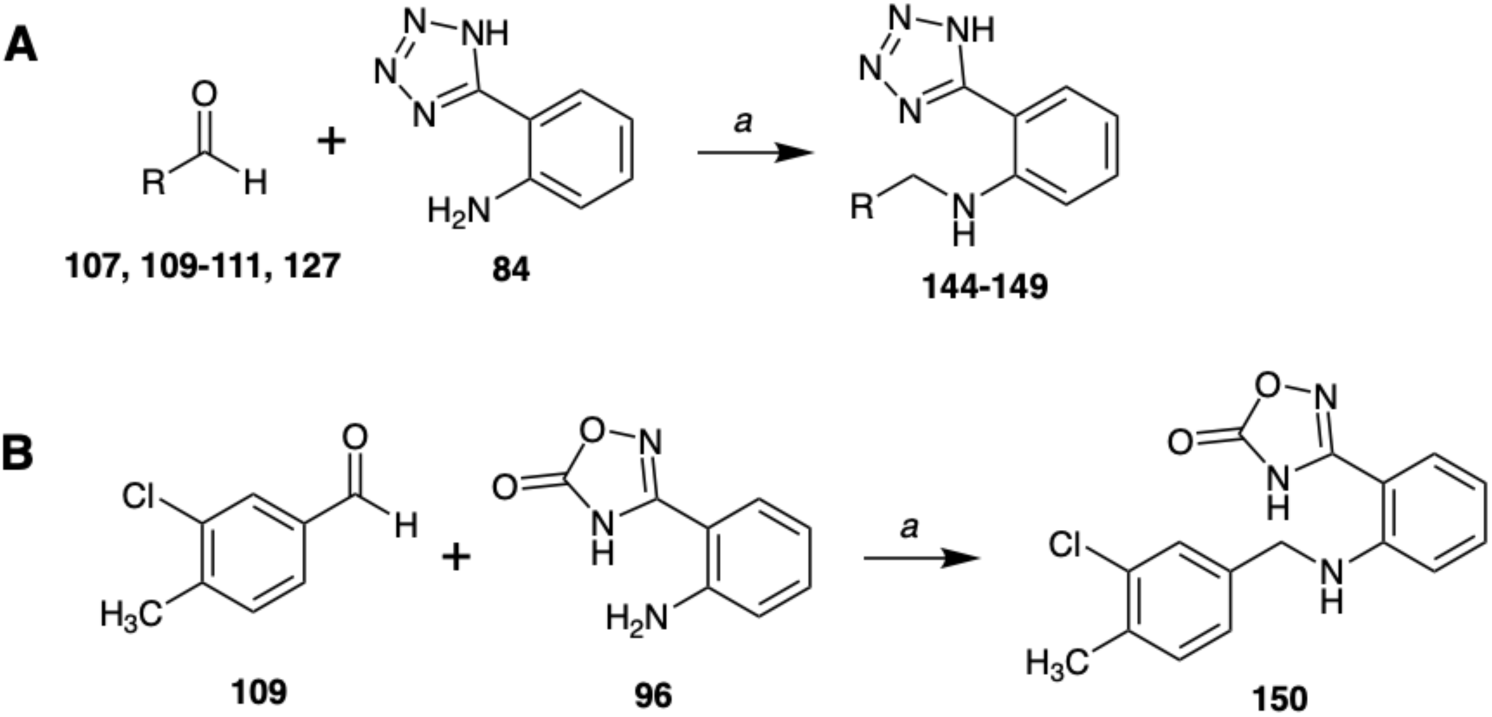
Synthesis of (arylamino)-N-(1H-tetrazol-5-yl)benzene derivatives and 3-(2-((3-chloro-4-methylbenzyl)amino)phenyl-1,2,4-oxadiazol-5(4H)-one (**150**). *Reagents and conditions:* **A:** *a:* MeOH, reflux, 16h; then CH3COOH, NaBH4, room temp., 7h. *B:* MeOH, reflux, 16h; then CH3COOH, NaBH4, room temp., 7h.

### Synthesis of diphenylmethylene derivatives (**169**, **171, 173**)

Further variation of the effect of the linker function between the two aryl rings was explored through the synthesis of compounds **169** and **173**, as shown in **Scheme 9**. A palladium-catalysed Suzuki-Miyaura reaction between arylboronic acids and benzylic bromides provided the required diphenylmethylene products 169 (37% yield) and 171 (56% yield). Methylbenzoate ester (**171**) was further hydrolysed to the corresponding benzoic acid derivative (**173**) for biological testing (67% yield).

**Scheme 9:**
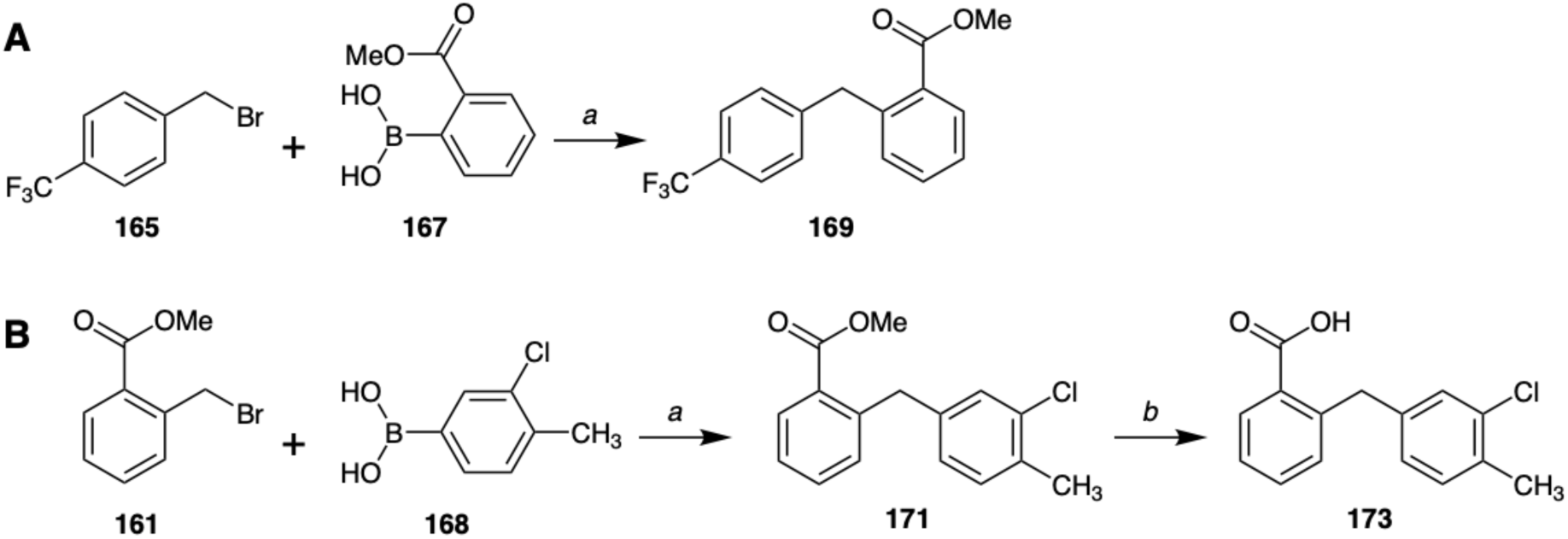
Synthesis of diphenylmethylene derivatives. **A:** *a:* Pd(OAc)2, PPh3, K3PO4, toluene, 80°C, 48h. **B:** *a:* Pd(PPh3)4, Na2CO3, EtOH, toluene, H2O, 80°C, 48h; *b:* LiOH, THF, MeOH, H2O, 70°C, 16h.

### Synthesis of N-phenylbenzamide derivatives (**179,182,186**)

Exploration the amide linker function as a comparator with the original sulfoamido linker was explored through the synthesis of N-phenylbenzamide derivatives (**179, 182, 186**). Methyl ester containing N-phenylbenzamides (**179, 182**) were readily synthesised via reaction of benzoyl chlorides (**174, 177**) with methyl 2-aminobenzoate (**18**). Finally, carboxylic acid containing N-phenylbenzamide **186** was synthesised via chlorination of carboxylic acid **184** followed by reaction of 3-chloro-4-methylbenzoyl chloride **185** with anthranilic acid **2**, as shown in **Scheme 10**.

**Scheme 10:**
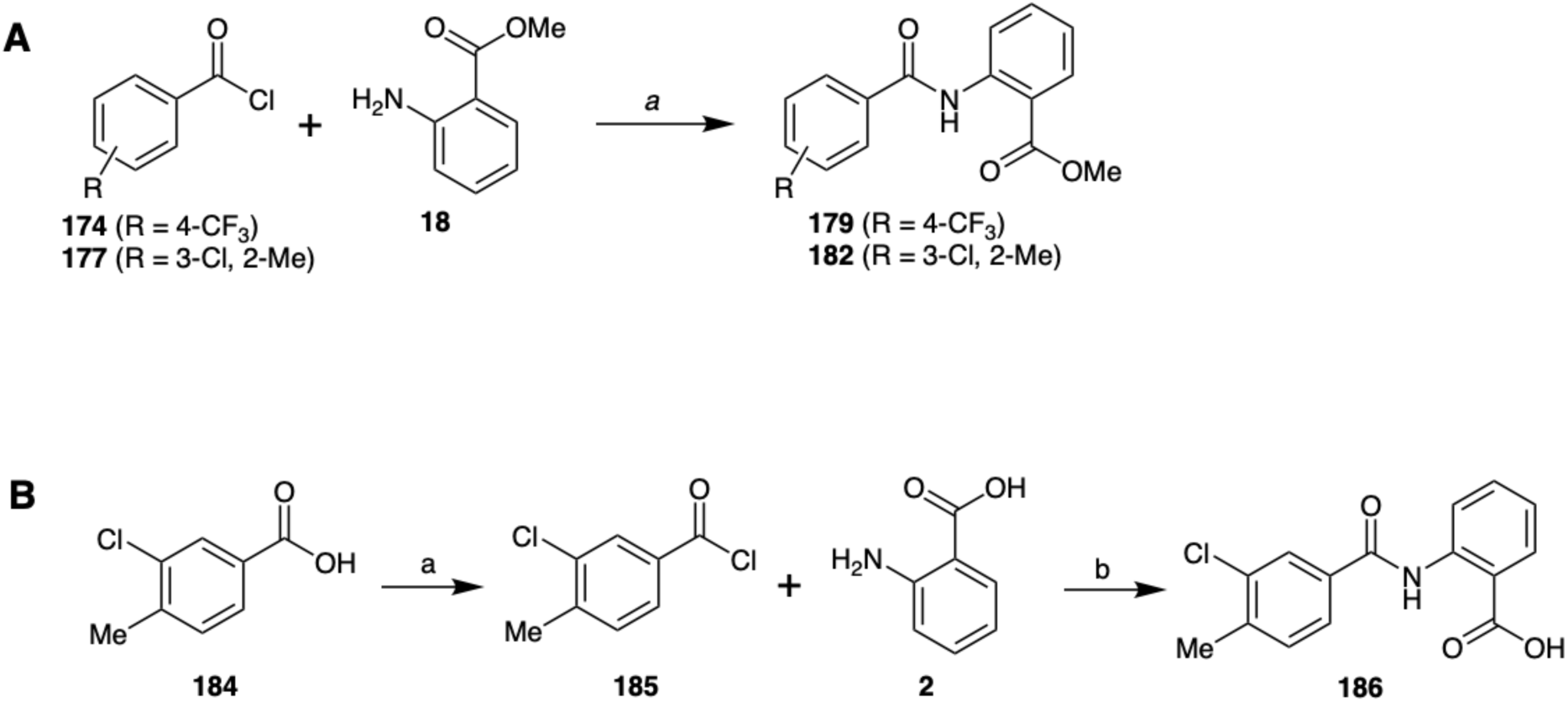
Synthesis of N-phenylbenzamide derivatives. **A:** NEt3, THF, 0°C to room temp., 6h. **B:** *a:* SOCl2, CH2Cl2, room temp., 24h; *b:* NEt3, CH2Cl2, 0°C to room temp., 16h.

#### Biological evaluation of the OH14 analogues

In order to determine the effects of the OH14 analogues on TRAIL sensitization in a biologically relevant context, MCF7 breast cancer cells were treated with the different compounds (10 μM), with or without the addition of 20ng/ml TRAIL (soluble human recombinant TRAIL, SuperKiller TRAIL, Enzo Life Sciences) and cell viability determined by colony formation assay (**Figure 6A-C**). The ability of the test compounds to sensitise cancer cells to TRAIL was measured as reduction of the number of colonies formed normalised to untreated controls. The parental compound **3** (OH14) was also evaluated, to compare the activity of the newly synthesised derivatives with the original hit compound. Additionally, cells were treated with compounds without TRAIL to assess the extent of cytotoxic or off-target effects of each compound. Those compounds reducing colony forming activity when combined with TRAIL, without affecting colony formation when administered alone, were of most interest, as they were concluded to be demonstrating TRAIL-dependent cytotoxicity. In addition to the parental compound OH14, 13 of the analogues tested exhibited significant reductions in colony forming ability compared to TRAIL stimulation alone (indicated by asterisks in **Figure 6**).

**Figure 6A-C.**
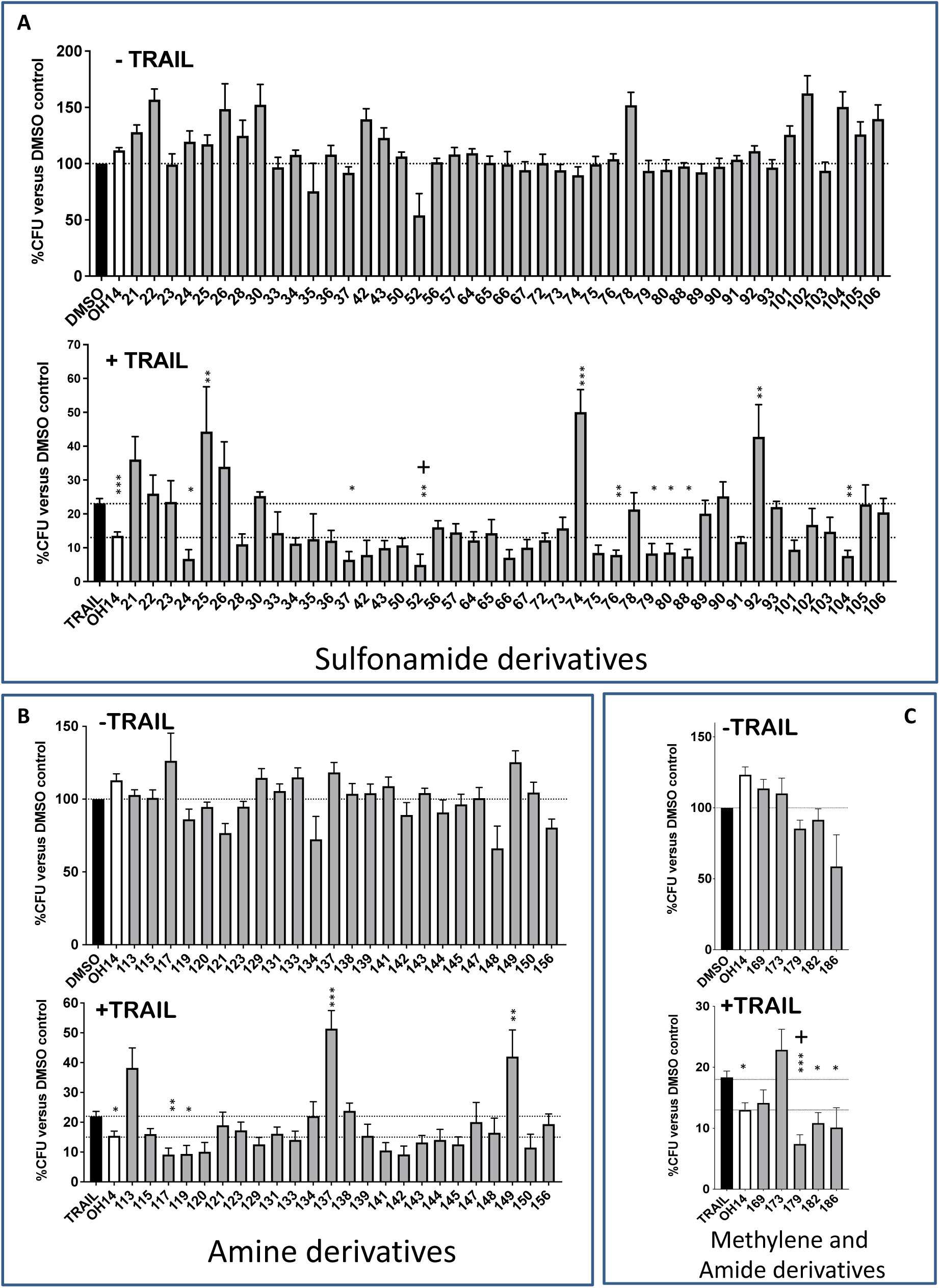
Comparison of OH14 analogues in the TRAIL-dependent clonogenic assay. MCF-7 cells were treated with compounds at 10 μM, alone or in combination with TRAIL (20 ng/mL). Colonies were counted after 10 days. %CFU was calculated from the number of cells plated and normalised to control (DMSO). Data shown is representative of at least three experiments. ± SEM. * p<0.05, ** p<0.005, *** p<0.0005. One-way ANOVA compared to TRAIL alone (black bar). Dashed lines represent the effect of no treatment (DMSO vehicle), TRAIL alone and OH14 in the presence of TRAIL, respectively. “+” p<0.05, two-tailed t-test (unpaired) of TRAIL-sensitising compounds compared to OH14+TRAIL.

#### Cytotoxicity of selected analogues

Cytotoxicity studies were performed for a selected set of analogues that showed a higher or equivalent level of selective TRAIL sensitisation compared to the original hit compound OH14 (**3**), to assess the lack of cytotoxic effects on non-cancerous cells. Analogues that showed significant cytotoxicity on MCF-7 breast cancer cells in the absence of TRAIL were also further re-profiled for cytotoxic effects. Compounds were additionally selected to represent the different analogue linker functions (sulfonamide, amine, methylene and amide) of the analogue series. Cytotoxicity studies were performed using the CellTiter-Blue Cell Viability Assay. The assay was performed on human embryonic kidney cells (HEK293), which are representative of non-tumorigenic cells (Figure 7).

**Figure 7:**
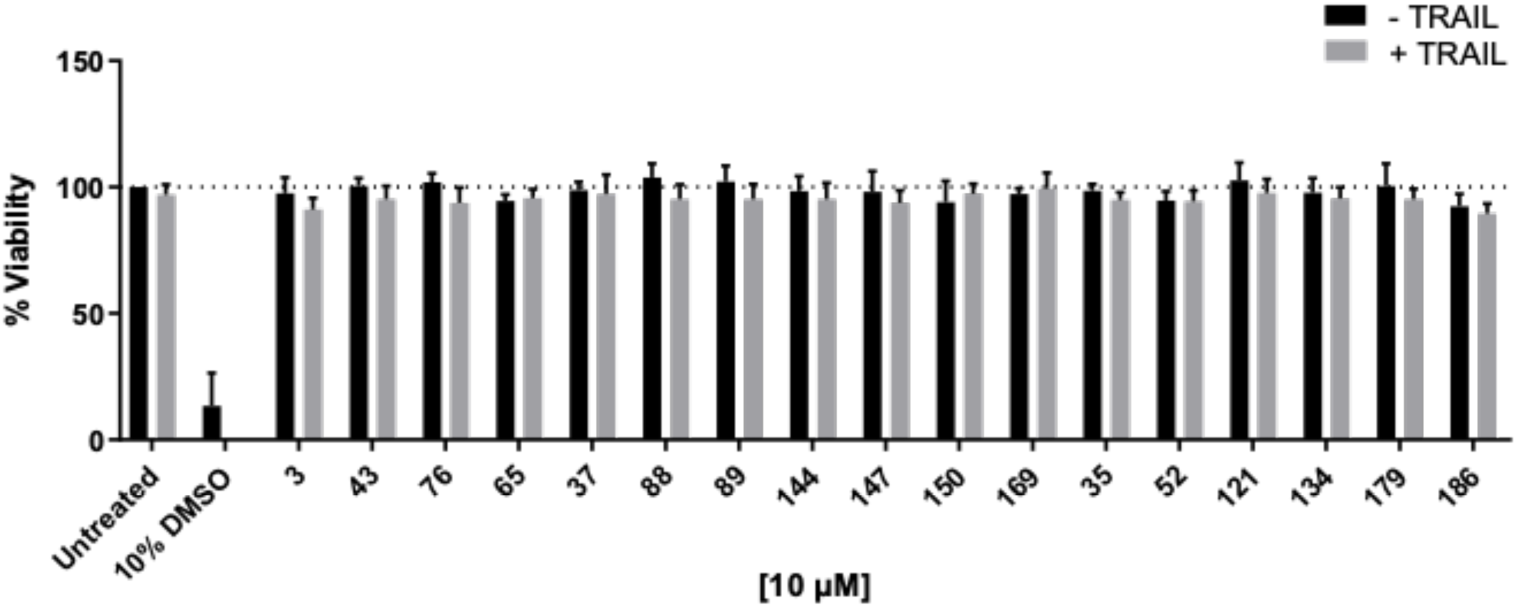
*Cell Viability Assay.* HEK293 cells were treated with compounds at 10 μM concentration, alone or in combination with TRAIL at 20 ng/mL. After 24 hours treatment the percentage of viable cells was calculated and normalised to the control (untreated TRAIL). 10% DMSO represents the positive control. Data shown is representative of at least 3 experiments ± SEM.

All the compounds showing ability to increase TRAIL sensitisation in the Colony Forming Assay (sulfonamides **43**, **76**, **65**, **37**, **88**, **89**; amines **144**, **147**, **150;** and diphenylmethylene **169**) did not induce a reduction of HEK293 cell viability compared to the untreated control, demonstrating a lack of cytotoxicity in this cell line. Compounds **35**, **52**, **53**, **121**, **134**, **170** and **186** were also investigated due to their ability to induce a reduction of colonies formation in the absence of TRAIL, but were also found not to have significant cytotoxic effects in the HEK293 cell line. Moreover, the non-toxicity of the original hit **3** was also confirmed, as well as the inability of TRAIL to induce cell death in non-cancerous cells.

#### Dose response study of selected tetrazole analogue of OH14 – compound 88

In order to further evaluate the activity of the compounds showing the best ability to increase TRAIL sensitisation at 10 μM concentration, we selected the tetrazole-containing analogue **88** for detailed dose-response profiling studies, as shown in **Figure 8**. This tetrazole analogue combined the properties of being amongst the most potent in terms of TRAIL sensitisation within MCF-7 breast cancer cells, alongside being devoid of cytotoxicity in the normal HEK293 cell line. Compound **88** was found to increase TRAIL sensitisation in a dose-dependent manner, with an IC_50_ value in the range of 15-19 μM. As shown in **Figure 8**, the effect of TRAIL alone (left graph) was dose-dependent, inducing a reduction of approximately 80% when tested at 20 ng/ml and 60% at the lowest concentration of 10 ng/ml. However, as demonstrated by the similar IC_50_ values calculated, **88** retained a similar ability to increase TRAIL sensitisation independently of the TRAIL concentration, thus indicating that the concentration of TRAIL does not significantly affect the activity of the compound.

**Figure 8:**
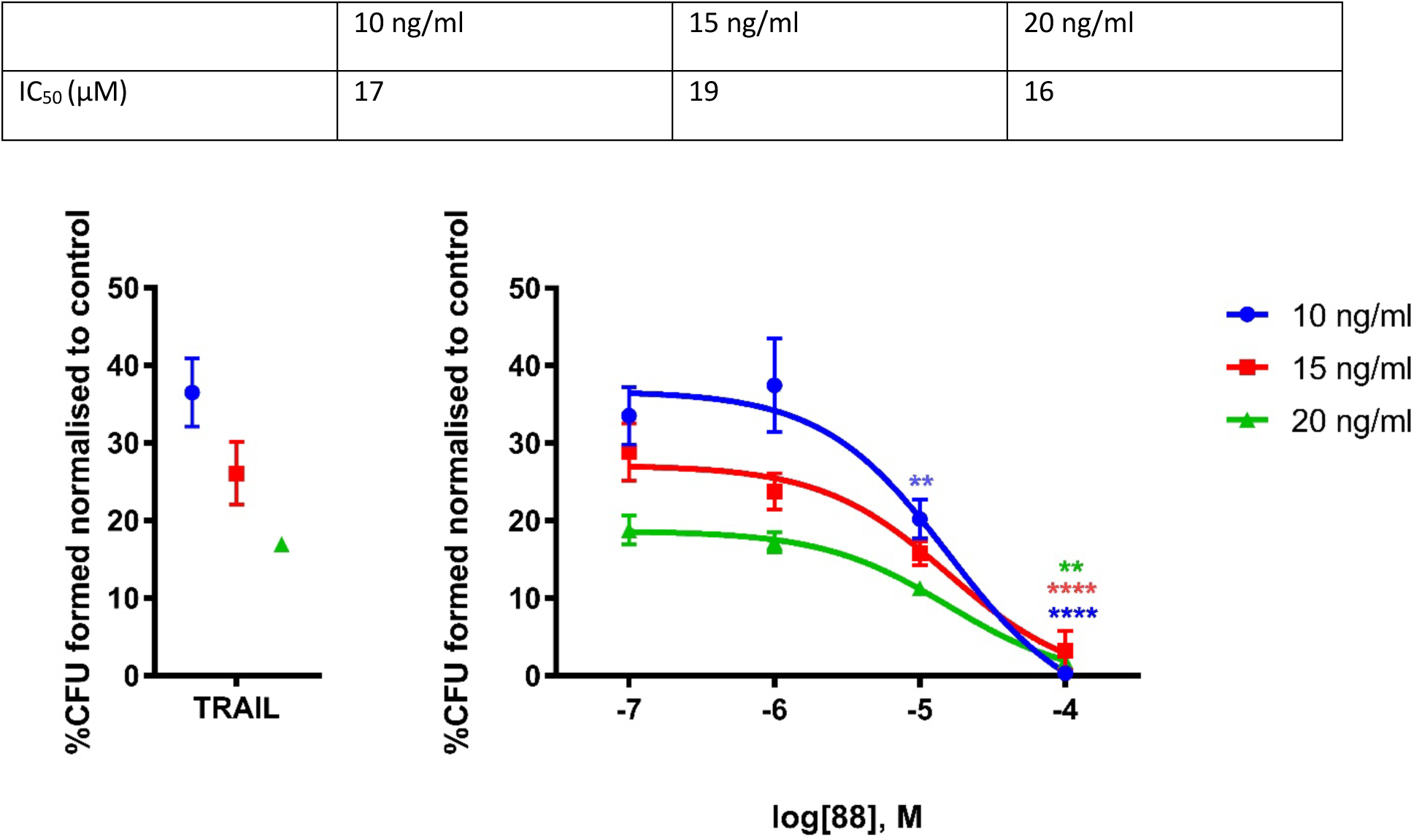
*Dose response curve and IC50 values for compound **88**.* IC50 values were calculated using GraphPad Prism 7.03, from the plot of log10(inhibitor concentration) vs % reduction in colony forming units (across three TRAIL concentrations).

#### In vitro early DMPK and toxicity studies

In order to investigate how the newly designed chemical modifications affected the pharmacokinetic properties of the compounds, some of the synthesised derivatives were subjected to preliminary *in vitro* pharmacokinetic studies. These studies were performed as an outsourced service by Cyprotex Ltd (Alderley Park, U.K.) and they include metabolic stability, Caco2 cell permeability and hERG inhibition. Results are shown in **Table 1**.

**Table 1:** In vitro DMPK and toxicity results.

| Compound | CL<sub>int</sub><br>(μl/min/mg<br>protein) | Standard<br>Error CL<sub>int</sub> | t<sub>1/2</sub><br>(min) | Mean<br>recovery (A2B) | Mean<br>recovery (B2A) | Efflux Ratio | IC<sub>50</sub> (μM) |
| --- | --- | --- | --- | --- | --- | --- | --- |
|  | Microsomal Stability Assay |  |  | Caco2 Cell Permeability Assay |  | hERG Inhibition Assay |  |
| **3** | 15.1 | 3.14 | 91.8 | 86.3 | 95.6 | 0.660 | >25 |
| **24** | Nd | Nd | Nd | Nd | nd | Nd | 23.6 |
| **79** | 79.5 | 12.7 | 17.4 | Nd | nd | Nd | >25 |
| **117** | 35.6 | 2.85 | 39.0 | 66.2 | 77.2 | 0.622 | >25 |
| **141** | 35.6 | 2.90 | 39.0 | 61.3 | 73.5 | 0.881 | >25 |
| **88** | 10.4 | 1.33 | 134 | 66.7 | 67.2 | 6.43 | >25 |
| **150** | 92.3 | 3.33 | 15.0 | 23.1 | 51.7 | 0.768 | 92.3 |

## Discussion

The continued development of targeted treatments for breast cancer is important for those patients who do not respond to current regimens and to improve on treatments with significant side effects. Although TRAIL exhibits low toxicity due to its notional specificity for cancer cells, most can acquire TRAIL-resistance mediated by cFLIP^3,10^. Prior to recent advances in molecular modelling, the consensus was that identification of a compound that would impair cFLIP function without targeting the structurally similar pro-apoptotic procaspase 8 may not be possible. Considering recent advances in our understanding of the mechanism of cFLIP incorporation into the DISC, opportunities to selectively target the DED domains on cFLIP are now coming to light^19^. Here we have focused molecular dynamic modelling on the DED1 domain as one of the critical interactive sites for cFLIP recruitment and have functionally confirmed the relevance of this interaction by mutagenesis. This allowed us to identify a specific small molecule inhibitor of cFLIP able to sensitise cancer cells including tumour-initiating cells to TRAIL. The relatively high (micromolar) concentration of OH14 (**3**) required to observe maximal effects on DISC recruitment, and moderate microsomal stability characteristics (**Supplementary Figure 6A and Table 1**) are likely to preclude it as an *in vivo* clinical candidate. Despite this however OH14 remains a useful pre-clinical tool for demonstrating proof-of-principle that pharmacological inhibition of cFLIP is sufficient to sensitize breast cancer cells, and in particular breast cancer stem cells, to TRAIL mediated killing and provides further proof-of-principle that pharmacological targeting of cFLIP is feasible.

It was originally thought that cFLIP exerts an inhibitory function in the DISC by primarily competing with procaspase-8 for binding to FADD.^20–23^ However, using a recombinant DISC model, Hughes *et al.* showed that increasing amounts of cFLIPL/S did not prevent procaspase 8 recruitment, casting doubt over the primacy of cFLIP’s direct role in binding FADD directly via its DED1 domain.^16^ In addition, recombinant cFLIP recruitment to FADD was shown to be procaspase 8-dependant.^5,16^ This suggested instead that cFLIP is primarily recruited indirectly to FADD by binding to procaspase ^8.16^

Previous mutational studies have been carried out to determine whether cFLIP binding occurs via DED1 or 2. Mutation of F144 in DED2 impaired cFLIP binding to FADD in a recombinant assay, in the absence of procaspase 8^16,23^. However, in the presence of procaspase 8, a low level of cFLIP F144 mutant was recruited to the DISC^14^. This was proposed to occur as a result of chain shortening, i.e., cFLIP with mutated DED2 can still bind to procaspase 8 but cannot recruit further molecules. These data suggested that cFLIP binds procaspase 8 via DED1, however, mutation of H7 in DED1 did not impair cFLIP recruitment^14,23^.

None of these studies modelled the interaction between cFLIP and procaspase 8, and therefore the DED1 mutations tested were limited to those predicted to interact with FADD (H7, E4, E11 and E46) ^21^. We build on the findings of Hughes *et al.* by modelling cFLIP interactions with procaspase 8. Our model supports the premise that cFLIP predominantly binds to procaspase 8 via DED1 and as expected, predicts different interacting residues to those described previously: K18, R38 and R45. We show that mutation of K18 and R45 residues suppressed the inhibitory function of cFLIP, suggesting impaired DISC recruitment. Whilst it is not possible to determine whether one or both of these residues are important, our findings support the prediction of Hughes *et al.* that cFLIP is incorporated into the DISC via the DED1 domain; a proposal recently validated by Fox et al who reported that H7A/R38D double mutant disrupts cFLIP binding to procaspase-8^23^. Furthermore, by addressing TRAIL sensitivity, we also provide direct evidence that DED1 is required for cFLIP function in physiological conditions.

Procaspase-8 is also incorporated into the DISC via DED1^16^ and the structural similarity between procaspase 8 and cFLIP suggests difficulties in selective targeting. However, by modelling DED1 interactions we were able to identify differences in the binding pockets of cFLIP and procaspase 8 that suggested differential targeting could be achieved; for example, the K18 and R45 residues are unique to the pocket of cFLIP. We used these models to identify OH14; a small molecule that binds to the pocket of cFLIP DED1 but not to that of procaspase 8. Using immunoprecipitation and FRET analysis we have shown that OH14 impairs cFLIP incorporation into the DISC as determined by recruitment to FADD. We go on to show that the R38 residue in the DED1 pocket is required for OH14 function, thus confirming target occupancy and function.

The fact that OH14 prevents cFLIP binding also further confirms the importance of cFLIP DED1 in DISC recruitment. We anticipate that an inhibitor designed to target DED2 of cFLIP would also impair the binding of cFLIP to FADD, and this has been demonstrated recently^24^ but that a cFLIP DED2 inhibitor would not prevent recruitment of cFLIP to procaspase 8 already present within the DISC, similar to the F144 mutant,^16^ and therefore functional efficiency may be reduced.

We show here that OH14 is an effective sensitising agent for TRAIL, allowing for the induction of caspase-dependent apoptosis in cancer cell lines. The degree of TRAIL sensitisation did vary across the breast cancer cell lines used. As these were chosen to be representative of different tumour subtypes, this may indicate that OH14/TRAIL is more effective for some breast tumours than others.

OH14 alone had no observable effect on cell viability. However, this has only been tested up to 100 μM for 3 hours, so it is formally possible that long-term inhibition of cFLIP binding may lead to disruption of protein stability and additional effects, for example in other pathways in which cFLIP is involved including NFkB and Wnt signalling.^4,25^

Based on these results we decided to prepare a series of OH14 analogues, to explore the activitu around this chemical scaffold. Taking into consideration all the biological data obtained for the sulfonamide derivatives, an initial structure-activity relationships analysis can be performed. Molecules retaining the original hydrophobic ring of OH14 (**3**) lost their activity when the carboxylic acid in *ortho* position of the anthranilic ring was replaced by a methyl ester in the *ortho (***21**), *meta (***22**) or *para (***23**) position. A reduction in activity was observed when an *ortho* isopropyl ester (**78**) or differently substituted *ortho* carboxamide groups (**102, 103, 105**), including the free carboxamide (**106**), were introduced. However, derivative **104**, which contains an *ortho* N,N-dimethyl carboxamide group, induced a slightly higher reduction of CFU in the presence of TRAIL compared to **3**. Groups such as *ortho* ethyl and *tert*-butyl esters (**50**, **80**) retained a similar activity compared to **3**, while the introduction of the tetrazole ring into the *ortho* position (**88**) induced a slightly improved ability to increase TRAIL sensitisation. For all the other derivatives, changes in both aromatic rings were explored. In general, the replacement of the hydrophobic phenylsulfonyl ring with heterocycles such as 2-furan, 2-thiophene and 3-pyridine were associated with activity retention in both carboxylic acid (**76, 65, 72**) and methyl ester derivatives (**57**, **34**). The introduction of a cyclohexane ring in both methyl ester and carboxylic acid analogues (**37** and **75**) induced a slight improvement in activity compared to **3.** Interesting results were obtained for the naphthalene derivatives. In the presence of a methyl ester group, both 2-and 1-naphthalene derivatives **35** and **36** showed a comparable TRAIL sensitising activity to **3**. The 2-napthalene derivative retained the activity in the presence of a 2-carboxylic acid (**73**) or of a tetrazole ring (**91**), while the carboxylic acid 1-napthalene derivative (**74**), completely lost its sensitising effect. The introduction of an 8-quinoline or 2-quinoline was associated with a slight improvement in activity in the presence of the carboxylic acid group (**66** and **67**), whereas both quinoline derivatives in the presence of the tetrazole ring (**92** and **93**) lost their TRAIL sensitising activity. All the derivatives showing an elongated arylsulfonyl linker (**33**, **56, 64**) demonstrated a comparable activity to **3**. Different hydrophobic substituents introduced on the phenylsulfonyl ring gave interesting results. The iodophenyl derivative **52** was considered as potentially cytotoxic, due to the significant reduction of CFU caused in the absence of TRAIL. On the contrary, groups such as the 4-trifluoromethyl (**24**), 5-chloro-3-methyl (**42**) and 3-chloro-4-methyl (**43**), containing a methyl ester, retained a similar or a slightly improved activity compared to **3**. The introduction of a tetrazole ring in the 4-trifluoromethylphenyl derivative (**90**) induced a complete loss of activity. Interesting results were obtained for the different derivatives of the 3-chloro-4-methylphenylsulfonyl analogue (**43**) synthesised. In contrast with the data obtained for hit compound **3**, the introduction of a tetrazole ring in *ortho* position (**89**) was associated with reduction of TRAIL sensitising activity. On the contrary, the analogous isopropyl ester derivative (**79**) and the methyl carboxamide derivative (**101**) retained a similar reduction in %CFU compared to **43** and hit **3**. The conflicting data obtained for analogues of TRAIL sensitising compounds **3** and **43**, in addition to all the results discussed above, suggest that the difference in activity observed for the synthesised derivatives is not related to a single modification or to a particular group, but it seems to be associated with a combination of the changes at both aromatic rings.

In accordance with the data obtained for the sulfonamide linker derivatives, the different biological activity of the amine derivatives seems to be correlated with modifications on both aromatic rings. Analogue **129**, the amine linker derivative of **43,** which contains a 3-chloro-4-methylphenyl ring and an *ortho* methyl ester, retained similar activity compared to the original sulfonamide compound **43** and to original hit compound OH14 (**3**). The substitution of a carboxylic acid (**117**), a tetrazole (**145)** and a dihydro-oxadiazole (**150**) group was associated with a slight improvement of TRAIL sensitising activity compared to **3**. The introduction of a 3-pyridine ring was associated with activity retention in the carboxylic acid group (**139**) and corresponding methyl ester (**131**) analogues. The 2- and 1-naphthalene derivatives (**141** and **142**) as well as the 8- and 2-quinoline derivatives (**119** and **120**), in the presence of a 2-carboxylic acid, showed a similar activity compared to the original hit compound OH14 (**3**), although a slight reduction of the CFU was observed for the 1-napthalene (**142**) and the 8-quinoline (**119**) derivatives in the absence of TRAIL. Interestingly, for the 2-methyl ester analogues, while the 2-napthalene **133** retained activity, the 1-napthalene **134** completely lost its TRAIL sensitising effect, increasing instead its ability to induce a reduction of CFU in the absence of TRAIL. The introduction of a tetrazole ring in the 2-napthalene derivative (**149**) induced a complete loss of ability to induce TRAIL sensitisation, whilst for the 2-quinoline derivative (**148**) the presence of a tetrazole ring was associated with a reduction of CFU in the absence of TRAIL. Compound **121**, characterised by a 4-trifluoromethylphenyl ring and a 2-methyl ester, although retaining a similar activity compared to **3**, also induced a reduction of the CFU formed in the absence of TRAIL, indicating potential additional off-target effects. Interestingly, while the replacement of the methyl ester of 4-trifluoromethylphenyl analogue **121** with a carboxylic acid group (**113**) was associated with loss of activity, the introduction of a tetrazole ring in **144** induced a reduction of the CFU comparable to **3**, contrary to the results obtained for the corresponding sulfonamide derivative (**90**). Controversial results were obtained also for the (4-trifluoromethyl)benzyl derivatives showing a longer amine linker, with the carboxylic acid derivative **143** showing a similar activity compared to **3**, while the corresponding methyl ester **138**, as well as the unsubstituted benzyl ring (**137**), did not show any TRAIL sensitising effect. Similarly, the inversion of the amine linker in **156** only showed a small sensitising effect, associated with a reduction of the CFU even in the absence of TRAIL.

The introduction of a methylene linker in derivatives containing a 3-chloro-4-methylphenyl ring (**173**) was associated with loss of TRAIL sensitisation activity compared to the corresponding sulfonamide- and amine-linker derivatives. On the contrary, the 4-trifluoromethylphenyl methylene derivative (**169)** retained a similar activity compared to **3**. Contrary results were obtained for the amide derivatives. A slight ability to induce a reduction of colonies formation in the absence of TRAIL was also observed for 2-methyl ester analogue **182**. The 2-methyl ester analogue **179**, containing a 4-trifluoromethylphenyl group, showed an increased TRAIL sensitisation activity compared to **3**, but also induced a reduction of the CFU in the absence of TRAIL. A similar effect was observed for the 3-chloro-4-methylphenyl derivative containing a 2-carboxylic acid group, **186**.

All the compounds showing ability to increase TRAIL sensitisation in the Colony Forming Assay (**43**, **76**, **65**, **37**, **88**, **89**, **144**, **147**, **150** and **169**) did not induce a reduction of cell viability compared to the control, demonstrating lack of cytotoxic effects. Compounds **35**, **52**, **53**, **121**, **134**, **170** and **186** were also investigated due to their ability to induce a reduction of colonies formation in the absence of TRAIL. Moreover, the non-toxicity of the original hit **3** was also confirmed, as well as the inability of TRAIL to induce cell death in non-cancerous cells.

From the DMPK results, we can make these observations. In general, all the compounds tested did not show any significant ability to inhibit hERG channels, thus suggesting lack of cardiotoxic effects. Good permeability in both directions across the Caco2 cells was observed for all compounds, indicating a predicted oral absorption potential. In terms of microsomal stability, most of the compounds showed a higher clearance and a shorter half-life time compared to **3**, whilst **88**, the *ortho* tetrazole derivative of **3**, was found to be more metabolically stable, increasing the half-life time of **3** from approximatively 92 mins to 134 mins.

While the key finding of this study is the successful pharmacological targeting of cancer cell activity via selective targeting of the cFLIP/TRAIL pathway, a limitation of this particular inhibitor is that this was only achievable at relatively high (75-100μM) concentrations. Therefore, we do not anticipate that OH14 in its current form will be a viable drug for clinical use but will instead provide a useful pre-clinical tool and a structural platform for further analogue development.

Thus, in pursuing the development of a targeted cFLIP inhibitor we have been able to demonstrate the importance of the DED1 domain for cFLIP function and use this knowledge to develop OH14. This study has shown that molecular inhibition of cFLIP in combination with TRAIL is a viable and effective strategy for targeting cancer cells.

## Methods

Experimental procedures and methods are fully described in the Supplemental Information file.

## Additional Information

### Acknowledgements

We would like to thank Dr Rob Clarke (University of Manchester, U.K.), Dr Ladislav Andera (University of Prague) and Dr Julia Gee (Cardiff University) for their kind gifts of cell lines used in this study. We would also like to thank Dr Michele Hughes (MRC Toxicology Unit, U.K.) for their help and advice throughout the study.

### Authors’ contributions

RC, AB and ADW conceived of the study, interpreted results, contributed to and revised the manuscript. RF designed experiments, acquired data, interpreted results and drafted the manuscript. OH, KYL, TR, ARS, GG, and AV designed experiments, acquired data, interpreted results and revised the manuscript. MM interpreted results and revised the manuscript.

### Ethics approval

Mouse experiments were completed in accordance with ARRIVE2.0 guidelines and approved by Cardiff University School of Biosciences Ethics Committee.

### Data availability

All datasets are securely archived and are available to researchers on request via the corresponding author.

## References

1. Greco S., Fabbri N., Spaggiari R., De Giorgi A., Fabbian F., Giovine A. Update on classic and novel approaches in metastatic triple-negative breast cancer treatment: a comprehensive review. Biomedicines 11(6): 1772 (2023).

2. Anderson R.L., Balasas T., Callaghan J., Coombes R.C., Evans J., Hall J.A., et al. A framework for the development of effective anti-metastatic agents. Nat. Rev. Clin. Oncol. 16(**3**), 185 (2019).

3. Piggott L., Omidvar N., Martí Pérez S., French R., Eberl M., Clarkson R.W. Suppression of apoptosis inhibitor c-FLIP selectively eliminates breast cancer stem cell activity in response to the anti-cancer agent TRAIL. Breast Cancer Res. 14;13(5):R88 (2011).

4. French R., Hayward O., Jones S., Yang W., Clarkson R.W. Cytoplasmic levels of cFLIP determine a broad susceptibility of breast cancer stem/progenitor-like cells to TRAIL. Mol Cancer 15;14:209 (2015).

5. Dickens L.S., Boyd R.S., Jukes-Jones R., Hughes M.A., Robinson G.L., Fairall L., et al. A death effector domain chain DISC model reveals a crucial role for caspase-8 chain assembly in mediating apoptotic cell death. Mol Cell 27;47(2):291–305 (2012).

6. Piggott L., Silva A., Robinson T., Santiago-Gómez A., Simões BM., Becker M., et al. Acquired resistance of ER-positive breast cancer to endocrine treatment confers an adaptive sensitivity to TRAIL through posttranslational downregulation of c-FLIP. Clin Cancer Res. 15;24(10):2452–2463 (2018).

7. Rahman M., Davis S.R., Pumphrey J.G., Bao J., Nau M.M., Meltzer P.S., et al. TRAIL induces apoptosis in triple-negative breast cancer cells with a mesenchymal phenotype. Breast Cancer Res Treat 113(2):217–30 (2009).

8. Forero-Torres A., Varley K.E., Abramson V.G., Li Y., Vaklavas C., Lin N.U., et al. Translational Breast Cancer Research Consortium (TBCRC). TBCRC 019: A phase II trial of nanoparticle albumin-bound paclitaxel with or without the anti-death receptor 5 monoclonal antibody tigatuzumab in patients with triple-negative breast cancer. Clin Cancer Res 15;21(12):2722–9 (2015).

9. Soria J.C., Márk Z., Zatloukal P., Szima B., Albert I., Juhász E., Pujol J.L., Kozielski J., Baker N., Smethurst D., Hei Y.J., Ashkenazi A., Stern H., Amler L., Pan Y., Blackhall F. Randomized phase II study of dulanermin in combination with paclitaxel, carboplatin, and bevacizumab in advanced non-small-cell lung cancer. J Clin Oncol 20;29(33):4442–51 (2011).

10. Yee L., Burris H.A., Kozloff M., Wainberg Z., Pao M., Skettino S. et al. Phase Ib study of recombinant human Apo2L/TRAIL plus irinotecan and cetuximab or FOLFIRI in metastatic colorectal cancer (mCRC) patients (pts); preliminary results. J Clin Oncol 27:4129 (2009).

11. Yee L., Fanale M., Dimick K., Calvert S., Robins C., Ing J. et al. A phase IB safety and pharmacokinetic (PK) study of recombinant human Apo2L/TRAIL in combination with rituximab in patients with low-grade non-Hodgkin lymphoma. J Clin Oncol 25:8078 (2010).

12. Day T.W., Sinn A.L., Huang S., Pollok K.E., Sandusky G.E., Safa A.R. c-FLIP gene silencing eliminates tumour cells in breast cancer xenografts without affecting stromal cells. Anticancer Res 29(10):3883–6 (2009).

13. Frew A.J., Lindemann R.K., Martin B.P., Clarke C.J., Sharkey J., Anthony D.A., et al. Combination therapy of established cancer using a histone deacetylase inhibitor and a TRAIL receptor agonist. Proc Natl Acad Sci U S A 12;105(32):11317–22 (2008).

14. Jiang Z., Li W., Hu X., Zhang Q., Sun T., Cui S., et al. Tucidinostat plus exemestane for postmenopausal patients with advanced, hormone receptor-positive breast cancer (ACE): a randomised, double-blind, placebo-controlled, phase 3 trial. Lancet Oncol. 20(6):806–815 (2019).

15. Shah, R.R. Safety and tolerability of histone deacetylase (HDAC) inhibitors in oncology. Drug Saf 42;235–245 (2019).

16. Hughes M.A., Powley I.R., Jukes-Jones R., Horn S., Feoktistova M., Fairall L., et al. Co-operative and hierarchical binding of c-FLIP and caspase-8: a unified model defines how c-FLIP isoforms differentially control cell fate. Mol Cell 17;61(6):834–49 (2016).

17. Hwang E.Y., Jeong M.S., Park S.Y., Jang S.B. Evidence of complex formation between FADD and c-FLIP death effector domains for the death inducing signaling complex. BMB Rep 47(9):488–93 (2014).

18. Fu T.M., Li Y., Lu A., Li Z., Vajjhala P.R., Cruz A.C., et al. Cryo-EM structure of caspase-8 tandem DED filament reveals assembly and regulation mechanisms of the death-inducing signalling complex. Mol Cell. 64 (2): 236–250 (2016).

19. Majkut J., Sgobba M., Holohan C., Crawford N., Logan A.E., Kerr E., et al. Differential affinity of FLIP and procaspase 8 for FADD’s DED binding surfaces regulates DISC assembly. Nat Commun 28;5:3350 (2014).

20. Yang J.K., Wang L., Zheng L., Wan F., Ahmed M., Lenardo M.J., et al. Crystal structure of MC159 reveals molecular mechanism of DISC assembly and FLIP inhibition Mol Cell 22;20(6):939–49 (2005).

21. Eberstadt M., Huang B., Chen Z., Meadows R.P., Ng S.C., Zheng L., et al. NMR structure and mutagenesis of the FADD (Mort1) death-effector domain. Nature 392:941–945 (1998).

22. Carrington P.E., Sandu C., Wei Y., Hill J.M., Morisawa G., Huang T., et al. The structure of FADD and its mode of interaction with procaspase-8. Mol Cell 22(5):599–610 (2006).

23. Fox J.L., Hughes M.A., Fairall L., Schwabe J.W.R., Morone N., Cain K., MacFarlane M. Cryo-EM structural analysis of FADD:Caspase-8 complexes defines the catalytic domain architecture for co-ordinated control of cell fate. Nature Comm. 12: 819. doi: 10.1038/s41467-020-20806-9 (2021)

24. Hillert L.K., Ivanisenko N.V., Busse D. et al. Dissecting DISC regulation via pharmacological targeting of caspase-8/c-FLIP_L_ heterodimer. Cell Death Differ 2;2117–2130 (2020)

25. Golks A., Brenner D., Krammer P.H., Lavrik I.N. The c-FLIP-NH2 terminus (p22-FLIP) induces NF-kappaB activation. J Exp Med 15;203(5):1295–305 (2006)

