## supplemental information for "The Discovery of Small Molecule Inhibitors of cFLIP that Sensitise Tumour Cells to TRAIL"

### SUPPLEMENTARY DATA

α1 · α2 · α3

| | |
|---|---|
| FADD-DED1 | M D P F L V L L H S V S S S L S S S E L T E L K F L C L G R - V G K R K L E R V - - - - - - |
| Procaspase-8-DED1 | - M D F S R N L Y D I G E Q L D S E D L A S L K F L S L D Y - I P Q R K Q E P I - - - - - - |
| Procaspase-8-DED2 | I S A Y R V M L Y Q I S E E V S R S E L R S F K F L L Q E E - I S K C K L D D D - - - - - - |
| cFLIP-DED1 | - - M S A E V I H Q V E E A L D T D E K E M L L F L C R D - - V - - - A I D V V P - - - - - - |
| cFLIP-DED2 | - - - Y R V L M A E I G E D L D K S D V S S L I F L M K D Y - M G R G K I S K E K - - - - - - |
| MC159-DED1 | - V P S L P F L R H L L E E L D S H E D S L L L F L C H D A A P G C T T V T Q - - - - - - - |
| MC159-DED2 | - - R Y R K L M V C V G E E L D S S E L R A L R L F A C - - N L - N P S L S T A L S E S - - |

α4 · α5 · α6

| | |
|---|---|
| FADD-DED1 | Q S G L D L F S M L L E Q N D L E P G H T E L L R E L L A S L - - - R R H D L L R R V D D F E - |
| Procaspase-8-DED1 | K D A L M L F Q R L Q E K R M L E E S N L S F L K E L L F R I - - - N R L D L L I T Y L N T R - |
| Procaspase-8-DED2 | M N L L D I F I E M E K R V I L G E G K L D I L K R V C A Q I - - - - N K S L L K I I N D Y - - |
| c-FLIP-DED1 | P N V R D L L D I L R E R G K L S V G D - - - L A E L L Y R V - - - R R F D L L K R I L K M D R |
| cFLIP-DED2 | - S F L D L V V E L E K L N L V A P D Q L D L L E K C L K N I - - - H R I D L K T K I Q K Y K Q |
| MC159-DED1 | - - - - - A L C S L S Q Q R K L - - - T L A A L V E M L Y V L - - - Q R M D L L K S R F G L S K |
| MC159-DED2 | S R F V E L V L A L E N V G L V S P S S V S V L A D M L R T L - - - R R L D L C Q Q L V E Y E Q |

**B**

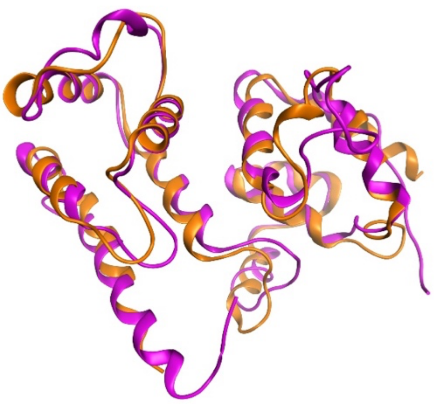

**C**

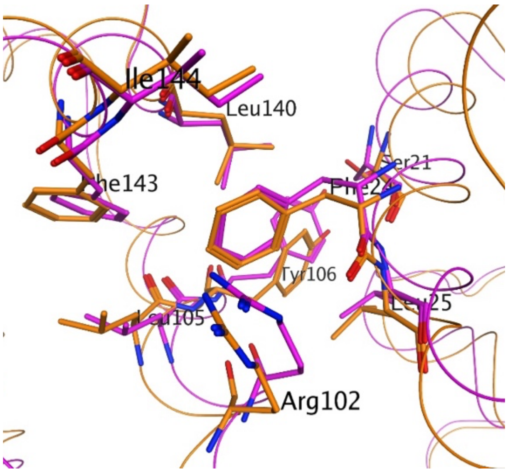


**Supplementary Figure 1: A.** *DED Amino acid sequence comparison with FADD, procaspase-8, c-FLIP and MC159*. Sequences have been aligned based on DEDs. Shared 'FL' sequence highlighted in yellow. Amino-acid sequences were obtained from Eberstadt, *et al.*, 1998. Table adapted from Eberstadt *et al.*, 1998[19]; **B.** Superposition of homology model of caspase-8, using the MC159 template (purple) and the caspase-8 crystal structure (orange - PDB:4ZBW); **C.** Superposition of the residues around the FL domain between the caspase-8 model (purple) and crystal structure (orange).

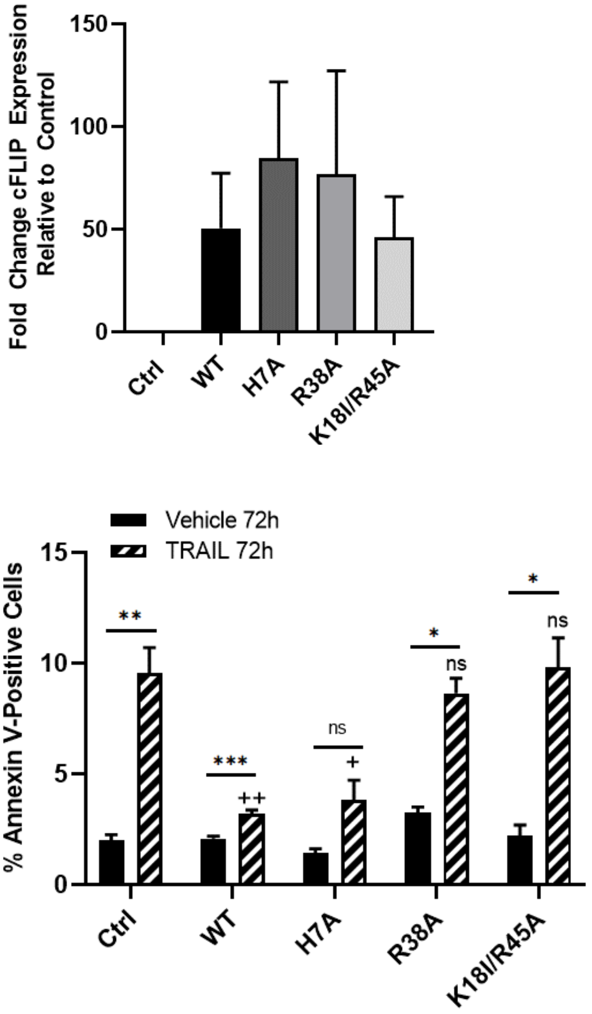


**Supplementary Figure 2: A.** Semi-quantitative evaluation of relative total levels of cFLIP determined by densitometry across three independent western blots for total cFLIP levels in HeLa cells stably transfected with WT or mutant cFLIPL and harvested from passages used in the viability assay. Band intensities within western blots were normalised against GAPDH and then expressed as relative normalised intensity compared to control (untransfected cells). Error bars represent standard deviation; **B.** HeLa cells stably transfected with WT or mutant cFLIPL (supplementary Figure 2A) were treated with TRAIL for 72 hours and cell death determined by annexin V staining using Incucyte real-time analysis. Cell death is expressed as the percentage of annexin positive cells following treatment and error bars represent standard error of the mean from three independent experiments. * = $p<0.05$, ** = $p<0.01$ *** = $p<0.001$ between Vehicle and TRAIL treated for each construct; + = $p<0.05$, ++ = $p<0.01$ between control TRAIL and mutant construct TRAIL.

| | | | | | |
|---|---|---|---|---|---|
| Compound OH1<br><br>LogP: 2.87<br>H-d: 2<br>H-a: 4 | $C_{16}H_{15}NO_3$<br>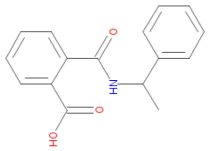<br>mw269.3 | Compound OH7<br><br>LogP: 3.3<br>H-d: 2<br>H-a: 4 | $C_{17}H_{17}NO_4$<br>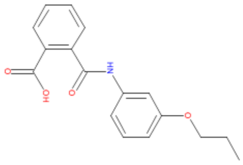<br>mw299.33 | Compound OH13<br><br>LogP: 4.7<br>H-d: 2<br>H-a: 4 | $C_{18}H_{11}Cl_2NO_3$<br>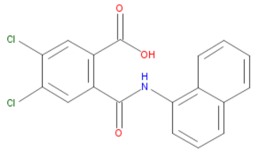<br>mw360.2 |
| Compound OH2<br><br>LogP: 3.3<br>H-d: 1<br>H-a: 3 | $C_{16}H_{11}ClO_3$<br>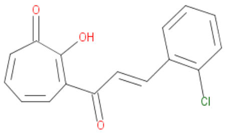<br>mw286.71 | Compound OH8<br><br>LogP: 5.15<br>H-d: 2<br>H-a: 4 | $C_{16}H_{10}INO_3S_2$<br>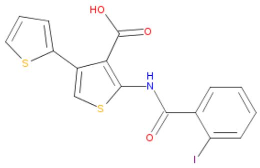<br>mw455.3 | Compound OH14<br><br>LogP: 4.01<br>H-d: 2<br>H-a: 5 | $C_{14}H_{11}Cl_2NO_4S$<br>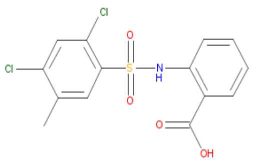<br>mw360.22 |
| Compound OH3<br><br>LogP: 3.95<br>H-d: 1<br>H-a: 3 | $C_{16}H_{10}Cl_2O_3$<br>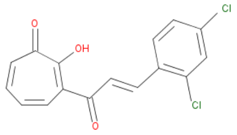<br>mw321.16 | Compound OH9<br><br>LogP: 3.46<br>H-d: 2<br>H-a: 4 | $C_{21}H_{17}NO_4$<br>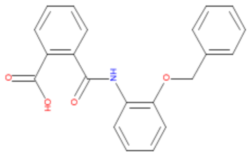<br>mw347.37 | Compound OH15<br><br>LogP: 4.01<br>H-d: 2<br>H-a: 5 | $C_{22}H_{23}FN_2O_3$<br><br>**Structure not available**<br><br>mw382.43 |
| Compound OH4<br><br>LogP: 4.66<br>H-d: 2<br>H-a: 6 | $C_{19}H_{25}NO_5S$<br>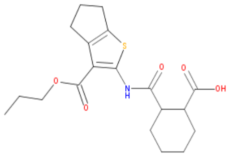<br>mw397.48 | Compound OH10<br><br>LogP: 5.35<br>H-d: 2<br>H-a: | $C_{17}H_{14}N_2O_2$<br>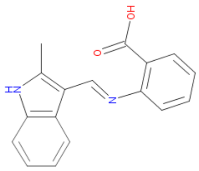<br>mw278.31 | Compound OH16<br><br>LogP: 4.01<br>H-d: 2<br>H-a: 5 | $C_{17}H_{18}N_4O_3S$<br>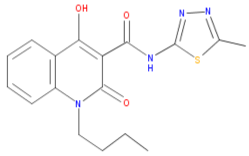<br>mw358.42 |
| Compound OH5<br><br>LogP: 2.6<br>H-d: 2<br>H-a: 4 | $C_{17}H_{15}NO_4$<br>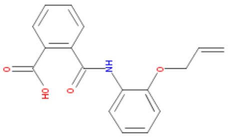<br>mw297.31 | Compound OH11<br><br>LogP: 3.23<br>H-d: 2<br>H-a: 5 | $C_{20}H_{19}FN_2O_3$<br>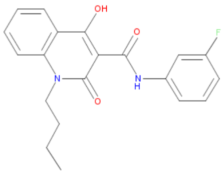<br>mw354.38 | Compound OH17<br><br>LogP: 4.01<br>H-d: 2<br>H-a: 5 | $C_{19}H_{14}BrNO_3S$<br>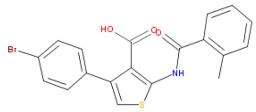<br>mw416.3 |
| Compound 6<br><br>LogP: 1.18<br>H-d: 3<br>H-a: 7 | $C_{18}H_{18}N_2O_5$<br>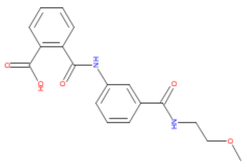<br>mw342.35 | Compound OH12<br><br>LogP: 5.58<br>H-d: 2<br>H-a: 5 | $C_{21}H_{22}N_2O_3$<br>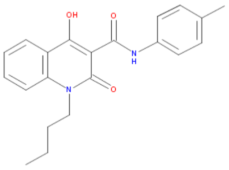<br>mw350.42 | Compound OH18<br><br>LogP: 4.01<br>H-d: 2<br>H-a: 5 | $C_{17}H_{19}NO_3S$<br><br>**Structure not available**<br><br>mw317.41 |
| | | | | Compound OH19 | $C_{18}H_{15}NO_4$ |

**Supplementary Figure 3:** Structure and properties of selected compounds from in silico screen. The 19 Selected compounds all adhered to the Lipinski rules. H-d = Hydrogen Donors, H-a = Hydrogen Acceptors. LogP = partition coefficient (measure of hydrophilicity).

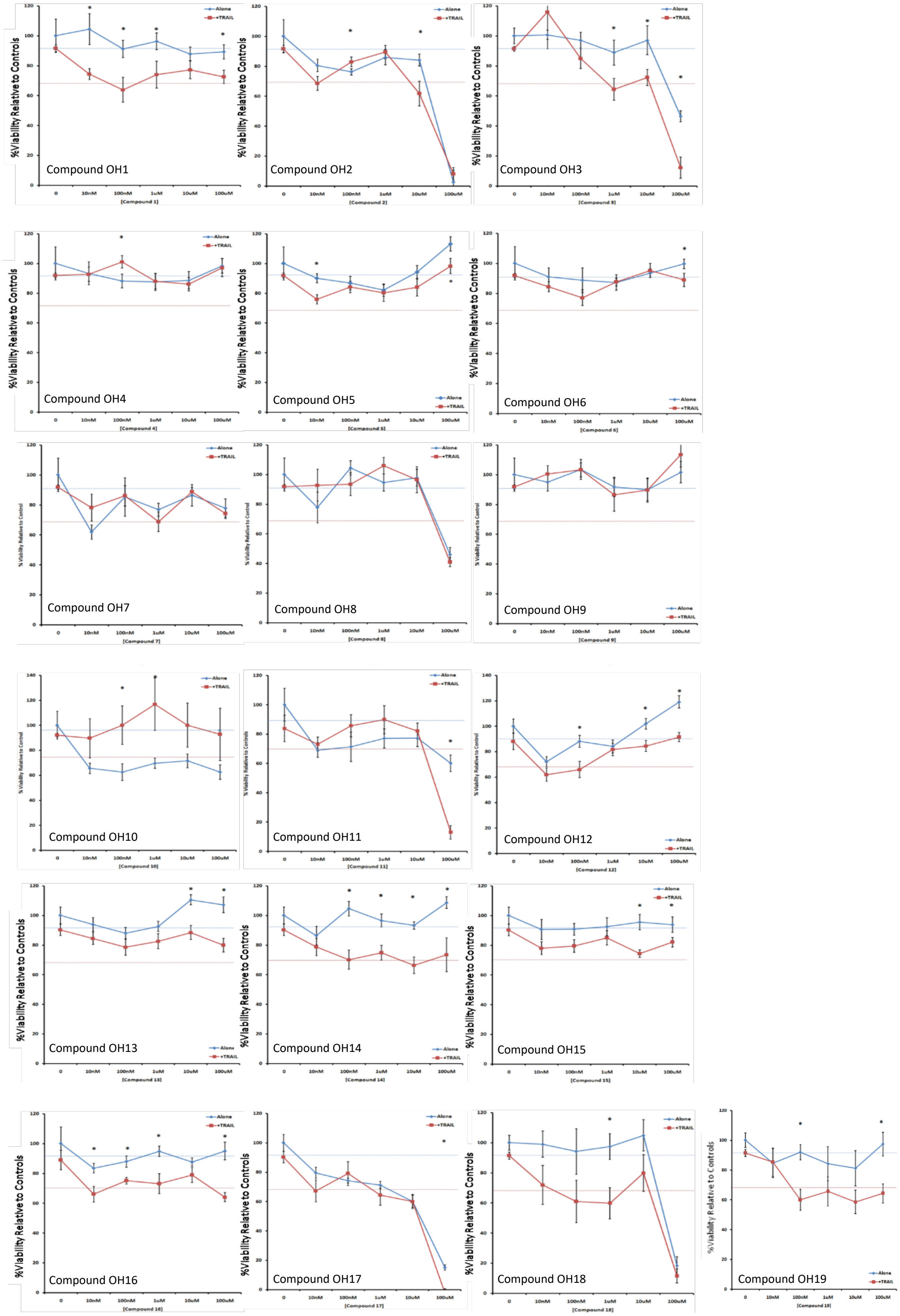
Compound OH1
Compound OH2
Compound OH3
Compound OH4
Compound OH5
Compound OH6
Compound OH7
Compound OH8
Compound OH9
Compound OH10
Compound OH11
Compound OH12
Compound OH13
Compound OH14
Compound OH15
Compound OH16
Compound OH17
Compound OH18
Compound OH19
%Viability Relative to Controls
Alone
+TRAIL

**Supplementary Figure 4:** Viability dose response of OH1-OH19 inhibitor panel on TRAIL resistant MCF-7 cells. The TRAIL resistant ER+ve cell line MCF7 was plated in 96 well plates in 100 µL media in order to achieve 70% confluency, then treated overnight with OH-inhibitor with (red) or without (blue) 20ng/mL killer TRAIL followed by CellTiter blue viability assay. Results represent three independent experiments with four wells per condition per experiment. Blue and red dashed lines (90% and 70% viability respectively) represent the previously published[1,2] effect of c-FLIP knock-down by siRNA and the combination of c-FLIP siRNA with TRAIL treatment respectively under these conditions. * = p-value <0.05. Concentrations of compounds tested were 10 nM, 100 nM, 1 µM, 10 µM and 100 µM.

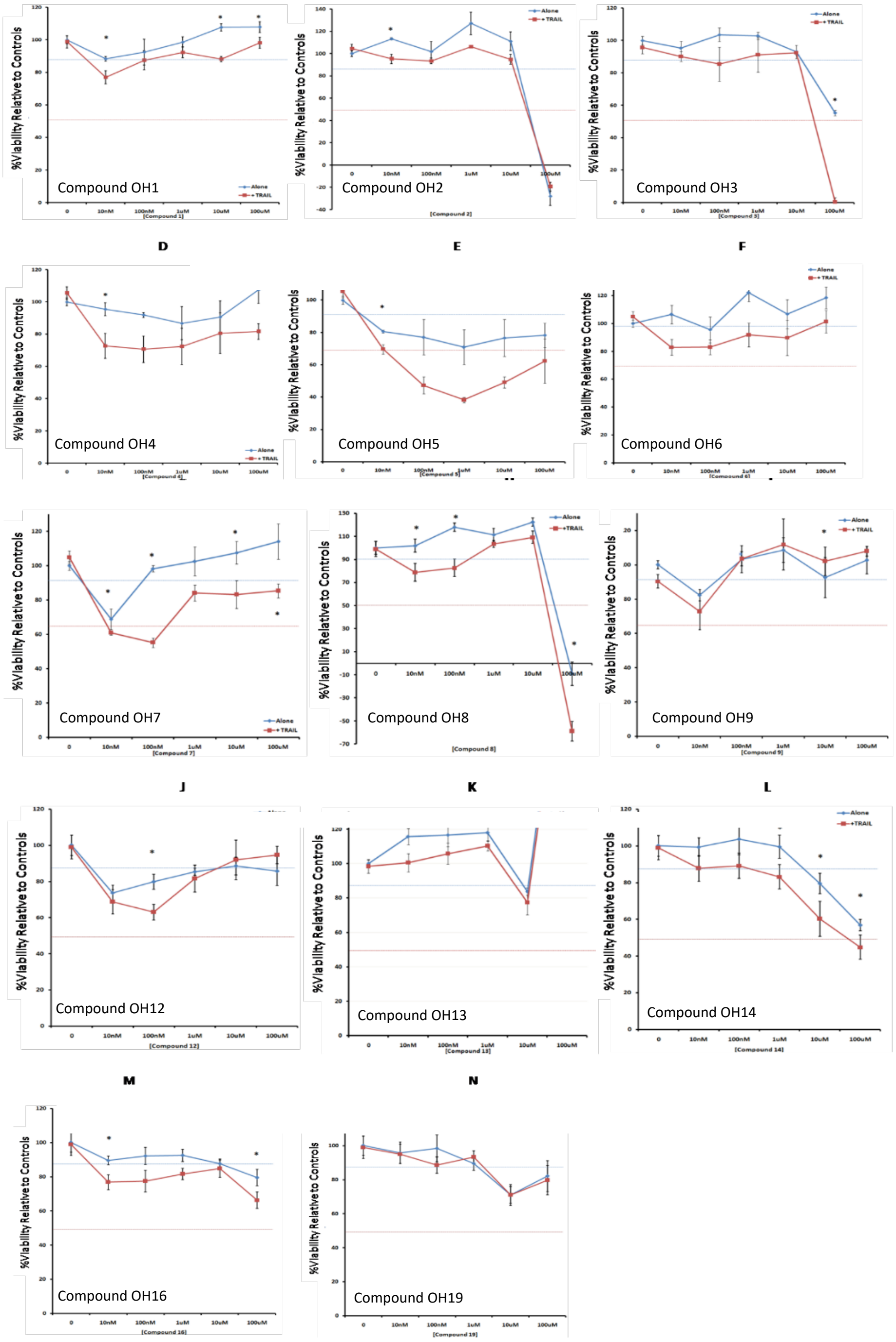


**Supplementary Figure 5:** Viability dose response of OH1-OH19 inhibitor panel on TRAIL resistant BT474 cells. The TRAIL resistant HER2/ER+ve cell line BT474 was plated in 96 well plates in 100 µl media in order to achieve 70% confluency, then treated overnight with selected OH-inhibitor with (red) or without (blue) 20ng/ml killer TRAIL followed by CellTiter blue viability assay. Compounds 10, 11, 15, 17 and 18 were not tested on this cell line due to either their toxicity or lack of efficacy in MCF-7 cells (Supp Fig.4), and their relative insolubility in DMSO. Results represent three independent experiments with four wells per condition per experiment. Blue and red dashed lines (90% and 70% viability respectively) represent the previously published[1,2] effect of c-FLIP knock-down by siRNA and the combination of c-FLIP siRNA with TRAIL treatment respectively under these conditions. * = p-value <0.05. Concentrations of compounds tested were 10 nM, 100 nM, 1 µM, 10 µM and 100 µM. Note that in addition to compound OH14 which showed the most consistent responses of all the compounds in both cell lines, other compounds such as compounds OH1, OH4 and OH7 also exhibited indicative responses in one cell line or in a limited number of doses. While these may be considered as potential candidates they were not progressed in this study.

| Compound | | Metabolic Stability (Species=Human) | | | | | | |
|---|---|---|---|---|---|---|---|---|
| Cyprotex Id | Customer Id | $CL_{int}$ (µL/min/mg protein) | SE $CL_{int}$ | $t_{1/2}$ (min) | n | Comments | Supplier Test Id | Control Group Id |
| CY0000140145 | OH14 | 15.1 | 3.14 | 91.8 | 5 | | 2643664 | 75833 |

**Supplementary Figure 6:** OH14 human microsomal stability data performed by Cyprotex UK Ltd. Microsomes (final protein concentration 0.5 mg/mL) and 0.1 M phosphate buffer were pre-incubated at 37 °C before addition of test compound (final substrate concentration 3 µM, 0.25% DMSO). Samples were mixed and immediately terminated by addition to cold acetonitrile. Samples were centrifuged at 2500 rpm for 30 minutes to precipitate protein. Supernatant was added to equal parts internal standard and samples analysed using Cyprotex generic LC-MS/MS conditions. OH14 had a $CL_{int}$ value of 15.1 µL/min/mg protein, and t1/2 = 91.8 min (n=5).

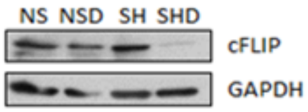


NS = Non-specific control
SH = cFLIP shRNA
D = Doxycycline 48h

**Supplementary Figure 7:** Western blot determination of endogenous cFLIP protein expression in MCF7 cells stably expressing the doxycycline-inducible shRNA lentiviral construct targeting cFLIP. Cells were treated with doxycycline (see methods) or vehicle control for 48 hours prior to cell harvest and analysis of cell lysates by western blot.

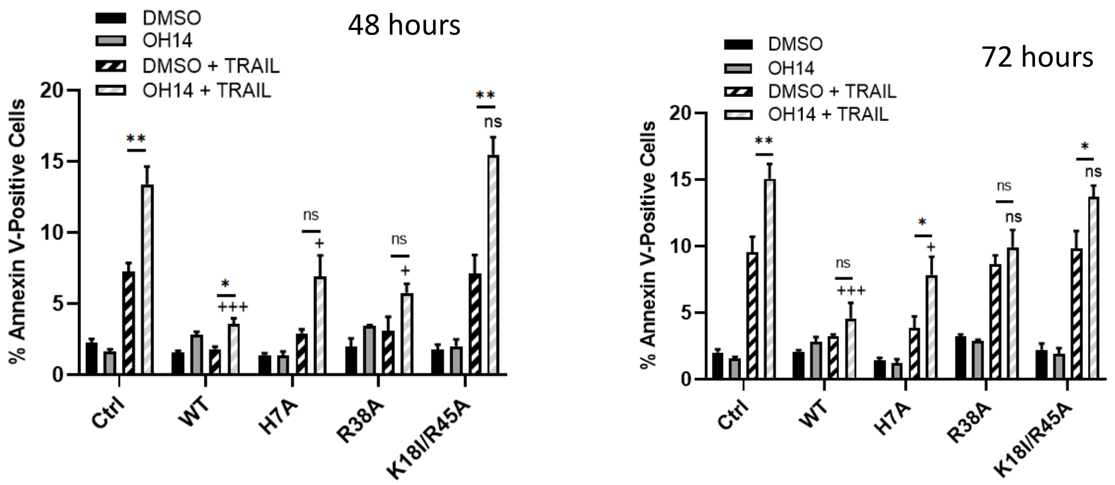


**Supplementary Figure 8:** HeLa cells stably transfected with WT or mutant cFLIPL were treated with 100 mM OH14 followed by 20ng/ml TRAIL for 48 hours or 72 hours and viability assessed by Annexin V staining using Incucyte® real-time analysis. Control is empty vector transfected cells. Data presented as absolute values of % annexin V positive cells. All experiments represent a minimum of three independent replicates. (*p<0.05, **p<0.01, ***p<0.001) indicates statistical significance of OH14 + TRAIL compared to TRAIL alone, + = p<0.05, ++ = p<0.01, +++ = p<0.001 between control OH14 + TRAIL and mutant construct OH14 +TRAIL.

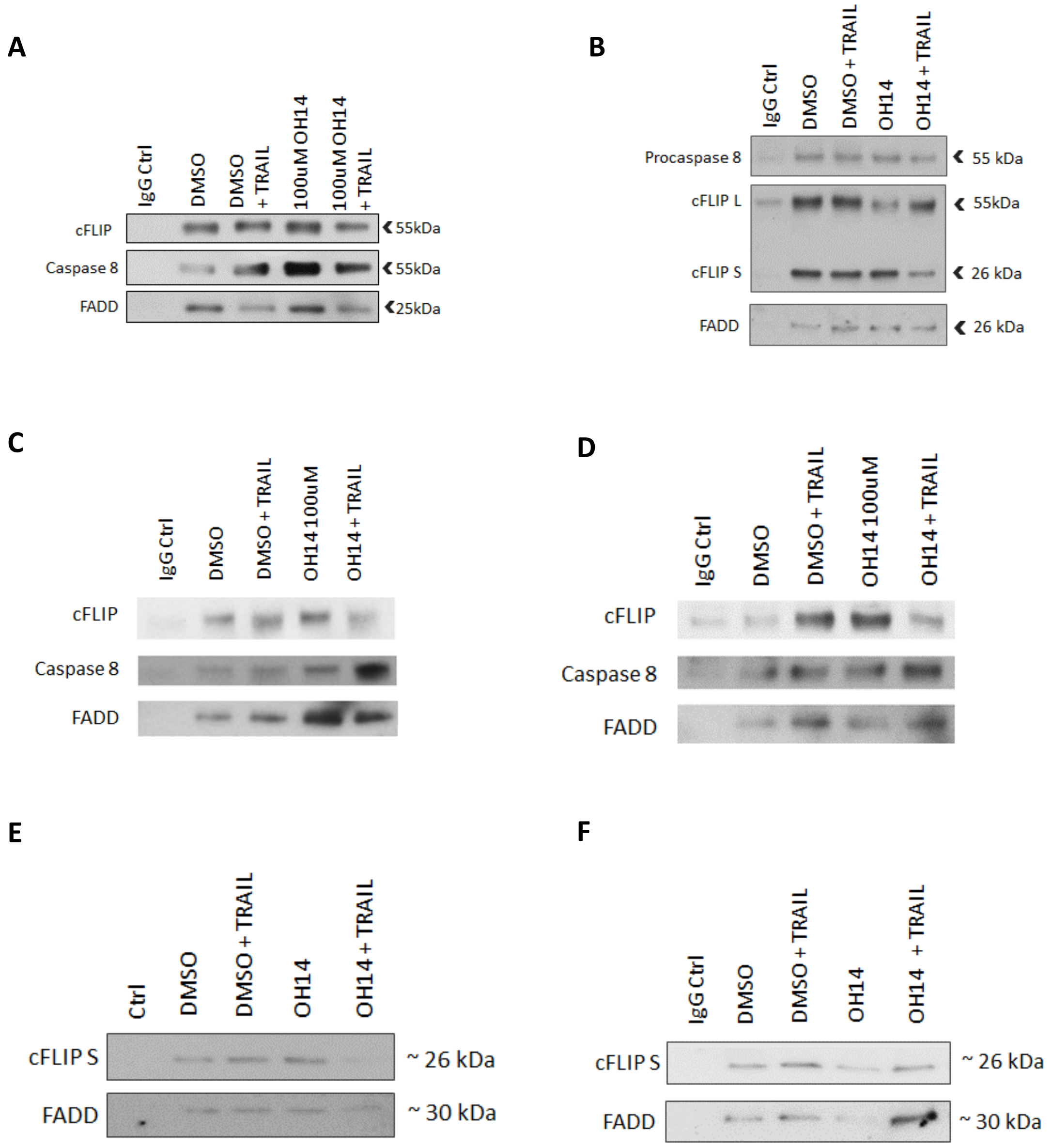


**Supplementary Figure 9:** OH14 impairs cFLIP recruitment to the TRAIL-induced DISC. **A-F:** Western blot analysis of FADD IP in MCF-7 cells pre-treated with 100mM OH14 for 1h followed by 2h of 20ng/ml TRAIL. Each western blot panel represents an independent set of cell cultures, protein extractions and IPs. The normalised band intensities from these panels are depicted in the graph of the data in main **Figure 3A**. An independent IgG control for the FADD IP is present for each western blot. cFLIP or cFLIP L = cFLIP Long isoform (55kDa); cFLIP S = cFLIP Short isoform (26kDa).

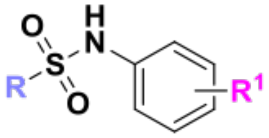


| Product | R | $R^1$ |
|---|---|---|
| **Alkyl(arylsulfonamidobenzoates)** | | |
| **21** | (2,4-dichloro-5-methyl)phenyl | 2-methyl ester |
| **22** | (2,4-dichloro-5-methyl)phenyl | 3-methyl ester |
| **23** | (2,4-dichloro-5-methyl)phenyl | 4-methyl ester |
| **24** | (4-trifluoromethyl)phenyl | 2-methyl ester |
| **25** | (3,4-dimethyl)phenyl | 2-methyl ester |
| **26** | (3,4-dichloro)phenyl | 2-methyl ester |
| **28** | (2,4-dimethyl)phenyl | 2-methyl ester |
| **30** | (2,5-dichloro)phenyl | 2-methyl ester |
| **33** | 1-phenylmethane | 2-methyl ester |
| **34** | 3-pyridine | 2-methyl ester |
| **35** | 2-naphthalene | 2-methyl ester |
| **36** | 1-naphthalene | 2-methyl ester |
| **37** | cyclohexane | 2-methyl ester |
| **42** | (2-methyl-5-chloro)phenyl | 2-methyl ester |
| **43** | (3-chloro-4-methyl)phenyl | 2-methyl ester |
| **50** | (2,4-dichloro-5-methyl)phenyl | 2-ethyl ester |
| **52** | (4-iodo)phenyl | 2-methyl ester |
| **56** | 1-(*p*-tolyl)methane | 2-methyl ester |
| **57** | 2-furan | 2- methyl ester |
| **78** | (2,4-dichloro-5-methyl)phenyl | 2- isopropyl ester |
| **79** | (3-chloro-4-methyl)phenyl | 2-isopropyl ester |
| **80** | (2,4-dichloro-5-methyl)phenyl | 2-*tert*-butyl ester |
| **(Arylsulfonamido)benzoic acids** | | |
| **64** | 1-(4-(trifluoromethyl)phenyl)methane | 2-carboxylic acid |
| **65** | 2-thiophene | 2-carboxylic acid |
| **66** | 8-quinoline | 2-carboxylic acid |

| | | |
|---|---|---|
| **67** | 3-quinoline | 2-carboxylic acid |
| **72** | 3-pyridine | 2-carboxylic acid |
| **73** | 2-napthalene | 2-carboxylic acid |
| **74** | 1-naphtalene | 2-carboxylic acid |
| **75** | cyclohexane | 2-carboxylic acid |
| **76** | 2-furan | 2-carboxylic acid |
| **(Arylsulfonamido)N-(1H-tetrazol-5-yl)phenyls** | | |
| **88** | (2,4-dichloro-5-methyl)phenyl | 2-(1*H*-tetrazol) |
| **89** | (3-chloro-4-methyl)phenyl | 2-(1*H*-tetrazol) |
| **90** | (4-trifluoromethyl)phenyl | 2-(1*H*-tetrazol) |
| **91** | 2-naphtalene | 2-(1*H*-tetrazol) |
| **92** | 8-quinoline | 2-(1*H*-tetrazol) |
| **93** | 2-quinoline | 2-(1*H*-tetrazol) |
| **Alkyl(arylsulfonamido)benzamides** | | |
| **101** | (3-chloro-4-methyl)phenyl | 2-N-methyl carboxamide |
| **102** | (2,4-dichloro-5-methyl)phenyl | 2-N-methyl carboxamide |
| **103** | (2,4-dichloro-5-methyl)phenyl | 2-N-methyl carboxamide |
| **104** | (2,4-dichloro-5-methyl)phenyl | 2-N,N-dimethyl carboxamide |
| **105** | (2,4-dichloro-5-methyl)phenyl | 2-N-isopropyl carboxamide |
| **106** | (2,4-dichloro-5-methyl)phenyl | 2-carboxamide |

**Supplementary Table 1:** Identity of arylsulfonamide derivatives for SAR study.

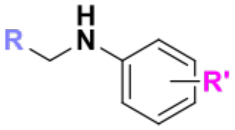


| Molecule | R | R' |
|---|---|---|
| **(Arylamino)benzoic acids** | | |
| **113** | (4-trifluoromethyl)phenyl | 2-carboxylic acid |
| **115** | (3,4-dimethyl)phenyl | 2-carboxylic acid |
| **117** | (3-chloro-4-methyl)phenyl | 2-carboxylic acid |
| **119** | 8-quinoline | 2-carboxylic acid |
| **120** | 2-quinoline | 2-carboxylic acid |
| **139** | 3-pyridine | 2-carboxylic acid |
| **141** | 2-naphtalene | 2-carboxylic acid |
| **142** | 1-naphtalene | 2-carboxylic acid |
| **143** | (4-trifluoromethyl)benzyl | 2-carboxylic acid |
| **156** | 5-chloro-2-methyl | 2-methylbenzoic acid |
| **Methyl(arylamino)benzoates** | | |
| **121** | (4-trifluoromethyl)phenyl | 2-methyl ester |
| **123** | (3,4-dimethyl)phenyl | 2-methyl ester |
| **129** | (3-chloro-4-methyl)phenyl | 2-methyl ester |
| **131** | 3-pyridine | 2-methyl ester |
| **133** | 2-naphtalene | 2-methyl ester |
| **134** | 1-naphtalene | 2-methyl ester |
| **137** | benzyl | 2-methyl ester |
| **138** | (4-trifluoromethyl)benzyl | 2-methyl ester |
| **(Arylamino)N-(1H-tetrazol-5-yl)phenyl** | | |
| **144** | (4-trifluoromethyl)phenyl | 2-(1H-tetrazole) |
| **145** | (3-chloro-4-methyl)phenyl | 2-(1H-tetrazole) |
| **147** | 8-quinoline | 2-(1H-tetrazole) |
| **148** | 2-quinoline | 2-(1H-tetrazole) |
| **149** | 2-naphtalene | 2-(1H-tetrazole) |
| **(Arylamino)N-(5-oxo-2,5-dihydro-1,2,4-oxadiazol-3-yl)phenyl** | | |

| | | |
|---|---|---|
| **150** | (3-chloro-4-methyl)phenyl | 2-(5-oxo-4,5-dihydro-1,2,4-oxadiazol) |

**Supplementary Table 2:** Identity of amine-linked derivatives for SAR study.

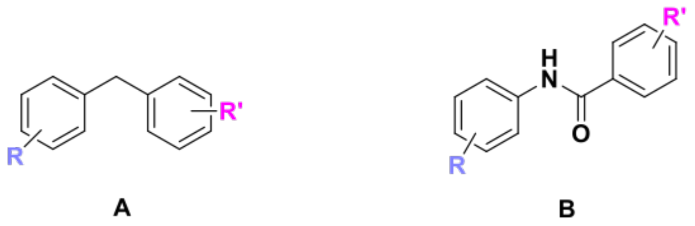


| Molecule | Linker | R | R’ |
|---|---|---|---|
| **169** | Methylene (**A**) | 4-trifluoromethyl | 2-methyl ester |
| **173** | Methylene (**A**) | 3-chloro-4-methyl | 2-carboxylic acid |
| **179** | Amide (**B**) | 4-trifluoromethyl | 2-methyl ester |
| **182** | Amide (**B**) | 3-chloro-2-methyl | 2-methyl ester |
| **186** | Amide (**B**) | 3-chloro-4-methyl | 2-carboxylic acid |

**Table 3:** Identity of methylene- and amide-linked derivatives.

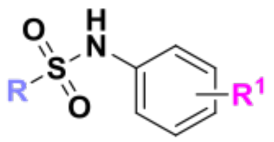

| Starting Sulfonyl Chloride | $R^1$ | R | Product | Yield% |
|---|---|---|---|---|
| **1** | (2,4-dichloro-5-methyl)phenyl | **18** | **21** | 49% |
| **1** | (2,4-dichloro-5-methyl)phenyl | **19** | **22** | 56% |
| **1** | (2,4-dichloro-5-methyl)phenyl | **20** | **23** | 46% |
| **4** | (4-trifluoromethyl)phenyl | **18** | **24** | 60% |
| **5** | (3,4-dimethyl)phenyl | **18** | **25** | 29% |
| **6** | (3,4-dichloro)phenyl | **18** | **26** | 65% |
| **8** | (2,4-dimethyl)phenyl | **18** | **28** | 55% |
| **10** | (2,5-dichloro)phenyl | **18** | **30** | 56% |
| **13** | 1-phenylmethane | **18** | **33** | 28% |
| **14** | 3-pyridine | **18** | **34** | 79% |
| **15** | 2-napthalene | **18** | **35** | 58% |
| **16** | 1-napthalene | **18** | **36** | 48% |
| **17** | Cyclohexane | **18** | **37** | 26% |
| **40** | 5-chloro-2-methylphenyl | **18** | **42** | 9% |
| **41** | 3-chloro-4-methylphenyl | **18** | **43** | 24% |
| **1** | (2,4-dichloro-5-methyl)phenyl | **44** | **50** | 43% |
| **46** | (4-iodo)phenyl | **18** | **52** | 25% |
| **48** | 1-(*p*-tolyl)methane | **18** | **56** | 15% |
| **49** | 2-furan | **18** | **57** | 57% |
| **59** | 1-(4-(trifluoromethyl)phenyl)methane | **2-COOH** | **64** | 23% |
| **60** | 2-thiophene | **2-COOH** | **65** | 22% |
| **61** | 8-quinoline | **2-COOH** | **66** | 28% |
| **62** | 3-quinoline | **2-COOH** | **67** | 35% |
| **Starting Ester** | | | | |
| **34** | 3-pyridine | **2-COOMe** | **72** | 30% |
| **35** | 2-napthalene | **2-COOMe** | **73** | 67% |
| **36** | 1-napthalene | **2-COOMe** | **74** | 90% |

| | | | | |
|---|---|---|---|---|
| **37** | cyclohexane | **2-COOMe** | **75** | 26% |
| **38** | 2-furan | **2-COOMe** | **76** | 32% |
| **(1H-tetrazol-5yl)aniline** | | | | |
| **84** | (2,4-dichloro-5-methyl)phenyl (**1**) | **-** | **88** | 18% |
| **84** | 3-chloro-4-methyl (**41**) | **-** | **89** | 16% |
| **84** | (4-trifluoromethyl)phenyl (**4**) | **-** | **90** | 42% |
| **84** | 2-napthalene (**15**) | **-** | **91** | 38% |
| **84** | 8-quinoline (**62**) | **-** | **92** | 26% |
| **84** | 3-quinoline (**63**) | **-** | **93** | 46% |
| **Starting carboxylic acid** | | | | |
| **69** | 3-chloro-4-methyl | 2-CONHMe | **101** | 30% |
| **3** | 2,4-dichloro-5-methyl | 2-CONHMe | **102** | 49% |
| **3** | 2,4-dichloro-5-methyl | 2-CONHEt | **103** | 44% |
| **3** | 2,4-dichloro-5-methyl | 2-CON(Me)$_2$ | **104** | 35% |
| **3** | 2,4-dichloro-5-methyl | 2-CON(Me)$_2$ | **105** | 53% |

**Supplementary Table 4:** Isolated yields and identities of alkyl(arylsulfonamidobenzoates)

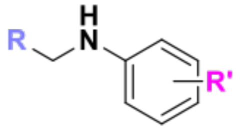


| Starting aldehyde | R | | Product | Yield |
|---|---|---|---|---|
| **107** | 4-(trifluoromethyl)phenyl | 2-COOH | **113** | 17% |
| **108** | (3,4-dimethyl)phenyl | 2-COOH | **115** | 52% |
| **109** | (3-chloro-4-methyl)phenyl | 2-COOH | **117** | 21% |
| **110** | 8-quinoline | 2-COOH | **119** | 43% |
| **111** | 3-quinoline | 2-COOH | **120** | 41% |
| **109** | (3-chloro-4-methyl)phenyl | 2-COOMe | **129** | 26% |
| **125** | 3-pyridine | 2-COOMe | **131** | 96% |
| **127** | 2-napthalene | 2-COOMe | **133** | 76% |
| **128** | 1-napthalene | 2-COOMe | **134** | 79% |
| **131** | 3-pyridine | 2-COOH | **139** | 28% |
| **132** | 4-pyridine | 2-COOH | **140** | 26% |
| **133** | 2-napthalene | 2-COOH | **141** | 89% |
| **134** | 1-napthalene | 2-COOH | **142** | 41% |
| **136** | (4-trifluoromethyl)benzyl | 2-COOH | **143** | 16% |
| **107** | 4-(trifluoromethyl)phenyl | 2-(1H-tetrazole) | **144** | 32% |
| **109** | (3-chloro-4-methyl)phenyl | 2-(1H-tetrazole) | **145** | 25% |
| **109** | (3-chloro-4-methyl)phenyl | 2-(1H-tetrazole)) | **146** | 15% |
| **110** | 8-quinoline | 2-(1H-tetrazole) | **147** | 12% |
| **111** | 2-quinoline | 2-(1H-tetrazole) | **148** | 11% |
| **127** | 2-napthalene | 2-(1H-tetrazole) | **149** | 75% |
| **Starting bromide** | | | | |
| **135** | benzyl | 2-COOMe | **137** | 5% |
| **136** | 4-(trifluoromethyl)phenyl | 2-COOMe | **138** | 26% |
| **Starting Compound** | | | | |
| **113** | 4-trifluoromethyl | 2-COOMe | **121** | 44% |
| **116** | 3,4-dimethyl | 2-COOMe | **123** | 13% |

**Supplementary Table 5:** Isolated yields and identities of amine-linked derivatives.

**General Methods.**

All experiments were performed with the approval of the Cardiff University School of Biosciences Ethics Committee.

Constructs

The pCMV2FLAG_cFLIPL overexpression vector, containing the full-length coding sequence of the long form of human c-FLIP (accession number NM_003879.4) and FLAG tag on the C-terminal was purchased from SinoBiological (HG11110-M-F), together with pCMV-FLAG control (CV005). The pTRIPz cFLIP (cFLAR) and nonspecific control inducible shRNA plasmids were purchased from Dharmacon/Horizon. FADD and cFLIP were cloned into e-CFP-C1 or e-YFP-C1 FRET plasmids respectively; both constructs were gifts from Dr Ladislav Andera, Institute of Molecular Genetics, Prague. All oligonucleotides were custom-designed and purchased from Sigma and all cloning reagents were purchased from New England Biolabs. Cells were transformed with constructs using lipofectamine 3000 (Invitrogen) according to the manufacturer's instructions or transduced with lentivral particles in the case of the pTRIPz shRNA vectors.

Site Directed Mutagenesis

Site directed mutagenesis was performed on the pCMV-cFLIPL-FLAG construct, using the QuickChange kit (Stratagene) according to the manufacturer's instructions to introduce the following mutations; H7A, R38A and K18I/R45A. The following mutagenic primers were used:
H7A: 5'-GTCTGCTGAAGTCATC**GC**TCAGGTTGAAGAAGCAC- 3'
R38A: 5' –GTGGTTCCACCTAATGTC**GC**GGACCTTCTGGATATTTTAC- 3'
K18I: 5'-CTTGATACAGATGAGA**TC**GAGATGCTGCTCTTTTTGTG-3'
R45A: 5'-CCTTCTGGATATTTTA**GC**GGAAAGAGGTAAGC-3'

Cell Lines

The human breast cancer cell lines MDA-MB-231$^{ER-HER2-}$ HCC1954$^{HER2+}$ and BT474 $^{ER+HER2+}$ were obtained from ATCC and tested for mycoplasma at least 3 passages before use . The MCF-7$^{ER+}$ cell lines were a gift from Dr Julia Gee, Cardiff University. The HeLa human cervical cancer cell line was a gift from Dr Ladislav Andera, Institute of Molecular Genetics, Prague. The primary-derived breast cancer cell lines were a gift from Dr Rob Clarke, University of Manchester. The SUM149 $^{ER-HER2-}$ line was purchased from Asterand Bioscience (Detroit, USA). All cell lines except SUM149 were cultured in RPMI 1640 medium (Invitrogen) supplemented with 10% foetal bovine serum (FBS) (Invitrogen), and 1% penicillin-streptomycin and L-glutamine mix (Invitrogen). The SUM 149 cell line was cultured in Hams F12 media (Invitrogen, Paisley, UK) supplemented with 5% fetal bovine serum (Sigma), 2mM L- glutamine (Invitrogen), 10mM HEPES (Invitrogen), 1µg/ml Hydrocortisone (Invitrogen) and 5µg/ml insulin (Invitrogen). All cell lines were cultured at 37℃ in 5% $CO_2$.

Reagents

Recombinant soluble human TRAIL was purchased as super-killer TRAIL from Enzo Life Sciences. Unless otherwise stated, cells were treated with 20 ng/ml TRAIL without cross-linking for 18 hours before subsequent assays. The pan-caspase inhibitor z-vad-fmk was purchased from R and D systems and used at a concentration of 20 µM. Cells were pre-treated with caspase inhibitor for 1 h prior to treatment with TRAIL.

<u>CellTitre-blue Assay</u>

Cells to be analysed were cultured in a 96-well plate format. On the day of analysis, 20 ⍰l of CellTiter-Blue reagent (Promega) was added to each well containing 100⍰l media. The plate was incubated for 1-4 h at 37℃ in 5% $CO_2$, before fluorescence was measured at 560/590 nm using a FLUOstar Optima plate reader (BMG Labtech, Offenberg, Germany).

<u>Incucyte Viability Assays</u>

Cells to be analysed were cultured in a 96-well plate format, each well containing 100 ⍰l media. Following treatment with TRAIL and/or OH14, Annexin V reagent (EssenBioscience) or Caspase-3/7 Green Dye for IncuCyte (1:1000) (Sartorius) was added to each well according to the manufacturer's instructions. The plate was incubated for up to 72 h at 37℃ / 5% $CO_2$ in the Incucyte live cell imager (EssenBioscience) where fluorescence was measured at 2 h intervals. MCF-7 pTRIPZ-shFLIP cells (which overexpress a doxycycline-inducible shRNA targeting cFLIP) were selected and maintained in RPMI 1640 media supplemented with 10% FBS and 2ug/mL puromycin. Cells were seeded at 5k cells/well (50k/mL, 100uL per well) into black-walled clear-bottom 96-well plates (Greiner) and incubated overnight. Cells were then treated with 2ug/mL doxycycline the following day for 48 hours. Cells were visually inspected under a fluorescence microscope for red fluorescence to verify doxycycline induced knockdown of c-FLIP. Cells were then treated with 75microM OH14 and Caspase-3/7 Green Dye, followed by 20ng/mL TRAIL after 1 hour. Cells were monitored and imaged in the IncuCyte live-cell imaging system for 48 hours, where images were acquired every 2 hours in the phase and green fluorescence channels.

<u>Colony Forming Assay</u>

Cells were seeded at a density of 185 cells/well in a 12-well plate format, so that cells were 50 per square cm, and cultured for 10 days[17]. To stain colonies, culture medium was removed and well surface was washed once with PBS. Crystal violet/ethanol mixture was applied to wells and incubated for 15 mins at room temperature. Solution was removed and wells were rinsed twice with PBS. Colonies containing approximately 32 or more cells (having undergone 5 or more divisions) were counted using a GelCount platereader and software (Oxford Optronix) set to count colonies of diameters 100-1000 ⍰m.

<u>Western Blotting</u>

Total cellular or cytoplasmic proteins were extracted from cultured cells and analysed by Western blotting. FADD antibody was purchased from Cell Signalling Technologies (mouse anti-FADD 2782), cFLIP antibodies used were purchased from Santa Cruz (5D8, sc136160) and Enzo Life Sciences (7F10, ALX-804-961-0100). Caspase-8 antibody was purchased from Cell Signalling Technologies (9746). To quantitate Western data, the pixel intensity of each band was quantified relative to its protein loading control (GAPDH, Santa Cruz, sc32233) by densitometry using the program ImageJ (http://imagej.nih.gov/ij/).

<u>Immunoprecipitation</u>

Adherent cells to be analysed were washed twice with ice cold PBS and harvested by scraping in pre-chilled lysis buffer containing protease inhibitors and incubated on ice for 30 mins. Insoluble material was removed by centrifugation at 13,000rpm for 10 min at 4°C. Cell lysates were pre-cleared to remove non-specific interaction proteins by adding protein A-

Sepharose suspension (17-0780-01–GE GE Healthcare, Sigma) to lysate at a ratio of 1:10 and incubating at 4°C with agitation for 1 h. Beads were removed by centrifugation then 0.4µg/ml of FADD antibody (rabbit anti-FADD H-181 SantaCruz sc-5559) or nonspecific isotype control (rabbit IgG, abcam ab 172730) was added to the cleared lysate and incubated overnight at 4°C with agitation. Protein A-Sepharose suspension was added to lysate/antibody solution at a ratio of 1:10 and incubated at 4 °C, with agitation for 2 h. Beads were collected by centrifugation at 13,000 rpm, 4°C, for 2 min and washed in lysis buffer. A total of 5 washes were performed, after which the required volume of lysis buffer and 5x Laemmli buffer was added directly to the beads, mixed, and heated for 2 min at 95C. The immunoprecipitated material was loaded directly onto an SDS gel using a Hamilton syringe for analysis by Western Blotting.

FRET

The TRAIL resistant MCF-7 or HeLa cells were transiently transfected with e-CFP-C1-FADD or e-YFP-C1-cFLIPL FRET constructs using Lipofectamine 3000 (Life Technologies). Pre-treatment with a pan-caspase inhibitor, z-vad-fmk (R and D systems) for 1 h before transfection was used for all FRET transfection experiments to prevent cell death during analysis, and cells treated with TRAIL and/or OH14 24h post-transfection. Measurements were performed on a Zeiss LSM 710 confocal microscope (Zeiss, Germany). Viable cells with similar CFP and YFP intensity were selected for analysis. At least 10 cells were recorded per experimental condition and exposure times were kept between 100-500 ms. Spectra were recorded using a 458/514 nm double dichroic excitation to facilitate excitation of CFP with the 458 nm laser and YFP with the 514 nm laser line. Image acquisition was set to obtain 5 intensity recordings pre-bleach and 60 seconds post-bleach (approximately 40 recordings). For intensity measurements, a HXP120V to Arc 100W mercury lamp (Zeiss, Germany) was used. Images were recorded on the Zeiss LSM 710 capture camera and LASOS Argon Laser with 458/488/514 nm band excitation filters was used for CFP and YFP imaging and photobleaching. All FRET measurements were performed at 37°C and 5% $CO_2$.

Molecular Modelling

The structures of the two DEDs of cFLIP and procaspase-8 were generated using MOE alignment and homology tool (Molecular Operative Environment, 2010) [Molecular Operating Environment (MOE) Integrated Computer-Aided Molecular Design Platform (Chemical Computing Group); www.chemcomp.com/Products.htm] using the structure of MC159 as template (PDB ID: 2BBR)[18]. MC159 was prepared in MOE where hydrogen atoms were added, residue ionization was set to pH 7.4 and the structure was minimized to keep all heavy atoms fixed until the RMSD gradient of 0.05 kcal mol$^{-1}$ Å$^{-1}$was obtained.

The known intramolecular interactions between the internal 'FL motif' on DED1 and a pocket on the internal surface of DED2 of cFLIP/procaspase-8[14] were used to model the intermolecular interactions between these proteins, with the FL motif of DED2 binding to a pocket on DED1. The intermolecular interaction between procaspase-8 and c-FLIP was then used to model the cFLIP:FADD and procaspase-8:FADD interactions (FADD crystal structure PDB ID: 1A1W)[19]. The resulting protein complexes were energy minimised in MOE, using the Amber 99 forcefield to ensure the gradient of 0.05 kcal mol-1 Å-1 was maintained. Following this, GROMACS software[20]. was used to perform molecular dynamics on the protein complexes using Gromos 96 forcefield and NPT working environment (300K, v-rescale; 1 atm, Berendsen), with a timestep of 0.002 ps. Initially, the system was solvated in a cubic box using spc216 water molecules, generating a minimum of 9 Å water between the protein and the edge of the box, and then it was energy minimised. Next, the whole system was initially equilibrated for 100ps, then the MD simulation was run for 15 ns. All systems reached stability within the first 3ns, but for analysis purpose, only the last 10ns of the simulation were considered. Where appropriate, the final frame of the simulation was then energy minimised and used in the virtual screening protocol (pharmacophore and docking). MD trajectories were visualised in VMD (Visual Molecular Dynamics).

A pharmacophore query was created based on the cFLIP:FADD interaction, using MOE. The FL motif of FADD was used as a guide to set up the query. The query was then used to filter the SPECS library, prepared with the import conformation tool of MOE, and the resulting hit compounds were docked independently on cFLIP and procaspase-8 using Glide [Schrodinger: Glide. 5.5 Edition. New York, NY (2009)], using the SP approach and rescoring the result with the XP method. The top 200 compounds that scored highly for cFLIP binding and low for procaspase-8 binding were visually examined, leading to a final selection of 19 compounds.

Statistical Analysis

Throughout the article, data are represented as mean and error bars as standard error of a minimum of three independent experiments, unless otherwise stated. Statistical significance was determined using a student's T-test for unpaired samples where direct comparisons were required between two conditions assuming non-normal distribution and similar variance (all assays) or one-way Anova with one independent variable and multiple conditions (FRET data). Key for statistical cut-offs on all graphs: * = $p<0.05$, ** = $p<0.01$, *** = $p<0.001$. L-Calc software was used to estimate stem cell number from serial dilutions of tumour xenografts: (http://www.stemcell.com/en/Products/All-Products/LCalc-Software.aspx)

**Chemistry experimental.**

All chemicals, reagents and solvents were purchased from commercial sources, and were used without further purification. TLC was performed on silica gel plates (Merck Kieselgel 60F254) and was developed by the ascending method. After solvent evaporation, compounds were visualised by irradiation with UV light at 254 nm and 366 nm. Flash column chromatography was performed using the Biotage Isolera One system. 1H and 13C NMR spectra were recorded on a Bruker AVANCE DPX500 spectrometer (500 MHz and 75 MHz respectively) and auto-calibrated to the deuterated solvent reference peak. Chemical shifts are given in [illegible] relative to tetramethylsilane (TMS); the coupling constants (J) are given in Hertz. TMS was used as an internal standard ([illegible] = 0) for 1H- NMR and CDCl3 served as an internal standard ([illegible] = 77.0) for 13C- NMR. Purity and mass of compounds were determined by UPLC-MS analyses, performed on a Waters UPLC system provided with Diode Array detector and Electrospray (positive and negative ion) MS detector. Stationary phase: Waters Acquity UPLC BEH C18 1.7 um 2.1 × 50 mm column. Mobile phase: H2O containing 0.1% Formic acid (A) and MeCN containing 0.1% Formic acid (B). Column temperature: 40 °C. Samples were prepared in acetonitrile at 1 µg/mL concentration. Injection volume: 2 µL. A linear gradient standard method (A) was used: 90% A (0.1 min), 90–0% A (2.6 min), 0% A (0.3 min), 90% A (0.1 min); flow rate 0.5 mL/min. Purity of the compounds tested in biological assays was >95%.

**Preparation and characterisation of synthetic intermediates.**

**Synthesis of intermediate isopropyl 2-aminobenzoate (77)**

To a solution of 2-aminobenzoic acid (1.5 g, 10.93mmol, 1 eq) in isopropanol (1.9 mL/mmol) was added 1.25 M HCl in isopropanol solution (1.9 mL/mmol). The reaction was refluxed overnight. The solution was then cooled to room temperature and concentrated in vacuum. The residue was partitioned between sat. aq. NaHCO3 solution (30mL/mmol) and ethylacetate (3x25 mL/mmol). The combined organic layers were washed with sat. aq. NaHCO3 solution (25 mL/mmol) and water (10 mL/mmol), dried over MgSO4 and concentrated under vacuum. The desired products was purified by flash column chromatography (*n*-hexane:EtOAc 100:0 v/v increasing to *n*hexane:EtOAc 9:1 v/v). 77 was obtained as yellow oil in 19% yield. $^{1}$H-NMR ($CDCl_3$), δ: 1.28 (d, J= 6.2 Hz, 6H, H-9,10), 5.14 (m, 1H, H-8), 5.63 (bs, 2H, NH2), 6.54-5.58
(m, 2H, H-aromatic), 7.15-7.19 (m, 1H, H-aromatic), 7.79 (dd, J1= 7.8 Hz, J2= 1.5 Hz, 1H, H-aromatic).$^{13}$C-NMR ($CDCl_3$), δ: 22.0 (2xCH3, C-9,10), 67.6 (CH, C-8), 116.2, 116.6, 131.2, 133.8 (CH, C-aromatic), 111.5, 150.4 (C, C-aromatic), 167.7 (C, C-7).

**Synthesis of intermediate 2-(1*H*-Tetrazol-5-yl)aniline (84)**

The 2-aminobenzonitrile (1 g, 8.46 mmol, 1 eq), sodium azide (1.2 eq), ammonium chloride (1.2 eq) and DMF (0.6 mL/mmol) were mixed and heated at 120° C for 72 hours. The solvent was evaporated under reduced pressure. The desired product was purified by flash column chromatography (Biotage Isolera One automated flash column chromatography, cartridge: SNAP KP 50g, *n*-hexane-EtOAc 100:0 increasing to *n*-hexane-EtOAc 0:100 in 10CV). 84 was obtained as yellow solid in 53% yield. **$^{1}$H-NMR (DMSO-$D_6$), δ**: 6.67-6.70 (m, 1H, H-aromatic), 6.88-6.92 (m, 1H, H-aromatic), 7.22-7.26 (m, 1H, H-aromatic), 7.72 (dd, J1= 7.8 Hz, J2= 1.3 Hz, 1H, H-aromatic), 9.66 (bs, 1H, NH). **$^{13}$C-NMR (DMSO-$D_6$), δ**: 116.0, 116.8, 128.5, 132.3 (CH, C-aromatic), 105.0, 147.9 (C, C-aromatic), 155.3 (C, C-7).

**Synthesis of methyl 2-(((5-chloro-2-methylphenyl)amino)methyl)benzoate (163)**

A mixture of methyl 2-(bromomethyl)benzoate (**161**) (0.2 g, 0.87 mmol, 1 eq), and 5-chloro-2-methylaniline (**153**) (0.062 g, 0.43 mmol, 1 eq) and TEA (1.5 eq) in methanol (1 mL/mmol) was heated at 85°C. overnight. The mixture was diluted with DCM,washed with saturated solution of NaHCO3 (10 mL/mmol), dried over sodium sulfate and concentrated in vacuum. The crude products were purified flash column chromatography (Biotage Isolera One automated flash column chromatography,cartridge: SNAP KP SIL 25g, *n*-hexane-EtOAc 100:0 increasing to *n*-hexane-EtOAc 80:20 in 15CV). **163** was obtained as white solid in 38% yield. **$^1$H-NMR ($CDCl_3$), δ**: 2.03 (s, 3H, H-16), 3.83 (s, 3H, H-8), 4.39 (bs, 1H, NH), 4.61 (d, J= 6.1 Hz, 2H, H-9), 6.48 (d, J= 2.0 Hz, 1H, H-15), 6.51 (dd, J1= 7.8 Hz, J2=2.0 Hz, 1H, H-13), 6.86 (dd, J1= 7.8 HZ, J2=0.7 Hz, 1H, H-aromatic), 7.26-7.29 (m, 1H, H-aromatic), 7.40-7.42 (m, 2H, H-aromatic), 7.90-7.92 (m, 1H, H-aromatic). **UPLC-MS: Rt** 2.49 (97%) **MS (ESI)$^+$:** 291.8 $[M+H]^+$.

**Synthesis of: methyl 2-(bromomethyl) benzoate (161)**

To a solution of the starting methyl 2-methylbenzoate (1.0 g, 6.65 mmol) in 7 mL of CHCl3, NBS (1.3 g, 7.31 mmol) and AIBN (0.013 g, 0.073 mmol) were added and the reaction was stirred at 70°C for 5 hours. The reaction was then cooled down to room temperature and the deposit of succinimide was filtered. The solvent was evaporated under reduced pressure. The crude product was then purified by flash column chromatography to obtain the pure product (Biotage Isolera One automated flash column chromatography, cartridge: SNAP KP SIL 100g, *n*-hexane-EtOAc 100:0 increasing to *n*-hexane-EtOAc 80:20 in 15CV). **161** was obtained as yellow oil in 82% yield. $^1$H-NMR ($CDCl_3$), δ: 3.95 (s, 3H, H-9), 4.98 (s, 2H, H-7), 7.38-7.40 (m, 1H, H-aromatic), 7.46-7.50 (m,2H, H-aromatic), 7.98-8.00 (m, 1H, H-aromatic).

**Synthesis of: methyl 2-(3-chloro-4-methylbenzyl)benzoate (171)**

To a solution of (3-chloro-4-methylphenyl)boronic acid (0.111 g, 0.94 mmol) and (2-bromomethyl)benzoate (0.150 g, 0.65 mmol) in EtOH (1.5 mL), water (0.4 mL) and toluene (1.2 mL) was added Na2CO3 1M (0.075 g, 0.72 mmol) and catalytic amount of Pd(PPh3)4. The reaction was stirred at 80°C for 2 days. The solution was then filtered through celite and the solvent was evaporated. The residue diluted with water and extracted with diethyl ether (3x10 mL). The organic layer was dried over MgSO4 and evaporated under reduced pressure. The crude product was used for the next step without further purification. **171** was obtained as colourless oil in 56% yield. **$^1$H-NMR ($CDCl_3$), δ**: 2.24 (s, 3H, H-7), 3.76 (s, 3H, H-16), 4.24 (s, 2H), 6.85-6.87 (m, 1H, H-aromatic), 7.03 (d, J= 7.9 Hz, 1H, H-aromatic), 7.04-7.06 (m, 1H, H-aromatic), 7.13 (d, J= 7.9 Hz, 1H, H-aromatic), 7.20-7.24 (m, 1H, H-aromatic), 7.35-7.38 (m, 1H, H-aromatic), 7.83 (dd, J1= 7.8 Hz, J2= 1.3 Hz, 1H, H-aromatic). **$^{13}$C-NMR ($CDCl_3$), δ:** 19.6 (CH3), 51.8 (CH3), 38.9 (CH2), 126.7, 127.1, 129.4, 130.8, 131.0, 131.6, 132.30 (CH, C-aromatic), 129.80, 133.49, 134.19, 140.24, 141.71 (C, C-aromatic), 167.9 (C, C-15).

**Synthesis of 3-chloro-4-methyl benzoyl chloride (185)**

SOCl2 (0.696 g, 5.85 mmol) was added to a solution of 3-chloro-4-methyl benzoic acid (0.200 g, 1.17 mmol) in CH2Cl2 (2 mL). The mixture was stirred and refluxed for 24 hours, then evaporated until dryness to afford the desired product. **185** was obtained as yellow oil in 50% yield. **$^1$H-NMR ($CDCl_3$), δ**: 2.09 (s, 3H, H-7), 7.30 (d, J= 8.3 Hz, 1H, H3), 7.83 (dd, J1= 8.3 Hz, J2= 1.9 Hz, 1H, H-2), 8.01 (d, J= 1.9 Hz, 1H, H-aromatic). **$^{13}$C-NMR ($CDCl_3$), δ:** 20.6 (CH3, C-7), 129.4, 131.3, 131.7 (CH, C-aromatic), 99.0, 124.2, 134.2 (C, C-aromatic),
144.6 (C, C-8).

#### 1. Sulfonamide Compounds

##### 1.1 General procedure for the synthesis of: arylsulfonamido benzoic acids (OH14, 64-67)

2-Aminobenzoic acid (1 eq), the appropriate arylsulfonyl chloride (**1**, **59**-**62**) (1 eq) and NaOH (1 eq) were stirred in water (10 mL/mmol) at 70°C overnight. The product was filtered, washed with water, dried and purified by recrystallisation or flash column chromatography.

- **2-((2,4-Dichloro-5-methylphenyl)sulfonamido) benzoic acid (3 – OH14)**
  Obtained as white solid in 50% yield. **$^1$H-NMR ($CDCl_3$), δ:** 2.40 (s, 3H, H-7), 7.09-7.13 (m, 1H, H-aromatic), 7.41-7.43 (m, 1H, H-aromatic), 7.49-7.53 (m, 1H, H-aromatic), 7.84 (s, 1H, H-6), 7.95-7.97 (m, 1H, H-aromatic), 8,23 (s, 1H, H-3), 11.71 (bs, 1H, NH), 14.15 (bs, 1H, OH). **$^{13}$C-NMR ($CDCl_3$), δ:** 19.5 (CH3, C-7), 116.9, 123.5, 132.1, 132.2, 134.3, 135.2 (CH, C-aromatic), 116.1, 128.9, 134.4, 136.7, 139.5, 140.0 (C, C-aromatic), 170.3 (C, C-14). **UPLC-MS: Rt** 2.03 (98%) **MS (ESI)$^+$:** 360.9 $[M+H]^+$.
- **2-(((4-(Trifluoromethyl)phenyl)methyl)sulfonamido)benzoic acid (64)**
  Obtained as white solid in 23% yield. **$^1$H-NMR (DMSO-$D_6$), δ:** 4.86 (s, 2H, H-8), 7.16-7.19 (m, 1H, H-aromatic), 7.47 (d, J= 8.0 Hz, 2H, Haromatic), 7.55-7.60 (m, 2H, H-aromatic), 7.70 (d, J= 8.0 Hz, 2H, H-aromatic), 7.99-8.01 (m, 1H, Haromatic), 10.83 (bs, 1H, NH), 13.92 (bs, 1H, OH). **$^{13}$C-NMR (DMSO-$D_6$), δ:** 57.3 (CH2, C-8), 117.8, 123.1, 125.7, 132.0, 132.1, 135.1 (CH, C-aromatic), 129.3 (d, J= 32.2 Hz, C) 116.0, 132.4, 134.2, 141.0 (C-aromatic), 170.2 (C, C- 15). **$^{19}$F-NMR (DMSO-$D_6$), δ:** -61.04 (s, 3F). **UPLC-MS: Rt** 2.12 (>98%) **MS (ESI)$^-$:** 358.1 $[M-H]^-$ .
- **2-(Thiophene-2-sulfonamido)benzoic acid (65)**
  Obtained as grey solid in 22% yield. **$^1$H-NMR (DMSO-$D_6$), δ:** 7.13-7.15 (m, 1H, H-aromatic), 7.17-7.21 (m, 1H, H-aromatic), 7.60-7.64 (m,2H, H-aromatic), 7.67 (dd, J1= 3.7 Hz, J2= 1.3 Hz, 1H, H-aromatic), 7.91-7.94 (m, 1H, H-aromatic), 7.96(dd, J1= 4.9 Hz, J2= 1.3 Hz, 1H, H-aromatic), 11.23 (bs, 1H, NH), 14.09 (bs, 1H, OH). **$^{13}$C-NMR (DMSO-$D_6$), δ**: 119.3, 124.2, 128.3, 132.0, 134.0, 134.9, 135.0 (CH. C-aromatic), 117.6, 139.2, 139.9 (C, C-aromatic), 170.2 (C, C- 11). **UPLC-MS: Rt** 1.89 (99%) **MS (ESI)$^+$:** 283.9 $[M+H]^+$.
- **2-(Quinoline-8-sulfonamido)benzoic acid (66)**
  Obtained as white solid in 28% yield. **$^1$H-NMR (DMSO-$D_6$), δ:** 6.93-6.96 (m, 1H, H-aromatic), 7.38-7.41 (m, 1H, H-aromati), 7.62 (dd, J1=8.4 Hz, J2= 0.7 Hz, 1H, H-aromatic), 7.69 (dd, J1= 8.3 Hz, J2= 4.2 Hz, 1H, H-aromatic), 7.73-7.78 (m,2H, H-aromatic), 8.30 (dd, J1= 8.3 Hz, J2= 1.3 Hz, 1H, H-aromatic), 8.49 (dd, J1= 8.4 Hz, J2= 1.7 Hz, 1H,H-aromatic), 8.52 (dd, J1= 7.3 Hz, J2= 1.3 Hz, 1H, H-aromatic), 8.95 (dd, J1= 4.2 Hz, J2= 1.7 Hz, 1H, H-aromatic),11.55 (bs, 1H, NH), 13.74 (bs, 1H, OH). **$^{13}$C-NMR (DMSO-$D_6$), δ:** 117.1, 122.9, 123.2, 126.1, 131.8, 133.1, 134.6, 135.3, 137.5, 151.9 (CH, Caromatic),117.0, 128.8, 140.2, 142.8, 153.6 (C, C-aromatic), 169.41 (C, C- 16). **UPLC-MS: Rt** 1.87 (99%) **MS (ESI)$^+$:** 329.1 $[M+H]^+$.
- **2-(Isoquinoline-4-sulfonamido)benzoic acid (67)**
  Obtained as white solid in 35% yield. **$^1$H-NMR (DMSO-$D_6$), δ:** 7.13-7.16 (m, 1H, H-aromatic), 7.54-7.59 (m, 2H, H-aromatic), 7.75-7.78 (m,1H, H-aromatic), 7.85-7.87 (m,

1H, H-aromatic), 7.95-7.98 (m, 1H, H-aromatic), 8.09-8.11 (m, 1H, Haromatic),8.23 (dd, J1= 8.5 Hz, J2= 1.1 Hz, 1H, H-aromatic), 9.04-9.07 (m, 1H, H-aromatic), 9.14 (d,J= 2.3 Hz, 1H, H-aromatic), 11.29 (bs, 1H, NH), 13.72 (bs, 1H, OH).**$^{13}$C-NMR (DMSO-D$_6$), δ:** 119.8, 124.4, 128.9, 129.3, 130.2, 132.0, 133.5, 135.0, 137.4, 146.5 (CH, Caromatic),118.2, 126.3, 132.2, 139.4, 148.9 (C, C-aromatic), 169.9 (C, C- 16). **UPLC-MS: Rt** 1.93 (99%) **MS (ESI)$^+$:** 329.1 [M+H]$^+$.

##### 1.2 General procedure for the synthesis of: methyl(arylsulfonamidobenzoates) (21-26, 28, 30, 33-37, 42-43, 78, 88-93)

To a stirred solution of the differently substituted aniline (**18,19,20,77,84**) (1 eq) in anhydrous pyridine (4 L/mmol) was added the appropriate arylsulfonyl chloride (**1,4,5,6,8,10,13,14,15,16,17,40,41,62,63)** (1.2 eq) under an atmosphere of nitrogen. The reaction mixture was stirred at room temperature for 6h and was then quenched by addition of water (10 mL/mmol). The aqueous layer was extracted with ethyl acetate (20 mL/mmol) and the organic layer was then dried over anhydrous sodium sulfate. The solvent was removed under vacuum, and the crude product was purified by recrystallisation or flash column chromatography.

- **Methyl 2-((2,4-dichloro-5-methylphenyl)sulfonamido)benzoate (21)**
  Obtained as white solid in 49% yield. **$^1$H-NMR (CDCl$_3$)**, **δ**: 2.33 (s, 3H, H-7), 3.86 (s, 3H, H-15), 6.93-6.96 (m, 1H, H-aromatic), 7.30-7.34 (m,1H, H-aromatic), 7.35 (s, 1H, H-aromatic), 7.46 (dd, J1= 8.4 Hz, J2= 0.8 Hz, 1H, H-aromatic), 7.90 (dd, J1= 8.0 Hz, J2= 1.4 Hz, 1H, H-aromatic), 7.99 (s, 1H, H-aromatic), 11.12 (bs, 1H, NH). **$^{13}$C-NMR** (**CDCl$_3$**), **δ**: 19.6 (CH3, C-7), 52.5 (CH3, C-15), 116.9, 122.5, 131.4, 131.9, 134.5, 135.6 (CH,C-aromatic), 115.2, 115.9, 129.9, 133.5, 139.7, 140.0 (C, C-aromatic), 168.1 (C, C-14). **UPLC-MS: Rt** 2.59 (100%) **MS (ESI**)$^+$: 376.1 [M+H]$^+$
- **Methyl 3-((2,4-dichloro-5-methylphenyl)sulfonamido)benzoate (22)**
  Obtained as white solid in 56% yield. **$^1$H-NMR (CDCl$_3$)**, **δ**: 2.36 (s, 3H, H-7), 3.96 (s, 3H, H-15), 7.33-
  7.37 (m, 1H, H-aromatic), 7.39-7.42 (m, 1H, H-aromatic), 7.51 (s, 1H, H-aromatic), 7.76-7.77 (m, 1H,
  H-aromatic), 7.80-7.82 (m, 1H, H-aromatic),7.89 (s, 1H, H-aromatic), 11.20 (bs, 1H, NH).**$^{13}$C-NMR**
  (**CDCl$_3$**), **δ**: 19.5 (CH3, C-7), 52.3 (CH3, C-15), 122.4, 125.6, 126.9, 129.6, 131.6, 133.5 (CH, C-
  aromatic), 120.5, 128.4, 135.9, 135.9, 136.1, 156.6 (C, C-aromatic), 160.6 (C, C-14). **UPLC-MS: Rt**
  2.30 (100%) **MS (ESI**)$^-$: 372.1 [M-H]$^-$.
- **Methyl 4-((2,4-dichloro-5-methylphenyl)sulfonamido)benzoate (23)**
  Obtained as yellow powder in 46% yield. **$^1$H-NMR (DMSO-D$_6$)**, **δ**: 2.39 (s, 3H, H-7), 3.78 (s, 3H, H-

15), 7.19-7.22 (m, 2H, H-aromatic), 7.80 (s, 1H, H-aromatic), 7.81-7.84 (m, 2H, H-aromatic), 8.13 (s,
1H, H-aromatic), 11.23 (bs, 1H, NH). **$^{13}$C-NMR** (**DMSO-D$_6$**), **δ**: 19.0 (CH3, C-7), 51.8 (CH3, C-15), 117.1, 130.6, 131.5, 133.5 (CH, C-aromatic), 124.4, 128.7, 134.7, 136.0, 139.0, 141.4 (C, C-aromatic), 165.5 (C, C-14). **UPLC-MS: Rt** 2.29 (100%) **MS (ESI)**$^-$: 372.1 [M-H]$^-$.

- **Methyl 2-((4-(trifluoromethyl)phenyl)sulfonamido)benzoate (24)**
  Obtained as white solid in 60% yield. **$^1$H-NMR (CDCl$_3$), δ:** 3.89 (s, 3H, H-15), 7.10-7.13 (m, 1H, H-aromatic), 7.71-7.72 (m, 1H, H-aromatic), 7.73-7.75 (m, 2H, H-aromatic), 7.95-7.97 (m, 1H, H-aromatic), 7.98-8.00 (m, 2H, H-aromatic), 10.76 (bs, 1H, NH).**$^{13}$C-NMR (CDCl$_3$), δ:** 52.6 (CH3, C-15), 119.3, 123.6, 126.2, 127.7, 131.3, 134.5 (CH, C-aromatic), 116.2, 134.7, 139.8, 142.8 (C, C-aromatic), 168.3 (C, C-14). **$^{19}$F-NMR (CDCl$_3$), δ**: -63.19, (s, 3F). **UPLC-MS: Rt** 2.39 (100%) **MS (ESI)$^+$:** 360.1 [M+H]$^+$.
- **Methyl 2-((3,4-dimethylphenyl)sulfonamido)benzoate (25)**
  Obtained as white solid in 29% yield. **$^1$H-NMR (DMSO-D$_6$), δ:** 2.25 (s, 6H, 2xCH3, H-7,8), 3.83 (s, 3H, H-16), 7.14-7.18 (m, 1H, H-aromatic),7.31 (d, J=7.8 Hz, 1H, H-aromatic), 7.45-7.48 (m, 1H, H-aromatic), 7.52 (dd, J1= 8.0 Hz, J2= 1.7 Hz, 1H,H-aromatic), 7.55-7.58 (m, 1H, H-aromatic), 7.59-7.61 (m, 1H, H-aromatic), 7.85 (d, J= 7.8 Hz, 1H, H-aromatic), 10.43 (bs, 1H, NH).**$^{13}$C-NMR (DMSO-D$_6$), δ:** 19.7, 19.9 (2xCH3, C-7,8), 53.1 (CH3, C-16), 119.9, 124.1, 124.9, 127.8, 130.6, 131.4, 134.9 (CH, C-aromatic), 117.8, 136.4, 138.4, 139.3, 143.4 (C, C-aromatic), 168.1 (C, C-15). **UPLC-MS: Rt** 2.35 (100%) **MS (ESI)$^+$:** 320.1 [M+H]$^+$.
- **Methyl-2-((3,4-dichlorophenyl)sulfonamido)benzoate (26)**
  Obtained as white solid in 65% yield. **$^1$H-NMR (DMSO-D$_6$), δ:** 3.88 (s, 3H, H-14), 7.16-7.20 (m, 1H, H-aromatic), 7.38 (d, J= 8.5 Hz, 1H, H-aromatic), 7.52-7.56 (m, 1H, H-aromatic), 7.67-7.70 (m, 1H, H-aromatic), 7.91 (s, 1H, H-6), 7.92-7.94 (m, 1H, H-aromatic), 8.17 (d, J= 8.5 Hz, 1H, H-aromatic), 11.07 (bs, 1H, NH).**$^{13}$C-NMR (DMSO-D$_6$), δ:** 53.3 (CH3, C-14), 118.5, 124.2, 128.6, 131.7, 132.1, 133.7, 135.2 (CH, C-aromatic), 127.1, 132.3, 135.0, 138.5, 139.8 (C, C-aromatic), 168.2 (C, C-13). **UPLC-MS: Rt** 2.46 (100%) **MS (ESI)$^+$:** 362.1 [M+H]$^+$.
- **Methyl 2-((2,4,-dimethylphenyl)sulfonamido)benzoate (28)**
  Obtained as white solid in 55% yield. **$^1$H-NMR (DMSO-D$_6$), δ:** 2.30 (s, 3H, H-8), 2.51 (s, 3H, H-7), 3.87 (s, 3H, H-15), 7.09-7.14 (m, 1H, H-aromatic), 7.20-7.23 (m, 2H, H-aromatic), 7.31 (d, J= 8.3 Hz, 1H, H-aromatic), 7.50-7.55 (m, 1H, H-aromatic), 7.78 (d, J= 8.3 Hz, 1H, H-aromatic), 7.89-7.91 (m, 1H, H-aromatic), 10.73 (bs, 1H, NH).**$^{13}$C-NMR (DMSO-D$_6$),** δ: 19.7 (CH3, C-8), 21.2 (CH3, C-7), 53.2 (CH3, C-15), 118.3, 123.5, 127.4, 130.3,131.6, 133.9. 135.1 (CH, C-aromatic), 116.2, 134.2, 137.0, 139.5, 144.6 (C, C-aromatic), 168.4 (C, C-14). **UPLC-MS: Rt** 2.39 (100%) **MS (ESI)$^+$:** 320.2 [M+H]$^+$.
- **Methyl 2-(2,5-dichlorophenyl)sulfonamido)benzoate (30)**
  Obtained as white solid in 56% yield. **$^1$H-NMR (DMSO-D$_6$), δ:** 3.88 (s, 3H, H-14), 7.15-7.20 (m, 1H, H-aromatic), 7.42 (d, J= 8.1 Hz, 1H, H-aromatic), 7.55-7.59 (m, 1H, H-

aromatic), 7.71 (d, J= 8.5 Hz, 1H, H-3), 7.78 (dd, J1= 8.5 Hz, J2= 2.4 Hz, 1H, H-4), 7.91-7.94 (m, 1H, H-aromatic), 8.14 (d, J= 2.4 Hz, 1H, H-6), 11.08 (bs, 1H, NH). **$^{13}$C-NMR (DMSO-D$_6$),** δ: 53.3 (CH3, C-14), 119.0, 124.4, 131.4, 131.7, 134.3, 135.3, 135.5 (CH, C-aromatic), 117.5, 129.8, 132.9, 137.7, 138.2 (C, C-aromatic), 168.1 (C, C-13). **UPLC-MS: Rt** 2.43 (100%) **MS (ESI)$^-$:** 358.0 [M-H]$^-$.

- **Methyl 2-((phenylmethyl)sulfonamido)benzoate (33)**
  Obtained as white solid in 28% yield. **$^1$H-NMR (DMSO-D$_6$), δ:** 3.82 (s, 3H, H-15), 4.69 (s, 2H, H-7), 7.19-7.22 (m, 3H, H-aromatic), 7.29-7.33 (m, 2H, H-aromatic), 7.34-7.36 (m, 1H, H-aromatic), 7.56-7,58 (m, 1H, H-aromatic), 7.60-7.74 (m, 1H, H-aromatic), 7.95 (dd, J1= 7.9 Hz, J2= 1.3 Hz, 1H, H-aromatic), 10.11 (bs, 1H, NH).**$^{13}$C-NMR (DMSO-D$_6$),** δ: 40.5 (CH3, C-15), 57.9 (CH2, C-7), 118.8, 123.4, 128.8, 128.9, 131.2, 131.5, 135.2 (CH, C-aromatic), 116.2, 129.3, 140.5 (C, C-aromatic), 168.1 (C, C-14). **UPLC-MS: Rt** 2.23 (>98%) **MS (ESI)$^-$:** 304.1 [M-H]$^-$.
- **Methyl 2-(pyridine-3-sulfonamido)benzoate (34)**
  Obtained as white solid in 79% yield. **$^1$H-NMR (DMSO-D$_6$), δ:** 3.80 (s, 3H, H-13), 7.00-7.04 (m, 2H, H-aromatic), 7.29-7.33 (m, 1H, H-aromatic), 7.40-7.45 (m, 1H, H-aromatic), 7.66 (dd, J1= 8.4 Hz, J2= 0.9 Hz, 1H, H-aromatic), 7.86 (dd, J1= 8.0 Hz, J2= 1,5 Hz, 1H, H-aromatic), 8.03-8.07 (m, 1H, H-aromatic), 8.65-8.68 (m, 1H, H-aromatic), 8.98 (s, 1H, H-2), 10.71 (bs, 1H, NH).**$^{13}$C-NMR (DMSO-D$_6$),** δ: 52.6 (CH3, C-13), 119.4, 123.6, 123.7, 131.3, 134.7, 134.9, 148.1, 153.5 (CH, C-aromatic), 116.3, 136.0, 139.7 (C, C-aromatic), 168.3 (C, C-12). **UPLC-MS: Rt** 2.06 (>98%) **MS (ESI)$^+$:** 293.0 [M+H]$^+$.

- **Methyl 2-(naphthalene-2-sulfonamido)benzoate (35)**
  Obtained as pink solid in 68% yield. **$^1$H-NMR (CDCl$_3$), δ:** 3.86 (s, 3H, H-18), 7.01-7.04 (m, 1H, H-aromatic), 7.44-7.48 (m, 1H, H-aromatic), 7.59-7.65 (m, 2H, H-aromatic), 7.72 (dd, J1= 8.5 Hz, J2= 0.7 Hz, 1H, H-aromatic), 7.83 (dd, J1= 8.6 Hz, J2= 1.8 Hz, 1H, H-aromatic), 7.87 (d, J= 8.1 Hz, 1H, H-aromatic), 7.89-7.91 (m, 2H, H-aromatic), 7.95 (d, J= 8.1 Hz, 1H, H-aromatic), 8.48 (s, 1H, H-10), 10.80 (bs, 1H, NH).**$^{13}$C-NMR (CDCl$_3$),** δ: 52.4 (CH3, C-18), 119.0, 122.2, 122.9, 127.5, 128.8, 128.92, 128.95, 129.3, 129.4, 131.1, 134.5 (CH, C-aromatic), 115.9, 132.0, 134.9, 136.3, 140.42 (C, C-aromatic), 168.32 (C, C-17).**UPLC-MS: Rt** 2.37 (100%) **MS (ESI)$^+$:** 342.2 [M+H]$^+$.
- **Methyl 2-(naphthalene-1-sulfonamido)benzoate (36)**
  Obtained as white solid in 48% yield. **$^1$H-NMR (CDCl$_3$), δ:** 3.85 (s, 3H, H-18), 6.94-6.97 (m, 1H, H-aromatic), 7.37-7.40 (m, 1H, H-aromatic), 7.50-7.54 (m, 1H, H-aromatic), 7.57-7.61 (m, 1H, H-aromatic), 7.63 (dd, J1= 8.4 Hz, J2= 0.8 Hz, 1H, H-aromatic), 7.69-7.72 (m, 1H, H-aromatic), 7.84 (dd, J1= 8.0 Hz, J2= 1.4 Hz, 1H, H-aromatic), 7.91 (d, J= 8.4 Hz, 1H, H-aromatic), 8.04 (d, J= 8.0 Hz, 1H, H-aromatic), 8.37 (dd, J1= 7.4 Hz, J2= 1.4 Hz, 1H, H-aromatic), 8.74-8.76 (m, 1H, H-aromatic), 11.10 (bs, 1H, NH).**$^{13}$C-NMR (CDCl$_3$),** δ: 52.4 (CH3, C-18), 118.0, 122.4, 123.9, 124.5, 126.9, 128.4, 128.9, 130.3, 131.0, 134.3, 134.7 (CH, C-aromatic), 115.3, 128.0, 134.2, 140.4 (C, C-aromatic), 168.3 (C, C-17).**UPLC-MS: Rt** 2.37 (100%) **MS (ESI)$^+$:** 342.2 [M+H]$^+$.

- **Methyl 2-(cyclohexanesulfonamido)benzoate (37)**
  Obtained as colourless oil in 26% yield. **$^1$H-NMR (DMSO-D$_6$), δ:** 1.08-1.13 (m, 2H, H-cyclohexane), 1.15-1.18 (m, 1H, H-cyclohexane), 1.50-1.57 (m, 2H, H-cyclohexane), 1.58-1.60 (m, 1H, H-cyclohexane), 1.78-1.81 (m, 2H, H-cyclohexane), 2.06-2.08 (m, 2H, H-cyclohexane), 2.98 (tt, J1= 12.1 Hz, J2= 3.4 Hz, 1H, H-1), 3.86 (s, 3H, H-14), 6.98-7.01 (m, 1H, H-aromatic), 7.52-7.55 (m, 1H, H-aromatic), 7.73 (dd, J1= 8.4 Hz, J2= 0.5 Hz, 1H, H-aromatic), 7.95-7.97 (m, 1H, H-aromatic), 10.31 (bs, 1H, NH).**$^{13}$C-NMR (DMSO-D$_6$),** δ: 52.5 (CH3, C-14), 25.0, 25.1, 26.2 (CH2, C-cyclohexane), 61.1 (CH, C-1), 117.6, 122.1, 131.4, 134.8 (CH. C-aromatic), 114.6, 141.6 (C, C-aromatic), 168.4 (C, C-13). **UPLC-MS: Rt** 2.35 (99%) **MS (ESI)$^-$:** 296.2 [M-H]$^-$.
- **Methyl 2-((2-chloro-5-methylphenyl)sulfonamido)benzoate (42)**
  Obtained as white solid in 9% yield. **$^1$H-NMR (DMSO-D$_6$), δ:** 2.51 (s, 3H, CH3, H-7), 3.84 (s, 3H, H-15), 7.18-7.21 (m, 1H, H-aromatic), 7.38-741 (m, 1H, H-aromatic), 7.45 (d, J= 8.2 Hz , 1H, H-5), 7.56-7.60 (m, 1H, H-aromatic), 7.64 (dd, J1=8.2 Hz, J2= 2.2 Hz, 1H, H-4), 7.83-7.88 (m, 1H, H-aromatic), 7.89 (d, J= 2.2 Hz, 1H, H-2), 10.71 (bs, 1H,NH).**$^{13}$C-NMR (DMSO-D$_6$),** δ: 19.4 (CH3, C-7), 53.2 (CH3, C-15), 120.1, 124.5, 129.1, 131.3, 133.8, 135.0,135.2 (CH, C-aromatic), 110.1, 131.6, 134.7, 140.3, 147.9 (C, C-aromatic), 168.1 (C, C-14). **UPLC-MS: Rt** 2.49 (99%) **MS (ESI)$^-$:** 338.0 [M-H]$^-$.
- **Methyl 2-((3-chloro-4-methylphenyl)sulfonamido)benzoate (43)**
  Obtained as white solid in 24% yield. **$^1$H-NMR (DMSO-D$_6$), δ:** 2.36 (s, 3H, CH3, H-7), 3.82 (s, 3H, H-15), 7.19-7.23 (m, 1H, H-aromatic), 7.44 (dd, J1= 8.3 Hz, J2= 1.0 Hz, 1H, H-aromatic), 7.56-7.6 0 (m, 1H, H-aromatic), 7.61-7.62 (m, 2H, H-aromatics), 7.81-7.82 (m, 1H, H-aromatic), 7.84-7.86 (m, 1H, H-aromatic), 10.42 (bs, 1H, NH).**$^{13}$C-NMR (DMSO-D$_6$),** δ: 19.9 (CH3, C-7), 53.0 (CH3, C-15), 121.0, 124.7, 126.5, 129.7, 130.3, 131.5, 134.8 (CH, C-aromatic), 119.1, 137.8, 138.0, 138.5, 139.0 (C, C-aromatic), 167.8 (C, C-14). **UPLC-MS: Rt** 2.45 (100%) **MS (ESI)$^+$:** 340.1 [M+H]$^+$.
- **Isopropyl 2-((2,4-dichloro-5-methylphenyl)sulfonamido)benzoate (78)**
  Obtained as white solid in 26% yield. **$^1$H-NMR (CDCl$_3$), δ:** 1.40 (d, J= 6.3Hz, 6H, H-16,17), 2.43 (s, 3H, H-7), 5.27-5.29 (m, 1H, H-15), 7.01-7.05 (m, 1H, H-aromatic), 7.38-7.42 (m, 1H, H-aromatic), 7.44 (s, 1H, H-6), 7.53-7.56 (m, 1H, H-aromatic), 7.99 (dd, J1= 8.0 Hz, J2= 1.5 Hz, 1H, H-aromatic), 8.09 (s, 1H, H-3), 11.35 (bs, 1H, NH).**$^{13}$C-NMR (CDCl$_3$),** δ: 19.6 (CH3, C-7), 21.8 (2xCH3, C-16,17), 69.6 (CH, C-15), 116.9, 122.4, 131.4, 131.8, 133.5, 134.2 (CH, C-aromatic), 115.9, 129.9, 134.9, 135. 5, 139.6, 139.9 (C, C-aromatic), 167.2 (C, C-14). **UPLC-MS: Rt** 2.78 (100%) **MS (ESI)$^+$:** 400.1 [M+H]$^+$.
- ***N*-(2-(1*H*-tetrazol-5-*yl*)phenyl)-2,4-dichloro-5-methylbenzenesulfonamide (88)**
  Obtained as off-white solid in 18% yield. **$^1$H-NMR (DMSO-D$_6$), δ:** 2.39 (s, 3H, H-7), 7.28-7.32 (m, 1H, H-aromatic), 7.48-7.51 (m, 2H, H-aromatic), 7.78 (s, 1H, H-2), 7.94-7.98 (m, 1H, H-aromatic), 8.16 (s, 1H, H-5), 11.37 (bs, 1H, NH). **$^{13}$C-NMR (DMSO-D$_6$),** δ: 19.5 (CH3, C-7), 119.2, 124.8, 129.5, 132.1, 132.7, 134.0 (CH, C-aromatic), 113.5, 129.1, 134.7, 135.5, 136.6, 136.8 (C, C-aromatic), 172.3 (C, C-14). **UPLC-MS: Rt** 2.26 (>97%) **MS (ESI)$^+$:** 386.1 [M+H]$^+$.

- ***N*-(2-(1*H*-tetrazol-5-yl)phen*yl*)-3-chloro-4-methylbenzenesulfonamide (89)**
  Obtained as white solid in 16% yield. **$^{1}$H-NMR (DMSO-D$_6$), δ:** 2.33 (s, 3H, H-7), 7.33-7.36 (m, 1H, H-aromatic), 7.45-7.48 (m, 2H, H-aromatic), 7.51-7.55 (m, 2H, H-aromatic), 7.66-7.68 (m, 1H, H-aromatic), 7.86-7.88 (m, 1H, H-aromatic), 10.67 (bs, 1H, NH).**$^{13}$C-NMR (DMSO-D$_6$),** δ: 20.1 (CH3, C-7), 122.3, 125.8, 125.9 127.3, 129.7, 132.3, 132.4 (CH, C-aromatic), 116.6, 134.6, 135.6, 138.2, 142.1 (C, C-aromatic), 179.4 (C, C-15). **UPLC-MS: Rt** 2.15 (>99%) **MS (ESI)$^+$:** 350.3 [M+H]$^+$.
- ***N*-(2-(1*H*-tetrazol-5-yl)phen*yl*)-4-(trifluoromethyl)benzenesulfonamide (90)**
  Obtained as white solid in 42% yield. **$^{1}$H-NMR (DMSO-D$_6$), δ:** 7.36-6.39 (m, 1H, H-aromatic), 7.44 (dd; J1= 8.1 Hz, J2= 1.0 Hz, 1H, H-aromatic), 7.52-7.55 (m, 1H, H-aromatic), 7.85 (dd, J1= 7.8 Hz, J2= 1.4 Hz, 1H, H-aromatic), 7.88-7.91 (m, 4H, H-aromatic), 10-73 (bs, 1H, NH).**$^{13}$C-NMR (DMSO-D$_6$),** δ: 123.0, 126.2, 126.9, 128.3, 129.9, 132.5 (CH, C-aromatic), 133.2 (d, J= 37.7 Hz, C) 117.0, 122.6, 124.8, 135.3, 143.0 (C-aromatic). **$^{19}$F-NMR (DMSO-D6), δ:** -61.76 (s, 3F). **UPLC-MS: Rt** 1.81 (100%) **MS (ESI)$^+$:** 370.1 [M+H]$^+$.
- ***N*-(2-(1*H*-tetrazol-5-*yl*)phenyl)naphthalene-2-sulfonamide (91)**
  Obtained as white solid in 38% yield. **$^{1}$H-NMR (DMSO-D$_6$), δ:** 7.26-7.29 (m, 1H, H-aromatic), 7.47-7.50 (m, 1H, H-aromatic), 7.54 (d, J=8.1Hz, 1H, H-aromatic), 7.63-7.66 (m, 1H, H-aromatic), 7.68-7.71 (m, 2H, H-aromatic), 7.82-7.84 (m, 1H, H-aromatic), 7.97-8.02 (m, 2H, H-aromatic), 8.10 (d, J= 8.1 Hz, 1H, H-aromatic), 8.47 (s, 1H, H-10), 10.78 (bs, 1H, NH).**$^{13}$C-NMR (DMSO-D$_6$),** δ: 121.5, 122.2, 125.4, 128.26, 128.29, 128.9, 129.6, 129.73, 129.78, 129.9, 134.8 (CH, C-aromatic), 136.0, 136.1(C, C-aromatic). **UPLC-MS: Rt** 2.07 (100%) **MS (ESI)$^+$:** 352.1 [M+H]$^+$.
- ***N*-(2-(1*H*-tetrazol-5-*yl*)phenyl)quinoline-8-sulfonamide (92)**
  Obtained as off-white solid in 26% yield. **$^{1}$H-NMR (DMSO-D$_6$), δ:** 7.11-7.14 (m, 1H, H-aromatic), 7.39-7.42 (m, 1H, H-aromatic), 7.55-7.58 (m, 1H, H-aromatic), 7.72-7.76 (m, 3H, H-aromatic), 8.26-8.28 (m, 1H, H-aromatic), 8.41-8.44 (m, 2H, H-aromatic), 8.48-8.49 (m, 1H, H-aromatic), 11.23 (bs, 1H, NH).**$^{13}$C-NMR (DMSO-D$_6$),** δ: 112.9, 118.6, 123.1, 124.1, 128.3, 129.3, 132.4, 134.4, 136.5, 151.3 (CH, C-aromatic), 126.1, 134.4, 137.4, 142.6 (C, C-aromatic). **UPLC-MS: Rt** 1.65 (>96%) **MS (ESI)$^+$:** 353.3 [M+H]$^+$.
- ***N*-(2-(1*H*-tetrazol-5-*yl*)phenyl)isoquinoline-4-sulfonamide (93)**
  Obtained as brown solid in 46% yield. **$^{1}$H-NMR (DMSO-D$_6$), δ:** 7.05-7.08 (m, 1H, H-aromatic), 7.21-7.25 (m, 1H, H-aromatic), 7.64 (d, J= 8.2 Hz, 1H, H-aromatic), 7.69-7.72 (m, 1H, H-aromatic), 7.88-7.91 (m, 1H, H-aromatic), 8.01 (d, J= 8.5 Hz, 1H, H-aromatic), 8.03-8.05 (m, 1H, H-aromatic), 8.09-8.11 (m, 1H, H-aromatic), 8.89 (s, 1H, H-3), 8.97 (s, 1H, H-2), 13.23 (bs, 1H, NH).**$^{13}$C-NMR (DMSO-D$_6$),** δ: 118.8, 124.3, 127.7, 128.6, 128.8, 129.2, 130.0, 133.2, 137.0, 146.4 (CH, C-aromatic), 120.5, 126.2, 134.9, 146.4, 148.7 (C, C-aromatic), 159.6 (C, C- 16). **UPLC-MS: Rt** 1.85 (100%) **MS (ESI)$^+$:** 353.1 [M+H]$^+$.

**1.3 General procedure for the synthesis of: alkyl(arylsulfonamido)benzoates (50, 52, 56-57)**

To a cooled (0°) stirring solution of alkyl anthranilate (**44,18**) (1 eq) in CH2Cl2 (4 mL/mmol) was added pyridine (7.5 eq). The appropriate arylsulfonyl chlorides (**1,46,48,49)** (1.2 eq) was then added slowly. The solution was allowed to warm to room temperature and stirred for 1-3 hours. The reaction was poured into saturated NaHCO3 solution (14 mL/mmol), extracted with CH2Cl2 (3 x 5mL/mmol), and washed with 1M HCl (15 mL/mmol). The combined organic phases were dried over MgSO4 and concentrated *in vacuo*. The desired products were purified by recrystallization or flash column chromatography.

- **Ethyl 2-((2,4-dichloro-5-methylphenyl)sulfonamido)benzoate (50)**
  Obtained as pink solid in 43% yield. **$^{1}$H-NMR (DMSO-D$_6$), δ:** 1.33 (t, J= 7.1 Hz, 3H, H-9), 2.39 (s, 3H, H-16), 4.36 (q, J= 7.1 Hz, 2H, H-8), 7.14-7.18 (m, 1H, H-aromatic), 7.40-7.42 (m, 1H, H-aromatic), 7.52-7.56 (m, 1H, H-aromatic), 7.83 (s, 1H, H-aromatic), 7.95 (dd, J1= 7.0 Hz, J2= 1.0 Hz, 1H, H-aromatic), 8.19 (s, 1H, H-aromatic), 11.10 (bs, 1H, NH).**$^{13}$C-NMR (DMSO-D$_6$),** δ: 14.3 (CH3), 19.5 (CH3), 62.3 (CH2, C-8), 118.0, 124.0, 131.7, 132.1, 134.1, 134.5 (CH, C-aromatic), 116.7, 129.0, 135.3, 136.7, 138.7, 140.0 (C, C-aromatic), 167.8 (C, C-7). **UPLC-MS: Rt** 2.69 (100%) **MS (ESI)$^-$:** 386.2 [M-H]$^-$.
- **Methyl 2-((4-iodophenyl)sulfonamido)benzoate (52)**
  Obtained as beige solid in 25% yield. **$^{1}$H-NMR (DMSO-D$_6$), δ:** 3.81 (s, 3H, H-8), 7.20-7.24 (m, 1H, H-aromatic), 7.42-7.44 (m, 1H, H-aromatic), 7.53 (d, J= 8.6 Hz, 2H, H-aromatic), 7.68-7.60 (m, 1H, H-aromatic), 7.85 (dd, J1= 6.5 Hz, J2= 1.5 Hz, 1H, H-aromatic), 7.95 (d, J= 8.6 Hz, 2H, H-aromatic), 10.41 (bs, 1H, NH).**$^{13}$C-NMR (DMSO-D$_6$),** δ: 53.1 (CH3, C-8), 121.2, 124.9, 128.9, 131.5, 134.8, 138.8 (CH, C-aromatic), 102.4, 119.4, 138.3, 138.9 (C, C-aromatic), 167.8 (C, C-7). **UPLC-MS: Rt** 2.39 (100%) **MS (ESI)$^+$:** 418.1 [M+H]$^+$.
- **Methyl 2-((p-tolylmethyl)sulfonamido)benzoate (56)**
  Obtained as white solid in 15% yield. **$^{1}$H-NMR (CDCl$_3$), δ:** 2.34 (s, 3H, H-7), 3.85 (s, 3H, H-16), 4.37 (s, 2H, H-8), 7.06-7.14 (m, 5H, H-aromatic), 7.50-7.53 (m, 1H, H-aromatic), 7.73-7.75 (m, 1H, H-aromatic), 8.02 (dd, J1= 7.9 Hz, J2= 1.6Hz, 1H, H-aromatic), 10.28 (bs, 1H, NH).**$^{13}$C-NMR (CDCl$_3$),** δ: 21.1 (CH3, C-7), 52.4 (CH3, C-16), 58.0 (CH2, C-8), 118.0, 122.6, 129.4, 130.5, 131.3, 134.7 (CH, C-aromatic), 115.4, 125.1, 138.7, 141.0 (C-aromatic), 167.9 (C, C- 15). **UPLC-MS: Rt** 2.30 (100%) **MS (ESI)$^-$:** 318.0 [M-H]$^-$.
- **Methyl 2-(furan-2-sulfonamido)benzoate (57)**
  Obtained as pink solid in 57% yield. **$^{1}$H-NMR (DMSO-D$_6$), δ:** 3.84 (s, 3H, H-12), 6.65 (dd, J1= 3.5 Hz, J2= 1.7 Hz, 1H, H-aromatic), 7.24-7.27(m, 2H, H-aromatic), 7.49 (dd, J1= 8.2 Hz, J2= 0.6 Hz, 1H, H-aromatic), 7.60-7.63 (m, 1H, H-aromatic), 7.89 (dd, J1= 7.8, J2= 1.3 Hz, 1H, H-aromatic), 7.96-7.98 (m, 1H, H-aromatic), 10.62 (bs, 1H, NH).**$^{13}$C-NMR (DMSO-D$_6$),** δ: 53.1 (CH3, C-12), 112.2, 118.6, 121.3, 125.0, 131.4, 134.7, 148.6 (CH, C-aromatic), 119.6, 137.9, 147.1 (C-aromatic), 167.9 (C, C- 11). **UPLC-MS: Rt** 2.10 (100%) **MS (ESI)$^+$:** 282.0 [M+H]$^+$.

**1.4 General procedure for the synthesis of: arylsulfonamido benzoic acids (72-76)**

To a solution of the appropriate methyl-(arylsulfonamido) benzoate (**34-37,57**) (1 eq) in THF (7 mL/mmol), H2O (3.5 mL/mmol), MeOH (3.5 mL/mmol) was added LiOH (3 eq). The reaction was stirred at 70°C overnight. After the reaction was allowed to cool to room temperature, the reaction mixture was acidified with 2M HCl. The product was extracted with ethyl acetate (3x10 mL/mmol). The organic layer was dried over MgSO4 and evaporated under reduced pressure. The product was purified by flash column chromatography or recrystallisation.

- **2-(Pyridine-3-sulfonamido)benzoic acid (72)**
  Obtained as off-white solid in 30% yield. **$^1$H-NMR (DMSO-D$_6$), δ:** 7.17-7.20 (m, 1H, H-aromatic), 7.52-7.54 (m, 1H, H-aromatic), 7.57-7.59 (m, 1H, H-aromatic), 7.60-7.62 (m, 1H, H-aromatic), 7.90 (dd, J1= 7.9 Hz, J2= 1.5 Hz, 1H, H-aromatic), 8.19-8.22 (m, 1H, H-aromatic), 8.80-8.82 (m, 1H, H-aromatic), 8.95-8.97 (m, 1H, H-aromatic), 11.23 (bs, 1H, NH), 13.60 (bs, 1H, OH).**$^{13}$C-NMR (DMSO-D$_6$),** δ: 119.8, 124.4, 124.9, 132.0, 134.9, 135.4, 147.7, 154.4 (CH, C-aromatic), 118.3, 125.6, 135.6 (C, C-aromatic), 169.93(C, C- 12).**UPLC-MS: Rt** 1.66 (99%) **MS (ESI)$^+$:** 278.9 [M+H]$^+$.
- **2-(Naphthalene-2-sulfonamido)benzoic acid (73)**
  Obtained as white solid in 68% yield. **$^1$H-NMR (DMSO-D$_6$), δ:** 7.07-7.11 (m, 1H, H-aromatic), 7.51-7.55 (m, 1H, H-aromatic), 7.57-7.59 (m, 1H, H-aromatic), 7.66-7.69 (m, 1H, H-aromatic), 7.70-7.74 (m, 1H, H-aromatic), 7.76-7.79 (m, 1H, H-aromatic) 7.87 (d, J= 8.0 Hz, 1H, H-aromatic), 8.01 (d, J= 8.0 Hz, 1H, H-aromatic), 8.09 (d, J= 8.7 Hz, 1H, H-aromatic), 8.17 (d, J= 8.2 Hz, 1H, H-aromatic), 8.60 (s, 1H, H-10), 11.28 (bs, 1H, NH), 14.00 (bs, 1H, OH).**$^{13}$C-NMR (DMSO-D$_6$),** δ: 118.8, 122.2, 123.8, 128.30, 128.34, 129.0, 129.81, 129.85, 130.2, 131.9, 134.9 (CH, C-aromatic), 117.1, 132.0, 134.9, 136.0, 140.2 (C, C-aromatic), 170.2 (C, C-17). **UPLC-MS: Rt** 2.12 (100%) **MS (ESI)$^+$:** 328.1 [M+H]$^+$.
- **2-(Naphthalene-1-sulfonamido)benzoic acid (74)**
  Obtained as yellow solid in 90% yield. **$^1$H-NMR (DMSO-D$_6$), δ:** 7.04-7.09 (m, 2H, H-aromatic), 7.48-7.53 (m, 2H, H-aromatic), 7.69-7.74 (m, 1H, H-aromatic), 7.76-7.80 (m, 1H, H-aromatic), 7.86 (d, J= 7.8 Hz, 1H, H-aromatic), 8.14 (d, J= 8.1 Hz, 1H, H-aromatic), 8.31 (d, J= 7.8 Hz, 1H, H-aromatic), 8.40 (d, J= 7.2 Hz, 1H, H-aromatic), 8.59 (d, J= 8.6 Hz, 1H, H-aromatic), 11.82 (bs, 1H, NH), 14.20 (bs, 1H, OH). **$^{13}$C-NMR (DMSO-D$_6$),** δ: 117.5, 123.2, 123.8, 124.9, 127.4, 127.6, 128.9, 131.0, 131.9, 134.9, 135.59 (CH, C-aromatic), 116.1, 129.8, 134.2, 140.5, 142.0 (C, C-aromatic), 170.3 (C, C-17). **UPLC-MS: Rt** 2.09 (100%) **MS (ESI)$^+$:** 328.1 [M+H]$^+$.
- **2-(Cyclohexanesulfonamido)benzoic acid (75)**
  Obtained as white solid in 26% yield. **$^1$H-NMR (DMSO-D$_6$), δ:** 1.08-1.14 (m, 1H, H-cyclohexane), 1.17-1.26 (m, 2H, H-cyclohexane), 1.38- 1.49 (m, 2H, H-cyclohexane), 1.53-1.60 (m, 1H, H-cyclohexane), 1.74-1.77 (m, 2H, H-cyclohexane), 1.97-2.00 (m, 2H, H-cyclohexane), 3.19-3.25 (m, 1H, H-1), 7.13-7.17 (m, 1H, H-aromatic), 7.58-7.62 (m, 1H, H-aromatic), 7.64 (dd, J1= 8.3 Hz, J2= 0.9 Hz, 1H, H-aromatic), 8.01 (dd, J1= 7.9 Hz, J2= 1.4 Hz, 1H, H-aromatic), 11.01 (bs, 1H, NH). **$^{13}$C-NMR (DMSO-D$_6$),** δ 26.4,

25.0, 26.28(CH2, C-cyclohexane), 60.0 (CH, C-1), 117.8, 122.8, 132.1, 135.0 (CH. C-aromatic), 132.2, 141.5 (C, C-aromatic), 170.5 (C, C- 13). **UPLC-MS: Rt** 2.04 (>99%) **MS (ESI)$^-$:** 282.1 [M-H]$^-$.

- **2-(Furan-2-sulfonamido)benzoic acid (76)**
  Obtained as white solid in 32% yield. **$^1$H-NMR (DMSO-D$_6$), δ:** 6.66-6.68 (m, 1H, H-aromatic), 7.17-7.20 (m, 1H, H-aromatic), 7.35 (dd, J1= 7.3 Hz, J2= 0.6 Hz, 1H, H-aromatic), 7.52-7.54 (m, 1H, H-aromatic), 7.58-7.62 (m, 1H, H-aromatic) 7.94-7.97 (m, 2H, H-aromatic), 11.44 (bs, 1H, NH), 13.89 (bs, 1H, OH). **$^{13}$C-NMR (DMSO-D$_6$),** δ 112.2, 118.6, 119.0, 124.8, 131.9, 135.0, 148.7 (CH, C-aromatic), 117.4, 139.5, 146.8 (C, C-aromatic), 170.2 (C, C- 11). **UPLC-MS: Rt** 1.81 (100%) **MS (ESI)$^-$:** 266.0 [M-H]$^-$.

**1.5 Synthesis of Isopropyl 2-((3-chloro-4-methylphenyl)sulfonamido)benzoate (79)**

2-((3-chloro-4-methylphenyl)sulfonamido)benzoic acid (0.100 g, 0.3 mmol), TBTU (0.09 g, 0.3 mmol) and DIPEA (0.08 g, 0.6 mmol) were stirred in anhydrous DMF (1 mL), and the resulting mixture was stirred at room temperature for 30 min, under N2 atmosphere. Isopropyl alcohol (0.02 g, 0.3 mmol) was then injected into the reaction mixture via syringe and stirring was continued at room temperature until completion of the reaction. The reaction mixture was diluted with DCM (10 mL) and the resulting mixture was extracted with sat.aq NH4Cl (10 mL) and sat.aq NaHCO3 (10 mL). The organic layer was dried over MgSO4 and the solvent was evaporated under vacuum. The desired product was purified by flash column chromatography (Biotage Isolera One automated flash column chromatography cartridge: ZIP KP SIL 10g, *n*-hexane-EtOAc 100:0 increasing to *n*-hexane-EtOAc 70:30 in 10 CV). **79** was obtained as colourless oil in 22% yield. **$^1$H-NMR (CDCl$_3$), δ:** 1.26 (d, J= 6.2 Hz, 6H, H-16,17), 2.30 (s, 3H, H-7), 5.11-5.14 (m, 1H, H-15), 6.97- 7.00 (m, 1H, H-aromatic), 7.17-7.19 (m, 1H, H-aromatic), 7.37-7.41 (m, 1H, H-aromatic), 7.52 (dd, J1= 7.8 Hz, J2= 1.9 Hz, 1H, H-aromatic), 7.59 (dd, J1= 8.3 Hz, J2= 1.0 Hz, 1H, H-aromatic), 7.74-7.76 (m, 1H, H-aromatic), 7.85 (m, 1H, H-aromatic), 10.68 (bs, 1H, NH). **$^{13}$C-NMR (CDCl$_3$),** δ: 20.2 (CH3, C-7), 21.7 (2xCH3, C-16,17), 69.5 (CH, C-15), 119.4, 123.2, 125.3, 127.7, 131.1, 131.3, 134.3 (CH, C-aromatic), 116.9, 135.1, 138.3, 140.1, 141.8 (C, C-aromatic), 167.3 (C, C-14). **UPLC-MS: Rt** 2.77 (100%) **MS (ESI)$^+$:** 368.8 [M+H]$^+$.

**1.6 Synthesis of *Tert*-Butyl 2-((2,4-dichloro-5-methylphenyl)sulfonamido)benzoate (80)**

Concentrated sulfuric acid (1eq) was added to a vigorously stirred suspension of anhydrous magnesium sulfate (4eq) in DCM (4mL/mmol). The mixture was stirred for 15 minutes, after which the 2-((2,4-dichloro-5-methylphenyl)sulfonamido)benzoic acid (**OH14**) (0.150 g, 0.416 mmol) was added. *Tert*-Butyl alcohol (5eq) was added last. The mixture was stoppered tightly and stirred at 25°C for 18h. The reaction mixture was then quenched with sat.aq. NaHCO3 solution and stirred until all the MgSO4 was dissolved. The solvent phase was separated, washed with brine, dried over MgSO4 and concentrated to afford the crude *tert*-butyl ester, which was purified by flash column chromatography (Biotage Isolera One automated flash column chromatography, cartridge: SNAP KP SIL 25g, *n*-hexane-EtOAc 100:0 increasing to *n*-hexane-EtOAc 60:40 in 15CV). **80** was obtained as yellow solid in 20% yield. **$^1$H-NMR (CDCl$_3$),**

**δ:** 1.32 (s, 9H, H-16,17,18), 2.32 (s, 3H, H-7), 6.90-6.93 (m, 1H, H-aromatic), 7.26- 7.30 (m, 1H, H-aromatic), 7.34 (s, 1H, H-6), 7.44 (dd, J1= 7.3 Hz, J2= 1.1 Hz, 1H, H-aromatic), 7.82 (dd, J1= 6.3 Hz, J2= 1.6 Hz, 1H, H-aromatic), 7.98 (s, 1H, H-3), 11.32 (bs, 1H, NH). **$^{13}$C-NMR ($CDCl_3$),** δ: 19.6 (CH3, C-7), 28.1 (3xCH3, C-16,17,18), 117.0, 122.4, 131.5, 131.8, 133.6, 133.9 (CH, C-aromatic), 83.0 (C, C-15), 129.8, 134.9, 134.5, 135.5, 139.5, 139.9 (C, C-aromatic), 167.0 (C, C-14). **UPLC-MS: Rt** 2.38 (>97%%) **MS (ESI)$^-$:** 414.1 $[M-H]^-$.

**1.7 General procedure for the synthesis of: alkyl(arylsulfonamido) carboxamide (101-105)**

A suspension of CDI (1.2eq) and the appropriate carboxylic acid (**3,69**) (1eq) in THF (6 mL/mmol) was stirred at room temperature overnight. A 2N solution of alkylamine in THF (1.5eq) was then added and the resulting solution was stirred at room temperature for 6 hours. The solvent was evaporated and the residue dissolved in ethyl acetate (10mL/mmol). The solution was washed sequentially with water and brine. The organic layer was dried over MgSO4 and then evaporated. Desired products were purified by column chromatography or recrystallisation.

- **2-((3-Chloro-4-methylphenyl)sulfonamido)-*N*-methylbenzamide (101)**
  Obtained as white solid in 30% yield. **$^1$H-NMR ($CDCl_3$), δ:** 2.39, (s, 3H, H-7), 2.94 (d, J= 4.8 Hz, 3H, H-15), 6.06 (bs, 1H, NH), 7.07-7.11 (m, 1H, H-aromatic), 7.26-7.29 (m, 1H, H-aromatic), 7.36 (d, J= 7.9 Hz, 1H, H-3), 7.42-7.46 (m, 1H, H-aromatic), 7.60 (dd, J1= 8.1 Hz, J2= 1.7 Hz, 1H, H-aromatic), 7.69 (d, J= 7.9 Hz, 1H, H-4), 7.79-7.81 (m, 1H, H-aromatic), 10.87 (bs, 1H, NH). **$^{13}$C-NMR ($CDCl_3$),** δ 20.2 (CH3), 26.7 (CH3), 121.7, 123.8, 125.4, 126.5, 127.7, 131.1, 132.6 (CH, C-aromatic), 121.8, 135.0, 138.4, 138.5, 141.5 (C, C-aromatic), 168.8 (C, C-14). **UPLC-MS: Rt** 2.11 (>99%) **MS (ESI)$^+$:** 339.1 $[M+H]^+$.
- **2-((2,4-Dichloro-5-methylphenyl)sulfonamido)-*N*-methylbenzamide (102)**
  Obtained as white solid in 49% yield. **$^1$H-NMR ($CDCl_3$), δ:** 2.41 (s, 3H, H-7), 3.02 (d, J=4.8 Hz, 3H, H-15), 6.17 (bs, 1H, NH), 7.02-7.05 (m, 1H, H-aromatic), 7.34-7.37 (m, 1H, H-aromatic), 7.40 (dd, J1= 7.8 Hz, J2= 1.0 Hz, 1H, H-aromatic), 7.45 (s, 1H, H-4), 7.54-7.57 (m, 1H, H-aromatic), 8.05 (s, 1H, H-1), 11.37 (bs, 1H, NH). **$^{13}$C-NMR ($CDCl_3$),** δ 19.6 (CH3), 26.8 (CH3), 118.7, 122.9, 126.7, 131.7, 132.5, 133.3 (CH, C-aromatic), 120.6, 130.0, 135.2, 135.5, 138.0, 139.7 (C, C-aromatic), 168.9 (C, C-14). **UPLC-MS: Rt** 2.23 (100%) **MS (ESI)$^+$:** 375.1 $[M+H]^+$.
- **2-((2,4-Dichloro-5-methylphenyl)sulfonamido)-*N*-ethylbenzamide (103)**
  Obtained as white solid in 44% yield. **$^1$H-NMR ($CDCl_3$), δ:** 1.28 (t, J= 7.2 Hz, 3H, H-16), 2.41 (s, 3H, H-7), 3.47-3.52 (m, 2H, H-15), 6.13 (bs, 1H, NH), 7.02-7.06 (m, 1H, H-aromatic), 7.34-7.37 (m, 1H, H-aromatic), 7.40-7.42 (m, 1H, H-aromatic), 7.45 (s, 1H, H-4), 7.55-7.57 (m, 1H, H-aromatic), 8.05 (s, 1H, H-1), 11.40 (bs, 1H, NH). **$^{13}$C-NMR ($CDCl_3$),** δ 14.6 (CH3), 19.6 (CH3), 35.0 (CH2, C-15), 118.7, 122.9, 126.7, 131.7, 132.5, 133.3 (CH, C-aromatic), 120.8, 130.0, 135.2, 135.5, 138.1, 139.7 (C, C-aromatic), 168.1 (C, C-14). **UPLC-MS: Rt** 2.33 (>98%) **MS (ESI)$^+$:** 389.1 $[M+H]^+$.
- **2-((2,4-Dichloro-5-methylphenyl)sulfonamido)-*N*,*N*-dimethylbenzamide (104)**

Obtained as white solid in 35% yield. **$^1$H-NMR ($CDCl_3$), δ:** 2.37 (s, 3H, H-7), 2.79 (s, 3H), 3.10 (s, 3H), 7.09-7.12 (m, 1H, H-aromatic), 7.18- 7.20 (m, 1H, H-aromatic), 7.32-7.36 (m, 1H, H-aromatic), 7.49 (s, 1H, H-4), 7.59-7.60 (m, 1H, H-aromatic), 7.93 (s, 1H, H-1), 8.88 (bs, 1H, NH). **$^{13}$C-NMR ($CDCl_3$),** δ 19.6 (CH3), 39.4 (CH3), 39.6 (CH3), 122.4, 124.0, 127.9, 130.8, 131.4, 133.2 (CH, C-aromatic), 126.1, 129.7, 135.1, 135.2, 135.7, 139.7 (C, C-aromatic), 169.3 (C, C-14). **UPLC-MS: Rt** 2.18 (>97%) **MS (ESI)$^+$:** 389.1 $[M+H]^+$.

- **2-((2,4-Dichloro-5-methylphenyl)sulfonamido)-*N*-isopropylbenzamide (105)**
  Obtained as white solid in 53% yield. **$^1$H-NMR ($CDCl_3$), δ:** 1.19, d, J=6.5 Hz, 6H, H-16,17), 2.31 (s, 3H-H-7), 4.14-4.20 (m, 1H-H-15), 5.85 (bs, 1H, NH), 6.92-6.95 (m, 1H, H-aromatic), 7.24-7.27 (m, 1H, H-aromatic), 7.30 (dd, J1= 7.8 Hz, J2= 1.4 Hz, 1H, H-aromatic), 7.35 (s, 1H, H-aromatic), 7.45 (dd, J1= 1.0 Hz, J2= 8.4 Hz, 1H, H-aromatic), 7.96 (s, 1H, H-4), 11.32 (bs, 1H, NH).**$^{13}$C-NMR ($CDCl_3$),** δ 19.6 (CH3, C-7), 22.6 (2xCH3, C-16,17), 42.1 (CH, C-15), 118.7, 122.9, 126.8, 131.7, 132.5, 133.4 (CH, C-aromatic), 120.8, 130.0, 135.2, 135.5, 138.1, 139.7 (C, C-aromatic), 167.5 (C, C-14). **UPLC-MS: Rt** 2.43 (>97%) **MS (ESI)$^+$:** 403.2 $[M+H]^+$.

##### 1.8 Synthesis of: 2-((2,4-Dichloro-5-methylphenyl)sulfonamido) benzamide (106)

N,N'-carbonyldiimidazole (0.050 g, 0.33 mmol) was added to a solution of 2-((2,4-dichloro-5-methylphenyl)sulfonamido)benzoic acid (0.100 g, 0.27 mmol) in [bmim]BF4 (1 mL). The resulting mixture was stirred at 80°C for 2h. Ammonium acetate (0.083 g, 1.08 mmol) and triethylamine (0.082 g, 0.81 mmol) were added to the reaction mixture. The reaction was heated at 80°C for 3h. After completion of the reaction, the product was extracted with ethyl acetate (10 mL), washed with 0.01N solution of HCl (15 mL), and dried over anhydrous MgSO4. The solvent was evaporated in vacuo, and the crude product was purified by flash column chromatography (Biotage Isolera One automated flash column chromatography, cartridge: SNAP KP SIL 25g, n-hexane:EtOAc 100:0 increasing to *n*-hexane-EtOAc 0:100 in 14CV). **106** was obtained as grey solid in 10% yield. **$^1$H-NMR ($CDCl_3$), δ:** 2.32 (s, 3H, H-7), 5.87 (bs, 2H, NH2), 6.94-6.98 (m, 1H, H-aromatic), 7.29-7.32 (m, 1H, H-aromatic), 7.36 (s, 1H, H-4), 7.39-7.41 (m, 1H, H-aromatic), 7.48-7.51 (m, 1H, H-aromatic), 7.97 (s, 1H, H-1), 11.50 (bs, 1H, NH). **$^{13}$C-NMR ($CDCl_3$),** δ: 19.6 (CH3, C-7), 118.3, 122.7, 127.8, 131.8, 133.4, 133.5 (CH, C-aromatic), 118.4, 130.0, 135.1, 135.5, 138.9, 139.8 (C, C-aromatic), 170.4 (C, C-14). **UPLC-MS: Rt** 2.13 (>98%%) **MS (ESI)$^+$:** 381.9 $[M+Na]^+$.

#### 2. Amine Compounds

##### 2.1 General Procedure for the synthesis of: arylamino derivatives (113, 115, 117, 119-120, 129, 131, 133-134, 144-145, 147-150)

A solution of the appropriate aryl benzaldehyde (**107,108,109,110,111,125,127,128**) (1eq) and differently substituted anilines (**2,18,84,96**) (1eq) in MeOH (2.5 mL/mmol) was refluxed overnight. The solvent was then removed under vacuum and the resulting residue was dissolved in acetic acid (3 mL/mmol). NaBH4 (2 eq) was then added in portions at 20°C and the mixture was then stirred at room temperature for 7 hours. The reaction mixture was added of water (4 mL/mmol) and ethyl acetate (4 mL/mmol). The pH of the aqueous layer was adjusted to 5 by the addition of 1M NaOH. The water layer was extracted with ethyl acetate (3x10 mL/mmol). The combined organic layers were dried over MgSO4 and concentrated under vacuum. The desired products were purified by flash column chromatography.

- **2-((4-(Trifluoromethyl)benzyl)amino)benzoic acid (113)**
  Obtained as white solid in 17% yield. **$^1$H-NMR ($CDCl_3$), δ:** 4.59 (s, 2H, H-8), 6.58 (m, 1H, H-aromatic), 6.67-6.70 (m, 1H, H-aromatic), 7.34-7.38 (m, 1H, H-aromatic), 7.50 (d, J= 7.9 Hz, 2H, H-aromatic), 7.63 (d, J= 7.9 Hz, 2H, H-aromatic), 8.05 (dd, J1= 8.0 Hz, J2= 1.5 Hz, 1H, H-aromatic), 8.20 (bs, 1H, NH), 11.22 (bs, 1H, OH). **$^{13}$C-NMR ($CDCl_3$),** δ 46.4 (CH2, C-8), 111.7, 115.6, 125.6, 125.7, 129.1, 129.4 (CH, C-aromatic), 109.1, 125.2, 127.0, 143.0, 151.3 (C, C-aromatic), 173.3 (C, C-15). **$^{19}$F-NMR ($CDCl_3$), δ:** -62.38 (3F). **UPLC-MS: Rt** 2.29 (>99%) **MS (ESI)$^+$:** 296.0 $[M+H]^+$.
- **2-((3,4-Dimethylbenzyl)amino)benzoic acid (115)**
  Obtained as white solid in 52% yield. **$^1$H-NMR ($CDCl_3$), δ:** 2.18 (d, J= 3.7 Hz, 6H, H-7,8), 4.33 (s, 2H, H-9), 6.52-6.55 (m, 1H, H-aromatic), 6.58 (d, J= 8.4 Hz, 1H, H-aromatic), 7.00-7.05 (m, 3H, H-aromatic), 7.23-7.27 (m, 1H, H-aromatic), 7.91 (dd, J1= 7.9 Hz, J2= 1.6 Hz, 1H, H-aromatic), 8.22 (bs, 1H, NH) 11.36 (bs, 1H, OH). **$^{13}$C-NMR ($CDCl_3$),** δ 19.4 (CH3), 19.8 (CH3), 46.7 (CH2, C-9), 111.9, 114.9, 124.4, 128.3, 129.9, 132.62, 135.4 (CH, C-aromatic), 108.8, 135.6, 136.0, 136.9, 151.7 (C, C-aromatic), 173.6 (C, C-16). **UPLC-MS: Rt** 2.34 (100%) **MS (ESI)$^+$:** 256.1 $[M+H]^+$.
- **2-((3-Chloro-4-methylbenzyl)amino)benzoic acid (117)**
  Obtained as white solid in 21% yield. **$^1$H-NMR ($CDCl_3$), δ:** 2.38 (s, 3H, H-7), 4.46 (s, 2H, H-8), 6.60-6.63 (m, 1H, H-aromatic), 6.65-6.68 (m, 1H, H-aromatic), 7.15-7.18 (m, 1H, H-aromatic), 7.21 (d, J= 7.8 Hz, 1H, H-aromatic), 7.34-7.37 (m, 2H, H-aromatic), 8.02 (dd, J1= 7.8 Hz, J2= 1.4 Hz, 1H, H-aromatic), 8.11 (m, 1H, H-aromatic), 10.77 (bs, 1H, NH), 12.73 (bs, 1H, OH). **$^{13}$C-NMR ($CDCl_3$),** δ 19.7 (CH3), 46.1 (CH2, C-8), 111.8, 115.3, 125.0, 127.4, 131.2, 132.6, 135.6 (CH, C-aromatic), 108.9, 134.6, 134.8, 138.1, 151.4 (C, C-aromatic), 172.8 (C-15) .**UPLC-MS: Rt** 2.37 (100%) **MS (ESI)$^+$:** 276.5 $[M+H]^+$.
- **2-((Quinolin-8-ylmethyl)amino)benzoic acid (119)**
  Obtained as yellow solid in 43% yield. **$^1$H-NMR (DMSO-$D_6$), δ:** 5.08 (s, 2H, H-10), 6.52-6.55 (m, 1H, H-aromatic), 6.71 (dd, J1= 8.5 Hz, J2= 0.6 Hz, 1H, H-aromatic), 7.24-7.29 (m, 1H, H-aromatic), 7.54-7.57 (m, 1H, H-aromatic), 7.60 (dd, J1= 8.2 Hz, J2= 4.1 Hz, 1H, H-aromatic), 7.68-7.70 (m, 1H, H-aromatic), 7.80 (dd, J1= 7.9 Hz, J2= 1.5 Hz, 1H, H-aromatic), 7.90 (dd, J1= 7.9 Hz, J2= 1.1 Hz, 1H, H-aromatic), 8.41 (dd, J1= 8.3 Hz, J2= 1.7 Hz, 1H, H-aromatic), 8.49 (bs, 1H, NH), 8.98 (dd, J1= 4.1 Hz, J2= 1.7 Hz, 1H, H-aromatic), 12.51 (bs, 1H, OH). **$^{13}$C-NMR (DMSO-$D_6$),** δ 42.6 (CH2, C-10), 112.0, 114.7, 122.0, 126.8, 127.6, 127.8, 132.2, 134.8, 136.9, 150.2 (CH, C-aromatic), 110.7, 128.4, 137.0, 146.2, 151.2 (C, C-aromatic), 170.4 (C, C- 17). **UPLC-MS: Rt** 1.95 (100%) **MS (ESI)$^+$:** 279.1 $[M+H]^+$.

- **2-((Quinoline-3-ylmethyl)amino)benzoic acid (120)**
  Obtained as yellow solid in 14% yield. **$^1$H-NMR (DMSO-D$_6$), δ:** 4.72 (s, 2H, H-10), 6.56-6.60 (m, 1H, H-aromatic), 6.72 (d, J= 8.06 Hz, 1H, H-aromatic), 7.27-7.31 (m, 1H, H-aromatic), 7.57-7.61 (m, 1H, H-aromatic), 7.71-7.75 (m, 1H, H-aromatic), 7.83 (d, J= 8.1 Hz, 1H, H-aromatic), 7.96 (d, J= 8.1 Hz, 1H, H-aromatic), 8.01 (d, J= 8.1 Hz, 1H, H-aromatic), 8.24 (s, 1H, H-aromatic), 8.45 (bs, 1H, NH), 8.93 (s, 1H, H-aromatic), 12.76 (bs, 1H, OH). **$^{13}$C-NMR (DMSO-D$_6$),** δ 112.1, 115.2, 127.2, 128.3, 129.1, 129.6, 132.2, 133.1, 134.9, 151.1 (CH, C-aromatic), 111.1, 127.9, 133.0, 147.3, 150.8 (C, C-aromatic), 170.4 (C, C- 17). **UPLC-MS: Rt** 1.62 (98%) **MS (ESI)$^+$:** 279.1 [M+H]$^+$.
- **Methyl 2-((3-chloro-4-methylbenzyl)amino)benzoate (129)**
  Obtained as grey solid in 26% yield. **$^1$H-NMR (CDCl$_3$), δ:** 2.26 (s, 3H, H-7), 3.79 (s, 3H, H-16), 4.31 (d, J= 5.7 Hz, 2H, H-8), 6.50-6.54 (m, 2H, H-aromatic), 6.81 (bs, 1H, NH), 7.06 (dd, J1= 7.8 Hz, J2= 1.1 Hz, 1H, H-aromatic), 7.09 (d, J= 7.8 Hz, 1H, H-aromatic), 7.24 (s, 1H, H-6), 7.84 (dd, J1= 7.9 Hz, J2= 1.6 Hz, 1H, H-aromatic), 8.08 (m, 1H, H-aromatic). **$^{13}$C-NMR (CDCl$_3$),** δ 19.7 (CH3, C-7), 51.5 (CH3, C-16), 46.2 (CH2, C-8), 111.6, 115.1, 125.2, 127.6, 131.1, 131.6, 134.68 (CH, C-aromatic), 110.3, 134.64, 134.7, 138.3, 150.7 (C, C-aromatic), 169.1 (C, C-15). **UPLC-MS: Rt** 2.69 (100%) **MS (ESI)$^+$:** 290.0 [M+H]$^+$.
- **Methyl 2-((pyridin-3-ylmethyl)amino)benzoate (131)**
  Obtained as yellow oil in 90% yield. **$^1$H-NMR (CDCl$_3$), δ:** 3.89 (s, 3H, H-14), 4.51 (d, J=5.7 Hz, 2H, H-6), 6.61 (d, J= 8.3 Hz, 1H, H-aromatic), 6.64-6.67 (m, 1H, H-aromatic), 7.28-7.30 (m, 1H, H-aromatic), 7.31-7.35 (m, 1H, H-aromatic), 7.69- 7.72 (m, 1H, H-aromatic), 7.96 (dd, J1= 8.0 Hz, J2= 1.6 Hz, 1H, H-aromatic), 8.23 (bs, 1H, NH), 8.54-8.56 (m, 1H, H-aromatic), 8.65 (s, 1H, H-2). **$^{13}$C-NMR (CDCl$_3$),** δ 51.6 (CH3, C-14), 44.5 (CH2, C-6), 111.5, 115.4, 123.6, 134.4, 134.7, 134.8, 148.6, 149.8 (CH, C-aromatic), 110.6, 131.7, 150.5 (C, C-aromatic), 169.1 (C, C-13). **UPLC-MS: Rt** 1.40 (96%) **MS (ESI)$^+$:** 243.1 [M+H]$^+$.
- **Methyl 2-((naphthalen-2-ylmethyl)amino)benzoate (133)**
  Obtained as yellow solid in 76% yield. **$^1$H-NMR (CDCl$_3$), δ:** 3.80 (s, 3H, H-19), 4.54 (d, J= 5.9 Hz, 2H, H-11), 6.50-6.54 (m, 1H, H-aromatic), 6.59 (d, J= 8.5 Hz, 1H, H-aromatic), 7.19-7.21 (m, 1H, H-aromatic), 7.35-7.37 (m, 1H, H-aromatic), 7.38-7.41 (m, 2H, H-aromatic), 7.70-7.72 (m, 2H, H-aromatic), 7.73-7.76 (m, 2H, H-aromatic), 7.86 (dd, J1= 8.2 Hz, J2= 1.6 Hz, 1H, H-aromatic), 8.20 (bs, 1H, NH).**$^{13}$C-NMR (CDCl$_3$),** δ 51.5 (CH3, C-19), 47.2 (CH2, C-11), 111.8, 114.9, 125.3, 125.5, 125.6, 126.1, 127.70, 127.79, 128.4, 131.6, 134.6 (CH, C-aromatic), 110.3, 132.7, 133.5, 136.4, 151.0 (C, C-aromatic), 169.1 (C, C-18). **UPLC-MS: Rt** 2.66 (100%) **MS (ESI)$^+$:** 292.0 [M+H]$^+$.
- **Methyl 2-((naphthalen-1-ylmethyl)amino)benzoate (134)**
  Obtained as white solid in 79% yield. **$^1$H-NMR (CDCl$_3$), δ:** 3.76 (s, 3H, H-19), 4.81 (d, J= 5.2 Hz, 2H, H-11), 6.53-6.57 (m, 1H, H-aromatic), 6.63 (d, J= 8.5 Hz, 1H, H-aromatic), 7.22-7.26 (m, 1H, H-aromatic), 7.32-7.35 (m, 1H, H-aromatic), 7.42-7.49 (m, 3H, H-aromatic), 7.71 (d, J= 8.3 Hz, 1H, H-aromatic), 7.81-7.83 (m, 1H, H-aromatic), 7.87 (dd,

J1= 8.0 Hz, J2= 1.7 Hz, 1H, H-aromatic), 7.97 (d, J= 8.3 Hz, 1H, H-aromatic), 8.09 (bs, 1H, NH).**$^{13}$C-NMR ($CDCl_3$),** δ 51.5 (CH3, C-19), 44.9 (CH2, C-11), 111.6, 114.9, 123.0, 125.1, 125.5, 125.7, 126.2, 127.9, 128.8, 131.6, 134.6 (CH, C-aromatic), 110.2, 131.3, 133.5, 133.8, 151.0 (C, C-aromatic), 169.0 (C, C-18). **UPLC-MS: Rt** 2.66 (100%) **MS (ESI)$^+$:** 292.0 $[M+H]^+$.

- **2-(*1H*-Tetrazol-5-*yl*)-*N*-(4-(trifluoromethyl)benzyl)aniline (144)**
  Obtained as white solid in 32% yield. **$^1$H-NMR (DMSO-$D_6$), δ:** 4.70 (s, 2H, H-8), 6.73-6.78 (m, 2H, H-aromatic), 7.28-7.31 (m, 1H, H-aromatic), 7.59 (d, J= 8.1 Hz, 2H, H-aromatic), 7.71 (d, J= 8.1 Hz, 2H, H-aromatic), 7.82-7.84 (m, 1H, H-aromatic). **$^{13}$C-NMR (DMSO-$D_6$),** δ 46.1 (CH2, C-8), 112.4, 116.3, 125.8, 128.1, 129.0, 132.7 (CH, C-aromatic), 106.3, 123.7, 127.9, 145.1, 146.7 (C-aromatic). **UPLC-MS: Rt** 1.95 (>99%) **MS (ESI)$^+$:** 320.1 $[M+H]^+$.
- ***N*-(3-Chloro-4-methylbenzyl)-2-(*1H*-tetrazol-5-*yl*)aniline (145)**
  Obtained as white solid in 25% yield. **$^1$H-NMR (DMSO-$D_6$), δ:** 2.30 (s, 3H, H-7), 4.54 (s, 2H, H-8), 6.74-6.78 (m, 2H, H-aromatic), 7.25 (d, J= 7.5 Hz, 1H, H-aromatic), 7.30-7.32 (m, 2H, H-aromatic), 7.41 (s, 1H, H-6), 7.81 (d, J= 7.5 Hz, 1H, H-aromatic). **$^{13}$C-NMR (DMSO-$D_6$),** δ 19.6 (CH3, C-7), 45.8 (CH2, C-8), 112.4, 116.2, 126.2, 127.7, 128.9, 131.8, 132.7 (CH, C-aromatic), 133.0, 133.7, 134.3, 138.7, 139.8 (C, C-aromatic), 149.7 (C, C-15). **UPLC-MS: Rt** 2.33 (100%) **MS (ESI)$^+$:** 300.0 $[M+H]^+$.
- ***N*-(Quinolin-8-ylmethyl)-2-(*1H*-tetrazol-5-*yl*)aniline (147)**
  Obtained as yellow solid in 12% yield. **$^1$H-NMR (DMSO-$D_6$), δ:** 5.18 (s, 2H, H-10), 6.71-6.74 (m, 1H, H-aromatic), 6.75-6.78 (m, 1H, H-aromatic), 7.27-7.30 (m, 1H, H-aromatic), 7.55-7.58 (m, 1H, H-aromatic), 7.59-7.62 (m, 1H, H-aromatic), 7.73-7.75 (m, 1H, H-aromatic), 7.80-8.82 (m, 1H, H-aromatic), 7.90-7.92 (m, 1H, H-aromatic), 8.40-8.42 (m, 1H, H-aromatic), 8.98-9.00 (m, 1H, H-aromatic). **$^{13}$C-NMR (DMSO-$D_6$),** δ 43.0 (CH2, C-10), 112.3, 115.9, 122.0, 126.8, 127.7, 127.9, 128.9, 132.8, 136.9, 150.3 (CH, C-aromatic), 128.4, 146.2, 147.1 (C-aromatic). **UPLC-MS: Rt** 1.68 (>96%) **MS (ESI)$^+$:** 303.3 $[M+H]^+$.
- ***N*-(Quinolin-3-ylmethyl)-2-(*1H*-tetrazol-5-*yl*)aniline (148)**
  Obtained as yellow solid in 11% yield. **$^1$H-NMR (DMSO-$D_6$), δ:** 4.82 (s, 2H, H-10), 6.5-6.78 (m, 1H, H-aromatic), 6.86-6.88 (m, 1H, H-aromatic), 7.29-31 (m, 1H, H-aromatic), 7.58-7.61 (m, 1H, H-aromatic), 7.72-7.75 (m, 1H, H-aromatic), 7.83-7.85 (m, 1H, H-aromatic), 7.92-7.95 (m, 1H, H-aromatic), 8.01-8.03 (m, 1H, H-aromatic), 8.29 (s, 1H, H-1), 8.97 (s, 1H, H-3). **$^{13}$C-NMR (DMSO-$D_6$),** δ 49.0 (CH2, C-10), 112.4, 116.3, 127.2, 128.3, 129.0, 129.1, 129.6, 132.6, 133.8, 151.2 (CH, C-aromatic), 106.7, 127.9, 132.9, 146.6, 147.3 (C-aromatic). **UPLC-MS: Rt** 1.59 (>96%) **MS (ESI)$^+$:** 303.1 $[M+H]^+$.
- ***N*-(Naphthalen-2-ylmethyl)-2-(*1H*-tetrazol-5-*yl*)aniline (149)**
  Obtained as white solid in 75% yield. **$^1$H-NMR (DMSO-$D_6$), δ:** 4.74 (s, 2H, H-11), 6.74-6.77 (m, 1H, H-aromatic), 6.85 (d, J=8.3 Hz, 1H, H-aromatic), 7.28-7.31 (m, 1H, H-aromatic), 7.47-7.52 (m, 2H, H-aromatic), 7.54 (d, J= 8.3 Hz, 1H, H-aromatic), 7.81-7.87 (m, 2H, H-aromatic), 7.90-7.92 (m, 3H, H-aromatic).**$^{13}$C-NMR (DMSO-$D_6$),** δ 46.9

(CH2, C-11), 112.5, 116.1, 125.8, 126.1, 126.1, 126.7, 128.0, 128.0, 128.6, 128.9, 132.70 (CH, C-aromatic), 132.72, 133.4, 137.8, 147.0 (C, C-aromatic). **UPLC-MS: Rt** 1.97 (>98%) **MS (ESI)$^-$:** 300.2 $[M-H]^-$.

- **3-(2-((3-Chloro-4-methylbenzyl)amino)phenyl)-1,2,4-oxadiazol-5(4H)-one (150)**
  Obtained as off-white solid in 40% yield. **$^1$H-NMR (DMSO-D$_6$), δ:** 2.37 (s, 3H, H-7), 4.50 (s, 2H, H-8), 6.70-6.73 (m, 2H, H-aromatic), 7.08 (bs, 1H, NH), 7.21 (d, J= 7.2 Hz, 1H, H-aromatic), 7.30-7.33 (m, 2H, H-aromatic), 7.39 (s, 1H, H-2), 7.55 (d, J= 7.2 Hz, 1H, H-aromatic), 12.79 (bs, 1H, NH).**$^{13}$C-NMR (DMSO-D$_6$),** δ 19.6 (CH3, C-7), 45.7 (CH2, C-8), 112.3, 116.1, 126.2, 127.7, 129.0, 131.8, 133.3 (CH, C-aromatic), 105.5, 133.7, 134.3, 139.5, 146.9, 158.7 (C, C-aromatic), 159.5 (C, C-15). **UPLC-MS: Rt** 2.35 (>99%) **MS (ESI)$^-$ :** 314.1 $[M-H]^-$.

#### 2.2 General Procedure for the synthesis of: methyl((aryl)amino)benzoates (121,123)

The appropriate (arylamino)benzoic acid (**113, 115**) (1 eq), TBTU (1 eq) and DIPEA (2 eq) was stirred in anhydrous DMF (1 mL/mmol eq), and the resulting mixture was stirred at room temperature for 30 min, under N2 atmosphere. CH3OH (1 eq) was then injected into the reaction mixture via syringe and stirring was continued at room temperature until completion of the reaction. The reaction mixture was diluted in DCM (10 mL/mmol eq) and the resulting mixture was extracted with sat.aq NH4Cl (1x10 mL/mmol eq) and sat.aq NaHCO3 (1x10 mL/mmol eq). The organic layer was dried overv MgSO4 and the solvent was evaporated under vacuum. The desired products were purified by flash column chromatography or recrystallisation.

- **Methyl 2-((4-(trifluoromethyl)benzyl)amino)benzoate (121)**
  Obtained as yellow solid in 44% yield. **$^1$H-NMR (CDCl$_3$), δ:** 3.81 (s, 3H, H-16), 4.45 (d, J= 5.8 Hz, 2H, H-8), 6.45-6.48 (m, 1H, H-aromatic), 6.54-6.57 (m, 1H, H-aromatic), 7.20-7.24 (m, 1H, H-aromatic), 7.39 (d, J= 8.0 Hz, 2H, H-aromatic), 7.52 (d, J= 8.0 Hz, 2H, H-aromatic), 7.85-7.88 (m, 1H, H-aromatic), 8.18 (bs, 1H, NH).**$^{13}$C-NMR (CDCl$_3$),** δ 51.6 (CH3, C-16), 46.5 (CH2, C-8), 111.5, 115.3, 125.6, 125.7, 131.7, 134.6 (CH, C-aromatic), 110.5, 125.6, 129.3, 143.1, 150.6 (C, C-aromatic), 169.1 (C, C-15). **$^{19}$F-NMR (CDCl$_3$), δ:** -62.43 (s, 3F). **UPLC-MS: Rt** 2.60 (>98%) **MS (ESI)$^+$:** 310.1 $[M+H]^+$.
- **Methyl 2-((3,4-dimethylbenzyl)amino)benzoate (123)**
  Obtained as yellow solid in 13% yield. **$^1$H-NMR (CDCl$_3$), δ:** 2.17 (d, J= 3.7 Hz, 6H, H-7,8), 3.77 (s, 3H, H-17), 4.30 (d, J=5.4 Hz, 2H, H-9), 6.49-6.52 (m, 1H, H-aromatic), 6.56-6.59 (m, 1H, H-aromatic), 7.00-7.02 (m, 2H, H-aromatic), 7.04 (s, 1H, H-aromatic), 7.20-7.23 (m, 1H, H-aromatic), 7.83 (dd, J1= 8.0 Hz, J2= 1.6 Hz, 1H, H-aromatic), 8.01 (bs, 1H, NH).**$^{13}$C-NMR (CDCl$_3$),** δ 19.4 (CH3), 19.8 (CH3), 51.4 (CH3, C-17), 46.83(CH2, C-9), 111.6, 114.7, 124.5, 128.5, 129.9, 131.5, 134.6 (CH, C-aromatic), 110.1, 135.4, 136.2, 136.8, 151.0 (C, C-aromatic), 169.1 (C-16). **UPLC-MS: Rt** 2.68(>99%) **MS (ESI)$^+$:** 270.1 $[M+H]^+$.

**2.3 Synthesis of: methylmethyl 2-(phenethylamino)benzoate (137)**

To a solution of (2-bromoethyl) benzene (0.150 g, 0.81 mmol) in MeCN (5 mL) was added methyl 2- aminobenzoate (0.122 g, 0.81 mmol) under nitrogen atmosphere. The resulting solution was stirred at reflux for 48 h. The formed salt was filtered off and the filtrate was concentrated under vacuum. The residue was dissolved in ethyl acetate (15 mL) and washed sequentially with water (10 mL) and brine (10 mL) and dried over sodium sulfate. The solvent was evaporated under reduced pressure and the residue was purified by flash column chromatography (Biotage Isolera One automated flash column chromatography, cartridge: ZIP KP 10g, *n*hexane-EtOAc 100:0 increasing to *n*-hexane-EtOAc 80:20 in 15CV. **137** was obtained as colourless oil in 5% yield. **$^1$H-NMR ($CDCl_3$), δ:** 2.90 (t, J= 7.5 Hz, 2H, H-7), 3.37 (t, J= 7.5 Hz, 2H, H-8), 3.76 (s, 3H, H-15), 6.49- 6.52 (m, 1H, H-aromatic), 6.62-6.64 (m, 1H, H-aromatic), 7.12-7.15 (m, 1H, H-aromatic), 7.18-7.20 (m, 2H, H-aromatic), 7.23-7.29 (m, 3H, H-aromatic), 7.69 (bs, 1H, NH), 7.81-7.83 (m, 1H, H-aromatic).**$^{13}$C-NMR ($CDCl_3$),** δ 51.4 (CH3, C-15), 32.9, 44.5 (CH2, C-7,8), 111.1, 114.5, 128.6, 127.7, 131.5, 131.7, 134.6 (CH, C-aromatic), 110.0, 139.2, 150.9 (C, C-aromatic), 168.9 (C, C-14). **UPLC-MS: Rt** 2.58(>97%) **MS (ESI)$^+$:** 256.1 $[M+H]^+$.

**2.4 Synthesis of methyl 2-((4-trifluoromethyl)phenethylamino)benzoate (138)**

A mixture of methyl 2-aminobenzoate (0.090 g, 0.59 mmol), 1-(2-bromoethyl)-4-(trifluoromethyl)benzene (0.150 g, 0.59 mmol), and K2CO3 (0.081 g, 0.59 mmol) in MeCN (2 mL) was stirred overnight under reflux. The mixture was extracted between ethylacetate (3 x 10 mL) and water (20 mL) and the organic layer was dried over sodium sulfate. The solvent was removed under vacuum and the residue was purified by flash column chromatography (Biotage Isolera One automated flash column chromatography, cartridge: ZIP KP SIL 30g, *n*-hexane-DCM 100:0 increasing to *n*-hexane-DCM 40:60 in 15CV). **138** was obtained as colourless oil in 26% yield. **$^1$H-NMR ($CDCl_3$), δ:** 3.06 (t, J= 7.2 Hz, 2H, H-8), 3.49-3.53 (m, 2H, H-9), 3.86 (s, 3H, H-16), 6.61-6.64 (m, 1H, H-aromatic), 6.70-5.72 (m, 1H, H-aromatic), 7.37-7.40 (m, 3H, H-aromatic), 7.59 (d, J= 8.0 Hz, 2H, H-aromatic), 7.80 (bs, 1H, NH), 7.93 (dd, J1= 8.0 Hz, J2= 1.6 Hz, 1H, H-aromatic).**$^{13}$C-NMR ($CDCl_3$),** δ 35.4, 44.0 (CH2, C-8,9) 51.4 (CH3, C-16), 111.0, 114.8, 125.4, 125.5, 129.1, 131.78 (CH, C-aromatic), 125.4 (d, J= 37.3 Hz, C), 110.2, 134.6, 143.4, 150.7 (C-aromatic), 168.9 (C, C- 15). **$^{19}$F-NMR ($CDCl_3$), δ:** -62.4 (s, 3F). **UPLC-MS: Rt** 2.68(100%) **MS (ESI)$^+$:** 324.2 $[M+H]^+$.

**2.5 General Procedure for the synthesis of (aryl)amino)benzoic acids (139, 141-143)**

To a solution of the appropriate methyl-(arylamino) benzoate (**131,133,134,136,163)** (1 eq) in THF (7 mL/mmol), H2O (3.5 mL/mmol), MeOH (3.5 mL/mmol) was added LiOH (3 eq). The reaction was stirred ad 70°C overnight. After the reaction was allowed to cool to room temperature, the reaction mixture was acidified with 2M HCl. The product was extracted with ethyl acetate (3x10 mL/mmol). The organic layer was dried over MgSO4 and evaporated under reduced pressure. The product was purified by flash column chromatography or recrystallisation.

- **2-((Pyridin-3-ylmethyl)amino)benzoic acid (139)**

Obtained as white solid in 28% yield. **$^1$H-NMR (DMSO-D$_6$), δ:** 4.52 (S, 2H, H-6), 6.56-6.60 (m, 1H, H-aromatic), 6.69 (d, J= 8.4 Hz, 1H, H- aromatic), 7.29-7.30 (m, 1H, H-aromatic), 7.34 (dd, J1= 7.7 Hz, J2= 4.7 Hz, 1H, H-4), 7.73 (d, J= 7.7 Hz, 1H, H-5), 7.81 (dd, J1= 8.0 Hz, J2= 1.4 Hz, 1H, H-aromatic), 8.30 (bs, 1H, NH), 7.47 (d, J= 4.7 Hz, 1H, H-3), 8.59 (s, 1H, H-2), 12.76 (bs, 1H, OH).**$^{13}$C-NMR (DMSO-D$_6$),** δ 43.7 (CH2, C-6), 112.1, 115.2, 124.1, 132.2, 134.8, 135.3, 148.6, 149.1 (CH, C-aromatic), 111.1, 135.4, 150.8 (C, C-aromatic), 170.4 (C, C-13). **UPLC-MS: Rt** 1.20 (>98%) **MS (ESI)$^+$:** 229.1 [M+H]$^+$.

- **2-((Naphthalen-2-ylmethyl)amino)benzoic acid (141)**
  Obtained as white solid in 89% yield. **$^1$H-NMR (DMSO-D$_6$), δ:** 4.64 (S, 2H, H-11), 6.54-6.58 (m, 1H, H-aromatic), 6.72 (d, J= 8.2 Hz, 1H, H-aromatic), 7.27-7.30 (m, 1H, H-aromatic), 7.47-7.53 (m, 3H, H-aromatic), 7.82 (dd, J1= 7.9 Hz, J2= 1.5 Hz, 1H, H-aromatic), 7.85-7.88 (m, 2H, H-aromatic), 7.89-7.92 (m, 2H, H-aromatic), 8.41 (bs, 1H, NH), 12.68 (bs, 1H, OH).**$^{13}$C-NMR (DMSO-D$_6$),** δ 45.5 (CH$_2$, C-11), 112.2, 115.0, 125.6, 126.0, 126.1, 126.7, 128.01, 128.04, 128.6, 132.1, 134.8 (CH, C-aromatic), 110.8, 132.7, 133.4, 137.5, 151.1 (C, C-aromatic), 170.5 (C, C- 18). **UPLC-MS: Rt** 2.32 (99%) **MS (ESI)$^+$:** 278.0 [M+H]$^+$.
- **2-((|Naphthalen-1-ylmethyl)amino)benzoic acid (142)**
  Obtained as white solid in 41% yield. **$^1$H-NMR (DMSO-D$_6$), δ:** 4.91 (s, 2H, H-11), 6.51-6.61 (m, 1H, H-aromatic), 6.78 (dd, J1= 8.7 Hz, J2= 0.4 Hz, 1H, H-aromatic), 7.32-7.35 (m, 1H, H-aromatic), 7.46-7.50 (m, 2H, H-aromatic), 7.55-7.61 (m, 2H, H-aromatic), 7.83 (dd, J1= 7.9 Hz, J2= 1.6 Hz, 1H, H-aromatic), 7.87 (dd, J1= 7.4 Hz, J2= 1.3 Hz, 1H, Haromatic), 7.97-8.00 (m, 1H, H-aromatic), 8.11-8.12 (m, 1H, H-aromatic), 8.26 (bs, 1H, NH), 12.63 (bs, 1H, OH).**$^{13}$C-NMR (DMSO-D$_6$),** δ 44.4 (CH2, C-11), 112.1, 115.0, 123.8, 125.3, 126.0, 126.3, 126.7, 128.1, 129.0, 132.1, 134.9 (CH, C-aromatic), 110.7, 131.3, 133.9, 134.6 (C, C-aromatic), 170.4 (C, C- 18). **UPLC-MS: Rt** 2.33 (99%) **MS (ESI)$^+$:** 278.1 [M+H]$^+$.
- **2-((4-(Trifluoromethyl)phenethyl)amino)benzoic acid (143)**
  Obtained as pale yellow solid in 16% yield. **$^1$H-NMR (CDCl$_3$), δ:** 3.07 (t, J= 7.06 Hz, 2H, H-alkyl), 3.54 (t, J= 7.06 Hz, 2H, H-alkyl), 6.66-6.69 (m, 1H, H-aromatic), 6.74-6.76 (m, 1H, H-aromatic), 7.41 (d, J= 7.7 Hz, 2H, H-aromatic), 7.44-7.46 (m, 1H, H- aromatic), 7.62 (d, J= 7.7 Hz, 2H, H-aromatic), 7.72 (bs, 1H, NH), 8.01 (dd, J1= 7.9 Hz, J2= 1.3 Hz, 1H, H-aromatic), 11.24 (bs, 1H, OH). **$^{13}$C-NMR (CDCl$_3$),** δ 35.3, 43.9 (2xCH2, C-8,9), 111.2, 115.0, 125.5, 129.1, 132.7, 135.7 (CH, C-aromatic), 128.8 (d, J= 32.42 Hz, C) 102.5, 108.7, 143.1, 151.3 (C, C-aromatic), 173.2 (C, C- 15). **$^{19}$F-NMR (CDCl$_3$), δ:** -62.3 (s, 3F). **UPLC-MS: Rt** 2.37 (100%) **MS (ESI)$^+$:** 310.1 [M+H]$^+$.
- **2-(((5-Chloro-2-methylphenyl)amino)methyl)benzoic acid (156)**
  Obtained as white solid in 53% yield. **$^1$H-NMR (CDCl$_3$), δ:** 2.14 (s, 3H, H-7), 4.68 (s, 2H, H-8), 5.94 (bs, 1H, NH), 6.23 (d, J= 2.0 Hz, 1H, H-2), 6.47 (dd, J1= 7.8 Hz, J2= 2.0 Hz, 1H, H-4), 6.97 (d, J= 7.8 Hz, 1H, H-5), 7.33-7.37 (m, 1H, H-aromatic), 7.42 (d, J= 7.3 Hz, 1H, H-aromatic), 7.49-7.52 (m, 1H, H-aromatic), 7.89 (dd, J1= 7.7 Hz, J2= 1.2 Hz, 1H,

H-aromatic), 13.09 (bs, 1H, OH).**$^{13}$C-NMR ($CDCl_3$),** 17.6 (CH3, C-7), 45.2 (CH2, C-8), 109.0, 115.3, 127.2, 128.0, 131.0, 131.4, 132.4 (CH, C-aromatic), 121.1, 130.1, 131.6, 141.3, 147.9 (C, C-aromatic), 169.1 (C, C- 15). **UPLC-MS: Rt** 2.19 (100%) **MS (ESI)$^+$:** 276.0 $[M+H]^+$.

#### 3. Methylene Compounds

##### 3.1 Synthesis of Methyl 2-(4-(trifluoromethyl)benzyl)benzoate (169)

A round bottom flask was charged with Pd(OAc)2 (0.001 eq), PPh3 (0.002 eq), (2-(methoxycarbonyl)phenyl)boronic acid (1.5 eq), and K3PO4 (2 eq), under N2 atmosphere. Then the 1-(bromomethyl)-4-(trifluoromethyl)benzene (0.150 g, 0.67 mmol, 1 eq) and toluene (4.5 mL/mmol) were added. The reaction was stirred at 80°C for 48 hours. The solution was diluted with diethyl ether (15 mL/mmol) and washed with NaOH 1M solution (10 mL/mmol), brine (10 mL/mmol) and then dried over MgSO4. The residue obtained was purified by flash column chromatography (Biotage Isolera One automated flash column chromatography, cartridge: SNAP KP SIL 25g, *n*-hexane-DCM 100:0 increasing to *n*-hexane-DCM 60:40 in 15CV. **169** was obtained as colourless oil in 37% yield. **$^1$H-NMR ($CDCl_3$), δ:** 3.84 (s, 3H, H-16), 4.47 (s, 2H, H-8), 7.24-7.26 (m, 1H, H-aromatic), 7.27 (d, J= 8.2 Hz, 2H, H-aromatic), 7.33-7.37 (m, 1H, H-aromatic), 7.47-7.51 (m, 1H, H-aromatic), 7.53 (d, J= 8.2 Hz, 2H, H-aromatic), 7.97 (dd, J1= 7.8 Hz, J2= 1.4 Hz, 1H, H-aromatic).**$^{13}$C-NMR ($CDCl_3$), δ:** 52.0 (CH3, C-16), 39.5 (CH2, C-8), 125.4, 126.7, 128.1, 128.8, 129.0, 129.7 (CH, C-aromatic), 125.5, 128.3, 128.7, 141.1, 145.1 (C, C-aromatic), 167.7 (C, C-15). **UPLC-MS: Rt** 2.54 (100%) **MS (ESI)$^+$:** 295.9 $[M+H]^+$.

##### 3.2 Synthesis of Methyl 2-(4-(trifluoromethyl)benzyl)benzoate (173)

To a solution of methyl 2-(3-chloro-4-methylbenzyl)benzoate (0.096 g, 0.35 mmol, 1 eq) in THF (7 mL/mmol), H2O (3.5 mL/mmol), MeOH (3.5 mL/mmol) was added LiOH (3 eq). The reaction was stirred ad 70°C overnight. After the reaction was allowed to cool to room temperature, the reaction mixture was acidified with 2M HCl. The product was extracted with ethyl acetate (3x10 mL/mmol). The organic layer was dried over MgSO4 and evaporated under reduced pressure. The product was purified by flash column chromatography (Biotage Isolera One automated flash column chromatography, cartridge: ZIP KP SIL 10g, *n*-hexane-EtOAc 100:0 increasing to *n*-hexane-EtOAc 40:60 in 14CV). **173** was obtained as a white solid in 67% yield. **$^1$H-NMR ($CDCl_3$), δ:** 2.24 (s, 3H, H-7), 4.31 (s, 2H, H-8), 6.88 (dd, J1= 7.7 Hz, J2= 1.5 Hz, 1H, H-4), 7.03 (d, J= 7.7 Hz, 1H, H-3), 7.06 (d, J= 1,5 Hz, 1H, H-6), 7.14-7.16 (m, 1H, H-aromatic), 7.25-7.28 (m, 1H, H-aromatic), 7.40-7.44 (m, 1H, H-aromatic), 8.01 (dd, J1= 7.8 Hz, J2= 1.3 Hz, 1H, H-aromatic), 11.36 (bs, 1H, OH).**$^{13}$C-NMR ($CDCl_3$), δ:** 19.6 (CH3, C-7), 38.9 (CH2, C-8), 126.6, 127.3, 129.4, 130.8, 131.7, 131.8, 133.1 (CH, C-aromatic), 128.2, 133.5, 134.2, 140.0, 142.8 (C, C-aromatic), 172.1 (C, C-15). **UPLC-MS: Rt** 2.31 (>98%) **MS (ESI)$^-$:** 259.1 $[M-H]^-$.

#### 4. Amide Compounds

##### 4.1 General Procedure for the synthesis of alkyl 2-arylamido benzoates (179, 182)

A solution of differently substituted benzoyl chloride (1 eq) in THF (1.5 mL/mmol) was slowly added at 0°C to a solution of the appropriate alkyl aminobenzoate (1.1 eq) and NEt3 (1.1 eq) in THF (6.5 mL/mmol). The reaction was stirred at room temperature for 6 hours. The formed salt was removed by filtration and washed with THF. The solvent was removed under reduced pressure. The crude product was purified by flash column chromatography or recrystallization.

- **Methyl 2-(4-(trifluoromethyl)benzamido)benzoate (179)**
  Obtained as white solid in 37% yield. **$^1$H-NMR (DMSO-D$_6$), δ:** 3.84 (s, 3H, H-16), 7.28-7.31 (m, 1H, H-aromatic), 7.69-7.33 (m, 1H, H-aromatic), 7.99 (d, J= 8.2 Hz, 2H, H-aromatic), 8.01-8.03 (m, 1H, H-aromatic), 8.16 (d, J= 8.2 Hz, 2H, H-aromatic), 8.46 (dd, J1= 7.3 Hz, J2= 1.0 Hz, 1H, H-aromatic), 11.59 (bs, 1H, NH).**$^{13}$C-NMR (DMSO-D$_6$), δ**: 53.1 (CH3, C-16), 121., 125.3, 126.5, 128.5, 131.1, 134.7 (CH, C-aromatic), 118.6, 132.4, 138.6, 139.3, 140.8, (C, C-aromatic), 164.1, 168.3 (C, C-8,15). **$^{19}$F-NMR (DMSO-D$_6$), δ:** -61.41 (s, 3F). **UPLC-MS: Rt** 2.52 (100%) **MS (ESI)$^+$:** 324.2 [M+H]$^+$.
- **Methyl 2-(3-chloro-2-methylbenzamido)benzoate (182)**
  Obtained as white solid in 28% yield. **$^1$H-NMR (CDCl$_3$), δ:** 2.57 (s, 3H, H-7), 3.93 (s, 3H, H-16), 7.16-7.20 (m, 1H, H-aromatic), 7.24-7.28 (m, 1H, H-aromatic), 7.48-7.51 (m, 2H, H-aromatic), 7.63-7.66 (m, 1H, H-aromatic), 8.11 (dd, J1= 6.3 Hz, J2= 1.5Hz, 1H, H-aromatic), 8.90-8.93 (m, 1H, H-aromatic), 11.45 (bs, 1H, NH).**$^{13}$C-NMR (CDCl$_3$), δ**: 17.1 (CH3), 52.4 (CH3), 123.0, 125.3, 126.0, 127.0, 131.0, 131.1, 134.3 (CH, C-aromatic), 115.3, 134.5, 136.1, 139.0, 141.4 (C, C-aromatic), 167.7, 168.7 (C, C-8, 15). **UPLC-MS: Rt** 2.50 (>98%) **MS (ESI)$^+$:** 304.1 [M+H]$^+$.

##### 4.2 Synthesis of 2-(3-chloro-4-methylbenzamido)benzoic acid (186)

A mixture of anthranilic acid (0.020 g, 0.14 mmol), NEt3 (0.018 g, 0.18 mmol) and CH2Cl2 (4 mL) was stirred at room temperature. 3-chloro-4-methyl benzoyl chloride (0.030 g, 0.15 mmol) was then added slowly at 0oC. The reaction was stirred overnight. The reaction mixture was washed with water and extracted with CH2Cl2 (20 mL x 2). The organic layers were combined, dried over sodium sulfate, and concentrated under reduced pressure. The product was purified by recrystallisation from EtOH/Water. **186** was obtained as white solid in 52% yield. **$^1$H-NMR (DMSO-D$_6$), δ:** 2.48 (s, 3H, H-7), 7.19-7.22 (m, 1H, H-aromatic), 7.40 (d, J= 7.9 Hz, 1H, H-13), 7.68-7.72 (m, 1H, H-aromatic), 7.82 (dd, J1= 7.9 Hz, J2= 1.6 Hz, 1H, H-14), 7.83 (d, J= 1.6 Hz, 1H, H- 10), 8.20 (dd, J1= 8.0 Hz, J2= 1.4 Hz, 1H, H-aromatic), 8.94-8.96 (m, 1H, H-aromatic), 11.85 (bs, 1H, NH).**$^{13}$C-NMR (DMSO-D$_6$), δ**: 20.2 (CH3, C-15), 120.5, 122.9, 125.3, 128.4, 131.2, 131.9, 135.9 (CH, C-aromatic), 113.8, 134.0, 135.1, 140.5, 142.3 (C, C-aromatic), 164.5, 171.2 (C, C-carbonyl). **UPLC-MS: Rt** 2.25 (99%) **MS (ESI)$^+$:** 290.1 [M+H]$^+$.
